# Large-Scale Mutagenesis Identifies Sequence Changes That Bias Tau Toward CTE-like Aggregation

**DOI:** 10.64898/2026.10.08.757173

**Authors:** Chad M. Dashnaw, Simran Rastogi, Nabil Morgan, Isaam Memon, Klaudia Jaczynska, Binh An Nguyen, Lorena Saelices, Charles L. White, Sarah Shahmoradian, Lukasz A. Joachimiak

## Abstract

Tau amyloid fibrils adopt disease-specific structures across tauopathies, yet the sequence determinants that favor one pathological fold over another remain poorly defined. Here we combine computational saturation mutagenesis and energetic profiling, in vitro aggregation, cryo-EM and cellular seeding assays to identify sequence changes that alter tau fold compatibility. Across six tauopathy fibril structures, approximately 12,000 substitutions revealed that mutational sensitivity is concentrated within interaction networks formed by amyloidogenic motifs rather than uniformly distributed across the fibril core. Comparison of Alzheimer’s disease (AD) and chronic traumatic encephalopathy (CTE) folds identified residues within the 335-363 region that differentially influence fold compatibility. The S341L/S352I double mutant accelerated aggregation and remodeled packing within the PAM4-containing amyloidogenic region, producing an alternative amyloid architecture in a minimal tau fragment. S341L/S352I biosensors preferentially responded to CTE-derived over Alzheimer’s disease-derived seeds, while aggregates generated from recombinant S341L/S352I tau exhibited CTE-like cellular templating behavior. Although these assemblies are not established to reproduce the atomic CTE fold, the results demonstrate that targeted sequence changes can bias tau aggregation and propagation toward a CTE-compatible state. This framework provides a strategy for defining sequence–structure relationships in polymorphic amyloids and developing conformation-selective models and biosensors.

## Introduction

The deposition of the microtubule-associated protein tau into β-sheet-rich amyloid assemblies underlies a large family of neurodegenerative disorders collectively known as tauopathies.^1–5^ Recent advances in cryo-electron microscopy (cryo-EM) have transformed our understanding of these diseases by revealing that tau adopts structurally distinct fibril conformations in different disorders.^1–6^ These findings support the concept that protein structure encodes disease phenotype and suggest that disease-specific tau conformations may serve as powerful diagnostic and therapeutic targets.^1–3^ Structural comparisons of ex vivo tau fibrils have uncovered a remarkable diversity of amyloid folds, ranging from highly divergent conformations to closely related structures that differ only within localized regions of the fibril core.^7^ The structurally characterized folds considered here include Alzheimer’s disease (AD), chronic traumatic encephalopathy (CTE), corticobasal degeneration (CBD), argyrophilic grain disease (AGD), progressive supranuclear palsy (PSP) and globular glial tauopathy (GGT).^5,7–9^ Computational analyses, including energetic profiling and alanine scanning, are beginning to reveal the sequence determinants and interaction networks that stabilize distinct tau folds.^10–12^ Nevertheless, discriminating between closely related conformations, such as AD and CTE fibrils, both of which adopt C-shaped architectures, remains challenging.^5,8,13,14^ Experimentally recreating disease-associated tau conformations in vitro or in cellular systems is similarly difficult, limiting our ability to determine how structure influences disease progression, propagation and toxicity.^13–17^

At the same time, advances in deep learning-based modeling and protein sequence design, including AlphaFold and ProteinMPNN^18–20^, now enable exploration of protein structure and sequence space and provide new opportunities to engineer amyloid assembly pathways. While these approaches have revolutionized the design of globular proteins, their application to intrinsically disordered proteins and amyloid structures remains challenging because aggregation emerges from dynamic conformational ensembles and weak intermolecular interactions rather than stable folded states.^21–25^ Moreover, because the cross-β architecture of amyloids can accommodate many distinct sequences, the sequence space compatible with amyloid formation is exceptionally large. ^20,26^ Generative approaches can therefore produce highly divergent, multi-mutation designs whose experimental behavior is difficult to attribute to specific residues or interactions. For this reason, we focused on saturation mutagenesis on fixed backbones combined with energetic profiling using established energy functions across a set of tauopathy amyloid structures as a more mechanistically interpretable and tractable strategy for defining the local sequence rules governing tau aggregation.^10–12^ Unlike generative design approaches that learn statistical relationships, saturation mutagenesis with an explicit energy function assigns each substitution a defined energetic cost, enabling direct identification of sequence tolerance, energetic hotspots and residues that are essential, permissive or disruptive for aggregation and seeding.

This approach is particularly valuable for amyloid systems because it directly reveals how single substitutions alter the energetics in cross-β scaffolds, relationships that can be difficult to extract from deep-learning predictions alone. For example, residues such as cysteines can regulate aggregation through transient intermolecular interactions, redox-sensitive chemistry or conformational dynamics that are difficult to predict from static structures alone.^27–30^ By systematically interrogating each residue within tauopathy fibril cores, saturation mutagenesis combined with an energy function provides an experimentally grounded map of the local energetic landscape governing aggregation and propagation. Importantly, this framework also allows direct comparison of mutational sensitivity across different disease-associated folds, revealing how distinct tau strains differentially depend on specific residues or interaction networks. Importantly, large mutational datasets, particularly when supported by experimental validation, will be essential for developing new models tailored to specific protein properties, such as structural stability, folding or aggregation propensity^11,15,31–33^. In contrast to ProteinMPNN-guided sampling, which prioritizes identification of globally compatible sequence solutions, saturation mutagenesis enables mechanistic dissection of the individual sequence determinants that control strain compatibility and seeded aggregation behavior.

Here, we integrate computational saturation mutagenesis with biochemical aggregation assays, cryo-EM and cellular seeding/propagation to define sequence features that control tau amyloid polymorphism. Our saturation mutagenesis data provides a rich dataset that informs on sequence sensitivity to tauopathy folds uncovering residues that are predicted to break fold compatibility or enhance it. We further demonstrate that the large dataset can be reduced to suggest key mutations that may help tune tau folding into defined states. This approach also identifies residue networks that distinguish closely related AD- and CTE-like fibril folds. We selected AD and CTE as the primary comparison because their ordered tau cores are globally similar but differ in localized R3/R4 packing, providing a stringent test of whether small sequence changes can discriminate related folds. We further show that alanine scanning alone does not sample the full amino-acid design space because it removes native side-chain bulk rather than testing larger or chemically distinct replacements. We find that the PAM4-containing amyloidogenic region of tau can independently drive amyloid formation and that pairwise substitutions at S341 and S352 remodel local packing within this region. In both minimal tau peptides and the 297-391 tau fragment, the S341L/S352I design enhances aggregation and produces seeding behavior consistent with a shift toward a CTE-like aggregate state. Cell lines harboring the double mutant show selectivity for CTE seeds compared to AD or CBD tauopathies. Lastly, aggregates derived from seeding WT 4R/4R tauRD biosensors with recombinant 297-391 S341L/S352I fibrils have similar templating behavior compared to CTE-derived aggregates. These findings support a model in which tau strains can be rationally tuned through local sequence changes that alter motif-motif interactions, providing a foundation for engineering disease-relevant amyloid models, conformation-selective biosensors and new strategies to interfere with pathological tau propagation.

## Results

### Computational saturation mutagenesis identifies residue networks that distinguish tau fibril folds

To define sequence features that govern tau fibril compatibility, we developed an in silico workflow that integrates disease-associated tau fibril structures, saturation mutagenesis and residue-contact analysis leveraging our previously established mutagenesis method adopted to amyloid fibrils (Fig. 1a).^11^ The analyzed tau repeat domain comprises microtubule-binding repeats R1-R4 and an R′ extension and includes the aggregation-prone regions (APRs) ^275^VQIINK^280, 306^VQIVYK^311, 337^VEVKSE^342^ and ^350^VQSKIGSLDNITH^362^ (herein Polymorphic amyloid motif of repeat 4, PAM4). Tau fibril conformations from AD (PDB ID 6HRE), CTE (PDB ID 6NWP), CBD (PDB ID 6TJX), AGD (PDB ID 7P6D), PSP (PDB ID 7P65) and GGT (PDB ID 7P66) were used as structural templates, providing six distinct packing environments across the ordered tau core (Fig. 1a). At every resolved position, each of the 19 non-native amino-acid substitutions were evaluated in 35 independent calculations. These calculations generated residue-level ΔΔG profiles in Rosetta Energy Units (REU), together with backbone RMSD values, that report the predicted energetic and structural consequences of mutation within each fibril fold (Fig. 1a). The distributions first revealed constraints that were shared across tau folds. Although most substitutions remained within a comparatively narrow energetic and structural range, several mutation classes produced long unfavorable tails in both ΔΔG and RMSD. Bulky hydrophobic or aromatic substitutions, proline and charged side chains were particularly disruptive when introduced into tightly packed sites, whereas smaller substitutions were more frequently tolerated (Supplementary Fig. 1). Energetic and structural disruption were related but not interchangeable. Some mutations incurred a large energetic penalty without extensive backbone displacement, whereas others sampled elevated RMSD with more modest energetic effects (Supplementary Fig. 2). We therefore retained both measurements when identifying potentially extreme or globally disruptive substitutions. Using ΔΔG = 138 REU and RMSD = 5 Å as outlier thresholds retained more than 93% of each dataset for comparative analysis, while isolating a small set of extreme cases for separate inspection (Supplementary Fig. 2).

**Figure 1.**
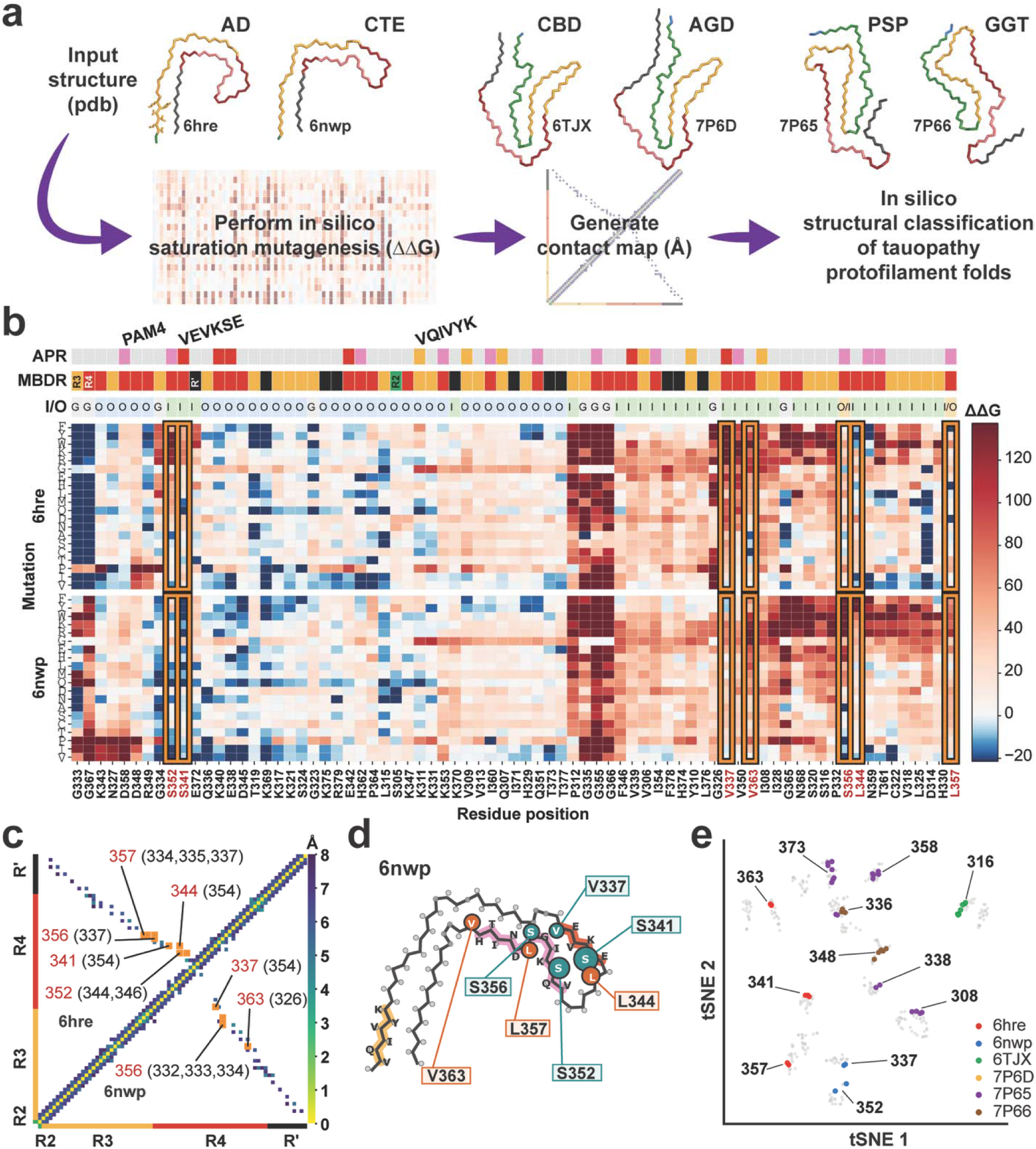
In silico structural classification identifies fold-discriminating mutations across tauopathy protofilaments. a. Computational workflow applied to AD (6HRE), CTE (6NWP), CBD (6TJX), AGD (7P6D), PSP (7P65) and GGT (7P66) tau fibrils. b. Hierarchical clustering of saturation-mutagenesis ΔΔG values (REU) for AD and CTE. The ΔΔG values were winsorized at the 5th and 95th percentiles for visualization. The color scale reports predicted ΔΔG values, with red indicating substitutions predicted to be destabilizing and blue indicating substitutions predicted to be stabilizing or more tolerated. Orange boxes mark positions with fold-distinct energetic profiles. The analyzed microtubule-binding domain repeat (MBDR) region comprises R1-R4 and R′: aggregation-prone regions (APRs) are annotated above the sequence. The upper annotations indicate APRs: VQIINK (green), VQIVYK (orange), VEVKSE (red) and PAM4 (pink), microtubule-binding domain repeat (MBDR) organization (R1 in blue, R2 in green, R3 (orange), R4 (red) and R4’(black)) and whether each residue is buried inside the fibril (I) or outside and solvent accessible (O) across the analyzed fibril templates. c. Split AD/CTE contact map. Contacts are Cβ-Cβ distances ≤8 Å (Ca for glycine); orange boxes mark contacts present in one fold but not the paired fold. d. CTE protofilament with fold-biased sites mapped as spheres. Teal labels mark the CTE-biased group (V337, S341, S352 and S356) and orange labels mark the AD-biased group (L344, L357 and V363). Sphere size is schematic and does not encode effect magnitude. APRs are VQIVYK (yellow), VEVKSE (red) and PAM4 (pink). e. t-SNE projection of the six-fold energetic profiles combined with contact features. Gray points are mutations without a fold-selective signature; colored points identify the fold for which the combined profile is most distinctive (6HRE, red; 6NWP, blue; 6TJX, green; 7P6D, yellow; 7P65, purple; 7P66, brown). Panels B-E were plotted in Visual Studio Code.

Residue-resolved ΔΔG heat maps showed that mutational sensitivity was strongly nonuniform along the tau sequence and depended on the surrounding fibril architecture (Supplementary Figs. 3). Broadly sensitive regions were concentrated in R3 and R4, including positions within and adjacent to ^306^VQIVYK^311, 337^VEVKSE^342^ and PAM4 amyloidogenic motifs and were enriched at buried packing interfaces.^11,12,23^ Native glycine and other small residues often formed sharply constrained signatures in which bulky, aromatic, charged, or proline substitutions were strongly unfavorable. By contrast, other positions tolerated multiple chemically distinct substitutions, producing localized neutral or favorable profiles even within otherwise constrained sequence segments. Hierarchical clustering of saturation-mutagenesis profiles grouped structure-amino acid substitution pairs according to their effects across tau fibril-core positions. Comparison of cluster occupancy and centroid distances revealed the extent to which mutational responses were shared within and differed between the tauopathy folds. Clustering positions across all six folds reorganized the heat map into groups with shared energetic behavior and exposed smaller groups whose mutational responses differed among the tauopathy templates (Supplementary Fig. 4). Removing the extreme ΔΔG/RMSD outliers (Supplementary Fig. 5) altered the visual ordering of these clusters but did not create new fold-selective patterns (Supplementary Fig. 6). The excluded mutations predominantly mapped to broadly constrained small-residue sites and therefore represented disruption of conserved packing or backbone geometry rather than evidence of fold specificity.

Pairwise energetic analyses were organized around structurally related comparisons of AD with CTE, CBD with AGD and PSP with GGT (Fig. 1a and Supplementary Figs. 7 and 8). In the hierarchical clustering comparison of AD and CTE saturation mutagenesis data, the strongest fold-discriminating signal localized to the R3/R4 region surrounding ^337^VEVKSE^342^ and PAM4. V337, S341, S352 and S356 formed the CTE-biased group shown in teal, whereas L344, L357 and V363 formed the AD-biased group shown in orange (Fig. 1b). The substitution maps further showed that fold selectivity was dependent on side-chain chemistry: individual positions could tolerate hydrophobic, aromatic, polar or charged substitutions in one structural background while disfavoring the same substitutions in the other fold. Thus, the ΔΔG maps identified both the positions and the specific amino-acid chemistries most likely to alter AD- or CTE-compatible packing. To define the structural basis for these energetic differences, we next compared residue-contact maps for each pair of fibril structures. Contacts were defined by Cβ-Cβ distances of 8 Å or less, using Cα for glycine and were considered fold-distinguishing when present in one structure but separated by more than 8 Å in the paired structure. This analysis connected local differences in mutational sensitivity to changes in long-range packing because the same sequence position can engage different interaction partners in different fibril folds. In the AD/CTE comparison, the energetically selected residues contributed differential contacts linking the R3 and R4 segments (Fig. 1c). Mapping the energetically and structurally selected sites onto the fibril models showed that they form spatially connected packing interfaces rather than contiguous linear motifs (Fig. 1d). In the CBD/AGD comparison, L282, L284, Q288 and S293 in R2 were linked with Y310 and S316 in R3 and E380 in R′, defining a network that spans separated sequence segments (Supplementary Figs. 7). The PSP/GGT network was broader and included N286 and C291 in R2, I308 and C322 in R3; K331, Q336, E338, D348, S352, D358 and N359 across R3/R4 and T373 near the R′ extension (Supplementary Figs. 8). In the CBD/AGD comparison, fold-distinguishing contacts linked residues in R2 with sites in R3, R4 and R′, revealing alternative packing among noncontiguous sequence segments (Supplementary Fig. 9a). The PSP/GGT comparison showed a broader redistribution of contacts spanning R2 through R′, consistent with more extensive differences in long-range organization between these folds (Supplementary Fig. 9b). Together, the energetic maps, differential contact plots and structural projections show that fold selectivity arises from distinct networks of interacting residues distributed across multiple packing interfaces rather than from a single universal amyloidogenic region.

Finally, dimensionality reduction of the combined energetic and contact features separated a subset of mutations with fold-selective signatures from the larger background of approximately 12,000 substitutions with shared or weakly distinguishing behavior (Fig. 1e). Mutations at the same residue frequently clustered into discrete interaction modules associated with a particular fibril template, indicating that fold selectivity reflects specific combinations of position, side-chain chemistry and structural environment rather than residue identity alone. These modules included mutations at positions distributed throughout the ordered core, including residues 308, 316, 336-341, 348, 352, 357-363 and 373. Several positions produced distinct clusters for the members of a structural pair, consistent with the same site supporting different amino-acid chemistries in alternative fibril architectures. Structural mapping of these fold-associated mutations revealed that they occupy spatially connected packing interfaces rather than a single contiguous sequence segment (Supplementary Fig. 10). In the AD/CTE pair, the selected sites converge on the interface between the ^337^VEVKSE^342^ and PAM4-containing regions, while the CBD/AGD and PSP/GGT comparisons identify broader networks spanning R2, R3, R4 and R′. The sets of tolerated substitutions also differed between paired structures, showing that fold discrimination depends not only on where a mutation occurs but also on the chemical properties of the introduced side chain. Together, the dimensionality-reduction and structural-mapping analyses define predicted networks of shared constraints and fold-selective interactions across the six tauopathy structures and provide a prioritized set of experimentally testable substitutions expected to alter fibril compatibility and strain-selective propagation.

### Alanine mutations in the amyloid core do not discriminate AD and CTE folds

To test whether computational predictions of fibril compatibility correspond to cellular tau aggregation, we employed a monomer incorporation assay in which cells stably propagate a tau strain and a tau mutant is tested for its ability to incorporate into the propagating aggregate.^11,15^ Systematic mutagenesis across the tau sequences yield incorporation profiles that mirror in silico predictions of stabilizing sites and are consistent with the structures.^11,15^ We selected 15 positions spanning the high-, intermediate- and low-destabilization ranges of the AD in silico alanine scan (Fig. 2a and Supplementary Fig. 11a). These sites sampled diverse regions of the ordered tau core. I308 and I328 lie within or adjacent to canonical aggregation-prone motifs, whereas D314, G335, S341, L344, S352, I354, G355, D358, V363, G367, L376 and

**Figure 2.**
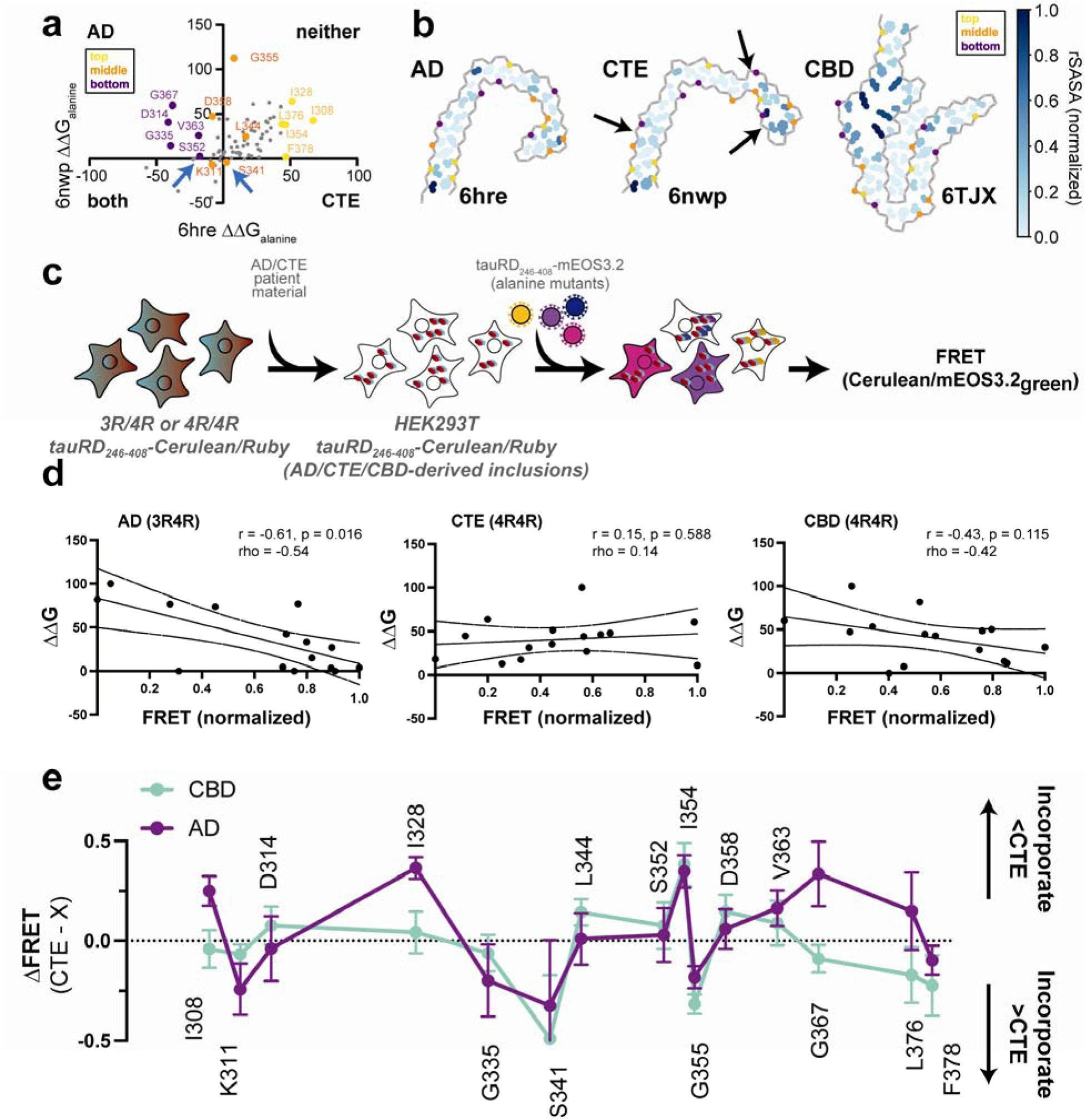
Computationally predicted fibril compatibility correlates with cellular tau incorporation in a polymorph-dependent manner. a. Comparison of Rosetta-predicted alanine-substitution ΔΔG values for the AD (PDB 6HRE) and CTE (PDB 6NWP) tau fibril folds. Positive values indicate predicted destabilization relative to the native residue. Mutations in the upper-right and lower-left quadrants are destabilizing in neither or both folds, respectively, whereas the remaining quadrants identify substitutions preferentially destabilizing to AD or CTE. The 15 positions selected for cellular analysis are labeled and colored according to the high-destabilization (“top,” gold), intermediate or mixed-effect (“middle,” orange) and low-destabilization or comparatively tolerated (“bottom,” purple) groups derived from the AD alanine scan. Blue arrows indicate S341 and S352. b. Locations of the tested residues mapped onto the AD (6HRE), CTE (6NWP) and CBD (6TJX) fibril structures. Residues are colored by normalized relative solvent-accessible surface area (rSASA), from buried (light cyan) to solvent exposed (dark blue). Arrows highlight the structurally divergent turn region in CTE. c. Schematic of the cellular incorporation assay. HEK293T biosensor cells expressing 3R/4R or 4R/4R tauRD tauRD_246-408_-Cerulean/Ruby reporters were seeded with AD-, CTE-, or CBD-derived brain material to generate intracellular tau aggregates and subsequently transduced with tauRD_246-408_*-*mEOS3.2 reporters carrying the indicated alanine substitutions. Incorporation into pre-existing aggregates was quantified by FRET between CFP and the green-emitting form of mEOS3.2. d. Relationships between unscaled predicted ΔΔG and normalized cellular FRET incorporation for AD-derived aggregates in 3R/4R cells (left), CTE-derived aggregates in 4R/4R cells (middle) and CBD-derived aggregates in 4R/4R cells (right). Lines show linear regressions with 95% confidence intervals. Pearson r with a two-sided p value and Spearman ρ are shown. e. Position-resolved differences in normalized incorporation between CTE-derived aggregates and AD- or CBD-derived aggregates. Purple, ΔFRET = CTE - AD; green, ΔFRET = CTE - CBD. Positive values indicate greater incorporation into CTE-derived aggregates, whereas negative values indicate greater incorporation into the comparator-derived aggregates. Points and error bars represent mean and ± SD from three replicate wells.

F378 represent additional packing environments. S341, L344 and S352 were also identified by the AD/CTE contact-map analysis, directly linking the cellular test set to the structural screen. Comparison of the predicted alanine-substitution effects in AD and CTE separated positions that were unfavorable in both folds from those preferentially tolerated by one amyloid fold (Fig. 2a). Mapping the selected sites onto the AD, CTE and CBD fibril structures further showed that they are distributed throughout the ordered core and encompass buried, partially buried and solvent-exposed environments (Fig. 2b). The structurally divergent CBD fold was included as a control for determining whether relationships identified using the AD-derived energetic landscape extended to a distinct tau conformation.

We next measured the incorporation of the corresponding alanine mutants into cells that propagate AD, CTE or CBD cellular tau aggregates using a FRET-based incorporation assay (Fig. 2c).^11,15^ AD- or CTE-derived patient material was introduced into HEK293T cells stably co-expressing WT 4R tauRD_246-408_-mRuby3 and either WT 4R or 3R tauRD_246-408_-mCerulean3 (Fig. 2c, 4R/4R and 3R/4R). After propagation of the tauopathy-associated fold, cells were transduced with tauRD-mEOS3.2 constructs containing individual alanine substitutions or controls. Incorporation of the mutant tau into the propagating tauopathy aggregates produced a FRET signal between the CFP-labeled aggregate and mEOS3.2-green and quantified using flow cytometry (Supplementary Fig. 12). Incorporation measurements were normalized within each tauopathy seed background to enable comparisons among substitutions and across AD-, CTE- and CBD-derived aggregates (Supplementary Fig. 11b).

The relationship between predicted energetic compatibility and cellular incorporation depended strongly on the tau fold (Fig. 2d). For AD-derived aggregates, predicted destabilization showed a moderate inverse correlation with normalized FRET incorporation (Pearson 0.61, 0.016, Spearman 0.54). Thus, alanine substitutions predicted to impose larger energetic penalties on the AD fibril generally incorporated less efficiently into AD-like aggregates. By contrast, no relationship was detected for CTE-derived aggregates (Pearson 0.15, 0.588, Spearman 0.14). CBD-derived aggregates showed a weaker inverse trend that did not reach statistical significance (Pearson 0.43, 0.115, Spearman 0.42). The weaker relationship for CBD was consistent with the selection of the tested substitutions from the AD energetic landscape rather than from the energetic extremes of the CBD fold. Together, these comparisons indicate that the alanine-scanning calculations capture incorporation competence more effectively for AD-derived aggregates than for CTE- or CBD-derived aggregates.

To determine whether differences in residue burial could explain the fold-dependent predictive performance, we compared the computational and cellular measurements with solvent-accessible surface area. We defined ΔSASA as the solvent-accessible surface area of a residue in the isolated monomer minus its accessible area in the modeled fibril, positive values therefore indicate burial upon fibril assembly. Predicted ΔΔG correlated positively with ΔSASA in the AD structure (Pearson 0.61, 0.016, Spearman 0.61), consistent with larger energetic penalties at more deeply buried sites (Supplementary Fig. 11c). The relationship was not detected for CTE (Pearson 0.24, 0.385, Spearman 0.04) and was weaker and nonsignificant for CBD (Pearson 0.42, 0.116, Spearman 0.42). Cellular incorporation itself did not correlate significantly with ΔSASA for any of the three folds, indicating that burial alone was insufficient to predict whether an alanine mutant would be incorporated.

Pairwise analysis nevertheless showed that structural differences between AD and CTE contributed to their distinct incorporation profiles. The difference in FRET incorporation between CTE and AD correlated with the corresponding difference in ΔSASA (Pearson 0.66, 0.007, Spearman 0.62, Supplementary Fig. 11d). Residues that were more buried in one tauopathy fold therefore tended to favor incorporation into that fold. By contrast, the CTE-AD difference in FRET did not correlate with the difference in total predicted ΔΔG (Pearson 0.09, 0.741, Spearman 0.07). Decomposition of the energy terms indicated that the energetic differences between the folds were distributed across several terms, with prominent contributions from backbone conformation and van der Waals packing at specific positions (Supplementary Fig. 11e). These results suggest that localized differences in fibril packing and solvent exposure provide information about strain-selective incorporation that is not fully captured by the total alanine-substitution score.

Direct comparison of the cellular profiles further revealed position-specific differences between CTE and the AD and CBD backgrounds (Fig. 2e). Here, positive ΔFRET values indicate greater incorporation into CTE-derived aggregates, whereas negative values indicate greater incorporation into AD- or CBD-derived aggregates. The direction and magnitude of these differences varied across the tau core rather than being confined to a single contiguous sequence element. Several substitutions favored CTE incorporation, whereas others favored AD or CBD, defining a distributed set of residues that distinguish the cellular compatibility of these folds. The AD- and CBD-derived incorporation profiles were therefore distinct from CTE, although the two comparisons did not produce identical residue-level patterns. Because the relatively open CTE architecture may leave some sites insufficiently packed for alanine substitutions to produce a strong energetic bias, we expanded the computational analysis to larger hydrophobic substitutions. Comparison of valine and isoleucine substitutions identified positions at which increasing side-chain volume was predicted to be more favorable for CTE than for AD (Supplementary Fig. 11f,g). This analysis suggests that substitutions introducing larger hydrophobic side chains may stabilize packing environments that are poorly filled by alanine and therefore provide a more effective strategy for generating CTE-selective variants. Together, these results demonstrate that computational fibril-compatibility energy scores can identify residues that regulate strain-dependent tau incorporation, particularly for AD-like aggregates. They also show that the relationship between fibril-core energetics and cellular propagation varies among tau folds. Total alanine-substitution energies were less predictive for CTE, whereas local differences in burial and packing better explained the divergence between AD and CTE incorporation. Thus, effective prediction of strain-selective compatibility may require considering both the magnitude of the energetic perturbation and the specific structural environment in which the substitution occurs.

### Designed tau variants alter predicted fibril compatibility and aggregation behavior

To test whether sequence variation can be used to tune compatibility with disease-associated tau fibril folds, we used structure-guided design to identify combinations of substitutions predicted to alter packing within AD- and CTE-like tau conformations (Fig. 3). Pairwise mutational energy maps revealed that candidate substitutions have distinct predicted effects depending on the fibril template (Fig. 3a). In particular, substitutions centered around S341/S352 and S356/L357 showed fold-dependent energetic patterns, with stronger predicted effects in the CTE template than in the AD template, suggesting that local sequence changes can selectively reshape compatibility with specific tau folds (Fig. 3a, b). A subset of designed variants was selected for experimental testing based on predicted energetic effects and their locations within the tau amyloid core. These designs included single and combinatorial substitutions near residues S341, S352, S356 and L357, positioned within or adjacent to structural elements that differ between AD- and CTE-associated tau fibrils (Fig. 3b). Mapping these variants onto AD- and CTE-like structures showed that the targeted residues cluster within buried or interfacial regions of the fibril core, where they are well positioned to influence packing, local conformational stability and strain compatibility (Fig. 3c,d).

**Figure 3.**
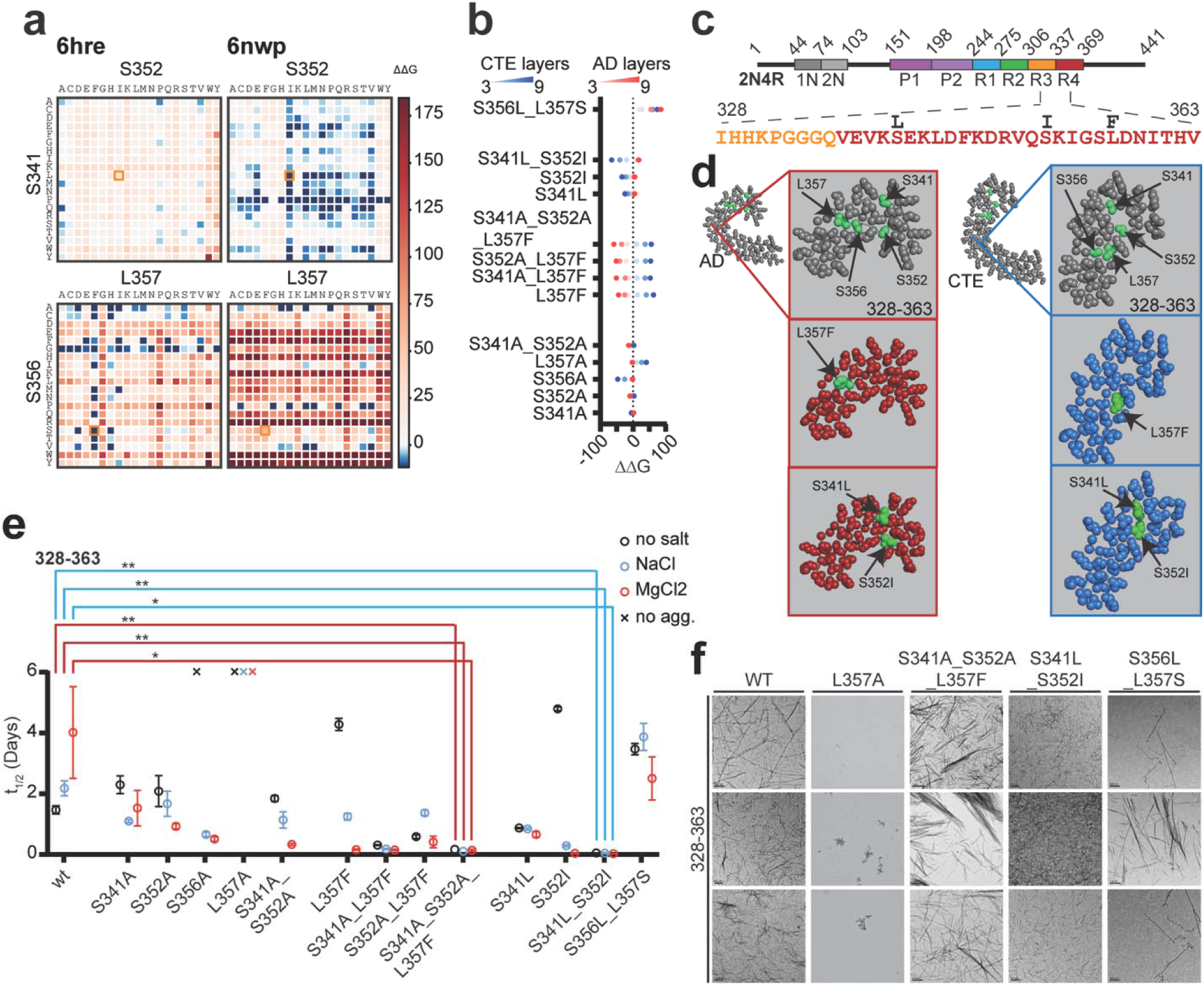
Designed substitutions alter predicted fold compatibility and accelerate tau aggregation. a. Saturation-mutagenesis maps for S341, S352, S356 and L357 in AD and CTE templates; boxes identify substitutions selected for testing. b. Predicted ΔΔG values for the selected single and combinatorial designs in three- and nine-layer AD and CTE models. c. Sequence context of the tau_328-363_ design region. d. Selected residues mapped onto AD (red) and CTE (blue) fibril cores; insets show local packing around L357, S341, S356 and S352. e. Aggregation half-times under no-salt, NaCl, MgCl_₂_ and no-aggregation control conditions. Points represent three replicate wells and error bars show mean ± SD. Brackets indicate pairwise comparisons with WT within the same buffer condition; *p < 0.05 and **p < 0.01. f. Endpoint negative-stain TEM images collected after the six-day aggregation assay. Scale bars are shown in each micrograph.

To determine how computationally selected mutations affect tau aggregation, we performed Thioflavin T (ThT) fluorescence aggregation assays using a minimal tau fragment spanning residues 328-363, which contains structural elements that differ between the AD and CTE polymorphs (Fig. 3c-e). We reasoned that the tau fragment spanning 328-363 can be stabilized by interactions between the N-terminal ^337^VEVKSE^342^ and C-terminal PAM4 amyloidogenic motifs thus defining a minimal fragment that may fold into a defined turn. We also tested a shorter tau_335-357_ peptide, however, this fragment did not produce detectable ThT fluorescence and no aggregates were observed by TEM, indicating that residues outside this shorter segment are required for assembly under these conditions (Supplementary Fig. 13a). By contrast, tau_328-363_ aggregated robustly despite lacking the canonical ^306^VQIVYK^311^ amyloid motif. WT tau_328-363_ aggregated with half-times of 1.46 ± 0.14 h in 10 mM sodium phosphate, 2.18 ± 0.25 h in 10 mM sodium phosphate with 200 mM NaCl and 4.01 ± 1.51 h in 10 mM sodium phosphate with 200 mM MgCl_2_ (Fig. 3e and Supplementary Table 1). These buffer conditions were selected to test the aggregation promiscuity of the designed sequences, as ionic conditions have been reported to influence tau aggregation and bias assembly toward CTE- or AD-like folds.^4^

Alanine mutagenesis showed that individual residues within the 328-363 region make distinct contributions to aggregation (Fig. 2e,c, Supplementary Fig. 13b,c and Supplementary Table 1). S341A and S352A did not abolish aggregation but altered buffer sensitivity. S341A slowed aggregation in no-salt buffer relative to WT but accelerated aggregation in NaCl and MgCl_2_, with half-times of 2.29 ± 0.29 h, 1.09 ± 0.05 h and 1.53 ± 0.58 h, respectively. S352A showed a similar trend, with half-times of 2.08 ± 0.51 h, 1.67 ± 0.41 h and 0.92 ± 0.09 h in no-salt, NaCl and MgCl_2_ conditions, respectively. The S341A/S352A double mutant further enhanced salt-dependent aggregation, particularly in MgCl_₂_, where the half-time decreased to 0.33 ± 0.03 h. By contrast, L357A completely suppressed detectable ThT aggregation under all tested conditions, indicating that L357 is essential for assembly of this minimal tau core. S356A also reduced aggregation in no-salt buffer but aggregated rapidly in NaCl and MgCl_2_, suggesting that this site contributes to buffer-dependent stabilization rather than serving as an absolute requirement for fibril formation (Fig. 2e,c, Supplementary Fig. 13b,c and Supplementary Table 1).

AD-stabilizing designs centered on L357 and further supported the importance of this region (Fig. 3 and Supplementary Fig. 14). Although L357A eliminated detectable aggregation, the conservative aromatic substitution L357F retained aggregation and became strongly accelerated in salt-containing buffers, with half-times of 4.28 ± 0.20 h in no salt, 1.25 ± 0.11 h in NaCl and 0.16 ± 0.02 h in MgCl_2_ (Fig. 2e,f, Supplementary Figs. 13b,c and 14 and Supplementary Table 1). Combining L357F with S341A or with S341A/S352A produced rapid aggregation across all conditions. S341A/L357F aggregated with half-times of 0.31 ± 0.02 h, 0.18 ± 0.02 h and 0.15 ± 0.02 h, whereas S341A/S352A/L357F aggregated with half-times of 0.17 h, 0.11 h and 0.14 ± 0.03 h in no-salt, NaCl and MgCl_₂_ conditions, respectively (Fig. 2e, f, Supplementary Fig. 14 and Supplementary Table 1). These results suggest that L357 is a key determinant of assembly and that aromatic packing at this position can compensate for or enhance the effects of mutations at S341 and S352.

Mutations predicted to stabilize the CTE-like fold produced stronger and more consistent effects on aggregation (Fig. 2e,c, Supplementary Fig. 15 and Supplementary Table 1). S341L accelerated tau_328-363_ aggregation across all buffer conditions, with half-times of 0.87 ± 0.02 h, 0.84 ± 0.03 h and 0.66 ± 0.09 h in no-salt, NaCl and MgCl_₂_ conditions, respectively. S352I showed a more pronounced salt dependence, aggregating slowly without added salt but rapidly in NaCl and MgCl_2_, with half-times of 4.79 ± 0.06 h, 0.29 ± 0.03 h and 0.03 ± 0.01 h, respectively. The combined S341L/S352I mutant produced the fastest and most buffer-independent aggregation profile, with half-times of 0.049 ± 0.006 h, 0.045 ± 0.024 h and 0.024 ± 0.006 h in no-salt, NaCl and MgCl_2_ conditions, respectively. These values were significantly faster than WT under each condition, supporting the prediction that this pairwise substitution stabilizes an aggregation-competent CTE-like interaction network (Fig. 2e,c, Supplementary Fig. 15 and Supplementary Table 1). By contrast, the S356L/L357S design, which was not predicted to stabilize the AD- or CTE-fold, aggregated more slowly than any of the strongest AD- and CTE-stabilizing variants, with half-times of 3.47 ± 0.19 h, 3.87 ± 0.45 h and 2.50 ± 0.71 h in no-salt, NaCl and MgCl_2_ conditions, respectively (Fig. 2e,f and Supplementary Table 1). Thus, not all substitutions within this region increase aggregation, rather, aggregation depends on the specific chemical features introduced at positions that shape local packing. TEM analysis confirmed that ThT-positive reactions contained fibrillar assemblies, whereas poorly aggregating conditions showed reduced or absent fibril formation (Fig. 3f, Supplementary Figs. 13c, 14b and 15b). Together, these data indicate that residues within tau_328–363_ are sufficient to drive amyloid formation independently of ^306^VQIVYK^3^^11^ and that mutations predicted to stabilize a CTE- or AD-like packing interface, particularly S341L/S352I and L357F-containing variants, markedly enhance aggregation of this minimal tau core. The alanine mutants further show that this region contains both permissive positions and essential packing residues, with L357 emerging as a critical determinant of fibril formation.

### Cryo-EM structural analysis shows the PAM4 motif as the nucleus of aggregation for the tau_328-363_ constructs

To determine how these mutations affect amyloid formation and fibril architecture, we first used cryo-EM to determine the structure of WT tau_328-363_ aggregates formed under no-salt conditions (Fig. 4 and Supplementary Fig. 16). Images were collected and processed in RELION.^34^ After postprocessing, the sequence was fit into the masked map using CryoFold^35^, followed by refinement in Phenix.^36^ Based on FSC analysis, the WT tau_328-363_ amyloid structure was determined at 3.46 Å resolution (Fig. 4a, Supplementary Fig. 16 and Supplementary Table 2). The fibrils have helical screw symmetry, a crossover distance of 463 Å, a twist of -1.84° and a rise of 4.73 Å (Fig. 4a, Supplementary Fig. 16 and Supplementary Table 2). The structure contains two protofilaments, each composed of one short chain spanning residues 349-363 and one long chain spanning residues 337-363 (Fig. 4a). No additional polymorphs were resolved in this dataset, likely reflecting the restricted conformational space available to the short peptide sequence. We next determined the cryo-EM structure of the S341L/S352I tau_328-363_ amyloid at 2.2 Å resolution (Fig. 4a, Supplementary Fig. 17 and Supplementary Table 3). Compared to WT, the mutant fibrils had a shorter crossover distance of 334 Å, a twist of -2.55° and a rise of 4.73 Å (Fig. 4a, Supplementary Fig. 17 and Supplementary Table 3). The mutant structure is also composed of two protofilaments, however, each protofilament contains two copies of chain A spanning residues 349-363 rather than the chain A/chain B arrangement observed in WT (Fig. 4a,b). Thus, the S341L/S352I mutations preserve amyloid formation but remodel the packing architecture of the tau_328-363_ core. Structural comparison of WT and S341L/S352I showed that the shared chain A segment adopts a very similar backbone conformation in both structures (Fig. 4b). The polymorphic amyloid motif of repeat 4, PAM4, spanning residues ^350^VQSKIGSLDNITH^362^, is nearly superimposable between WT chain A and S341L/S352I chain A, with an RMSD of 0.28 Å and 0.40 Å, respectively (Fig. 4b). The largest local difference occurs at position 352, where the S352I mutation introduces a bulkier hydrophobic side chain that forms a stabilizing interaction with V363 and leads to a more open conformation compared to WT (Fig. 4b). This increased steric bulk likely prevents the remaining sequence from curving back to form the stabilizing salt-bridge interactions observed in chain B of the WT tau_328-363_ structure (Fig. 4a,b). Although S341L was not resolved within the ordered core of the S341L/S352I structure, because this residue lies in the N-terminal disordered region of the mutant fibril, in silico mutational analysis predicted that both S341L and S352I destabilize the WT tau_328-363_ fold (Fig. 4c,d). S341L was predicted to have a larger effect on the WT structure than S352I, with ΔΔG values of 15.5 and 5.9, respectively. These data suggest that the mutations do not simply stabilize the original WT packing arrangement. Instead, they appear to disfavor the WT chain A/chain B architecture while allowing the PAM4-containing segment to assemble into an alternative, more symmetric amyloid structure.

**Figure 4.**
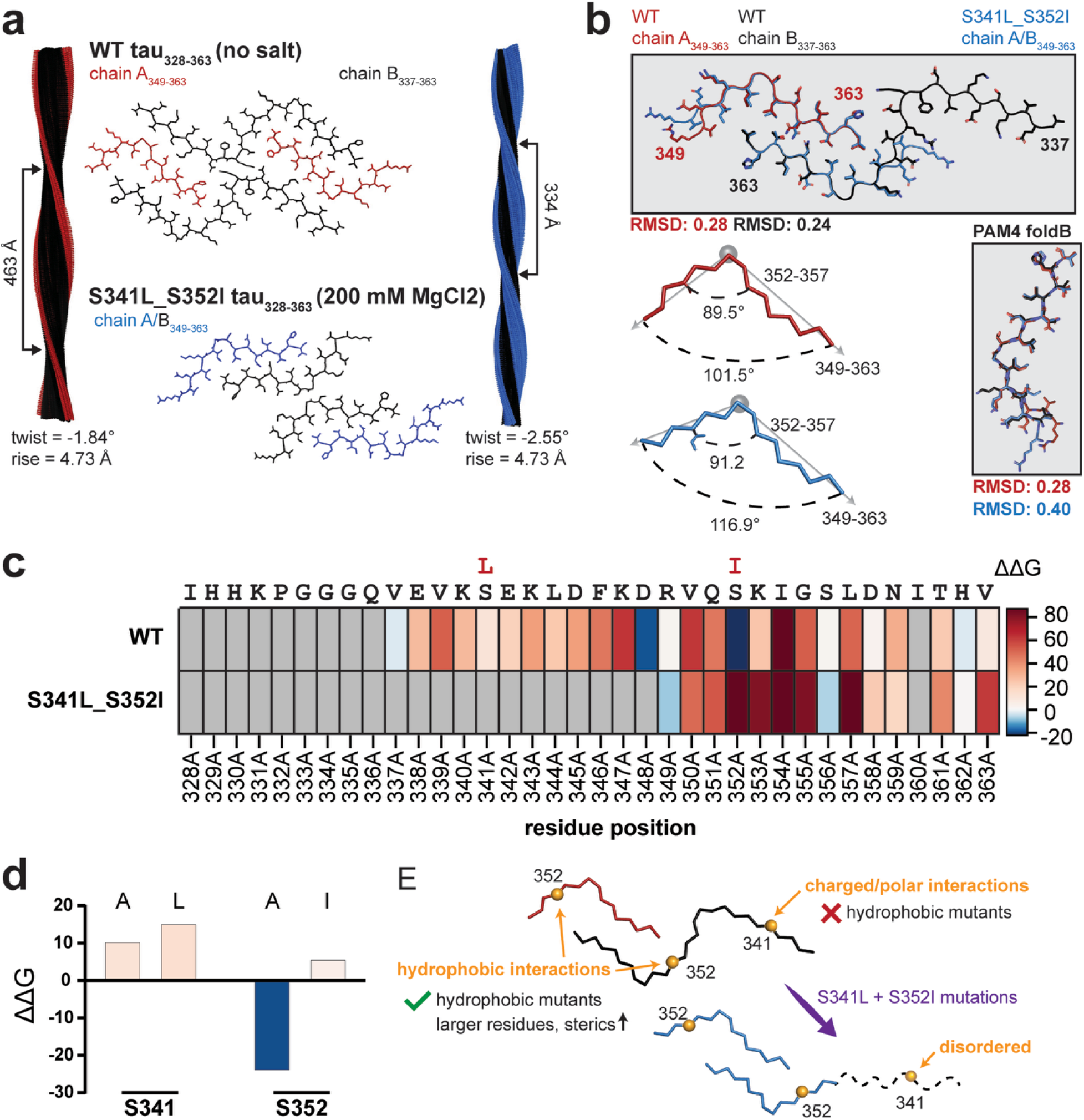
CTE-biased S341L/S352I mutations remodel packing between VEVKSE and PAM4. a. Cryo-EM structures of WT and S341L/S352I tau_328-363_ fibrils and the local packing region spanning residues 349-363. b. Structural overlay of the shared ordered segments. RMSDs are Cα RMSDs after PyMOL alignment of the shared ordered residues. c. Residue-level ΔΔG profiles for WT and S341L/S352I; gray denotes positions not evaluated or not resolved in the modeled core. d. Predicted effects of substitutions at S341 and S352. e. Model in which S341L/S352I changes local hydrophobic and polar packing and permits an alternative PAM4-centered architecture. The data establish that PAM4-centered packing can support assembly under these conditions but do not distinguish whether PAM4 or VEVKSE engages first during nucleation.

Residue-level energetic analysis further supported this interpretation. Compared with WT, the S341L/S352I structure showed a shifted ΔΔG profile across the 328-363 region, with the strongest stabilizing contributions concentrated within the PAM4-containing segment (Fig. 4c). The in silico alanine scan identified residues 354, 352, 357, 355 and 353 as the top stabilizing positions in the S341L/S352I structure, with ΔΔG values ranging from 150 to 80. By comparison, the top stabilizing residues in the WT tau_328-363_ structure were 354, 347, 350, 355 and 339, with ΔΔG values ranging between ∼100-50 (Fig. 4c). Thus, the mutant structure places greater energetic weight on the central PAM4 interface, particularly around S352I and neighboring residues. Substitution analysis at S341 and S352 within the WT context further highlights the importance of the native serine residues. Particularly at S341, hydrophobic replacements were disfavored as this site is surrounded by polar or charged residues (Fig. 4c-e). At S352, steric clashes play an important role in energetics as an isoleucine was found to be unfavorable whereas an alanine is predicted to be stabilizing (Fig. 4c-e). This provides a structural rationale for the enhanced aggregation behavior of S341L/S352I, rather than globally disrupting the fibril fold, the mutations remodel local side-chain packing and favor a more hydrophobic, ordered interaction network within the tau_328-363_ core (Fig. 4e). Together, these structural and computational analyses indicate that S341L/S352I shifts tau_328-363_ from the WT chain A/chain B packing arrangement toward a PAM4-centered amyloid architecture (Fig. 4). The finding that tau_328-363_ forms amyloid despite lacking both ^275^VQIINK^280^ and ^306^VQIVYK^311^ supports the idea that PAM4 is an independent amyloid-driving element. These structures show that PAM4-centered packing is sufficient to support assembly under these conditions even though the fragment also contains ^337^VEVKSE^342^, they do not establish that PAM4 universally dominates ^337^VEVKSE^342^ during nucleation. This is consistent with prior work showing that PAM4 can self-assemble into multiple amyloid conformations resembling tauopathy-associated folds.^12^ Although ^306^VQIVYK^3^^11^ is considered to initiate tau aggregation^11,23,37,38^, all ex vivo tauopathy structures reported to date also contain PAM4 and these data suggest that PAM4 has intrinsic aggregation and structural propensity and can strongly influence tau polymorphism.^12^ Thus, regions outside the canonical ^306^VQIVYK^311^ motif can support tau assembly and targeted mutations within 328-363 can redirect the energetic and structural landscape of tau amyloid.

### Cellular seeding assays suggest that S341L/S352I is compatible with a CTE-like fold

To test whether the S341L/S352I double mutant can bias tau assembly toward a CTE-like fold, we introduced these mutations into the recombinant tau fragment spanning residues 297-391. WT tau_297-391_ and S341L/S352I tau_297-391_ constructs were purified recombinantly and their aggregation propensity was compared using a ThT fluorescence assay under no-salt, NaCl and MgCl_₂_ conditions (Supplementary Fig. 18a,b). S341L/S352I tau_297-391_ aggregated more readily than WT tau_297-391_, with half-times of 0.97 ± 0.07 h, 1.67 ± 0.02 h and 4.53 ± 0.92 h in no-salt, NaCl and MgCl_2_ conditions, respectively, compared with WT tau_297-391_ half-times of no detectable aggregation, 3.09 ± 0.63 h and 4.10 ± 0.43 h under the same conditions (Supplementary Fig. 18a,b and Supplementary Table 4). Thus, as observed for the tau_328-363_ peptide, the S341L/S352I mutations increase the intrinsic aggregation propensity of a longer tau fragment that contains the AD/CTE fibril core. Although S341L/S352I tau_297-391_ showed evidence of aggregation under all three buffer conditions, TEM revealed that the no-salt and NaCl reactions primarily contained amorphous material, whereas the MgCl_2_ condition produced fibrillar assemblies suitable for further analysis (Supplementary Fig. 18b). Cryo-EM grids were prepared from the MgCl_2_ fibrils, however, initial image processing indicated that the filaments were largely straight and lacked sufficient helical twist for high-resolution three-dimensional reconstruction. Structural analysis was therefore limited to 2D classification (Supplementary Fig. 18c). Because cryo-EM could not resolve the atomic structure of these assemblies, we next used cellular seeding and incorporation assays to infer whether the designed aggregates propagated with AD-like or CTE-like behavior (Fig. 5). We first determined whether the endpoint products of the recombinant tau aggregation reactions were competent to seed intracellular tau assembly. WT and S341L/S352I tau_297-391_ aggregation products were delivered into HEK293T tau biosensor cells and seeded aggregation was quantified by FRET 72 h later (Fig. 5a). Initial experiments used NK13 cells expressing tauRD with the frontotemporal dementia-associated P301S mutation.^39^ These cells were optimized for sensitive detection of tau aggregates and are relatively permissive with respect to seed conformation.^39^ WT and S341L/S352I endpoint samples produced robust FRET responses when the corresponding aggregation reactions contained NaCl or MgCl_2_, whereas samples generated without salt produced little or no activity (Fig. 5b). Thus, the conditions that generated ThT fluorescence and fibrillar material by transmission electron microscopy also produced assemblies capable of templating intracellular tau aggregation.

**Figure 5.**
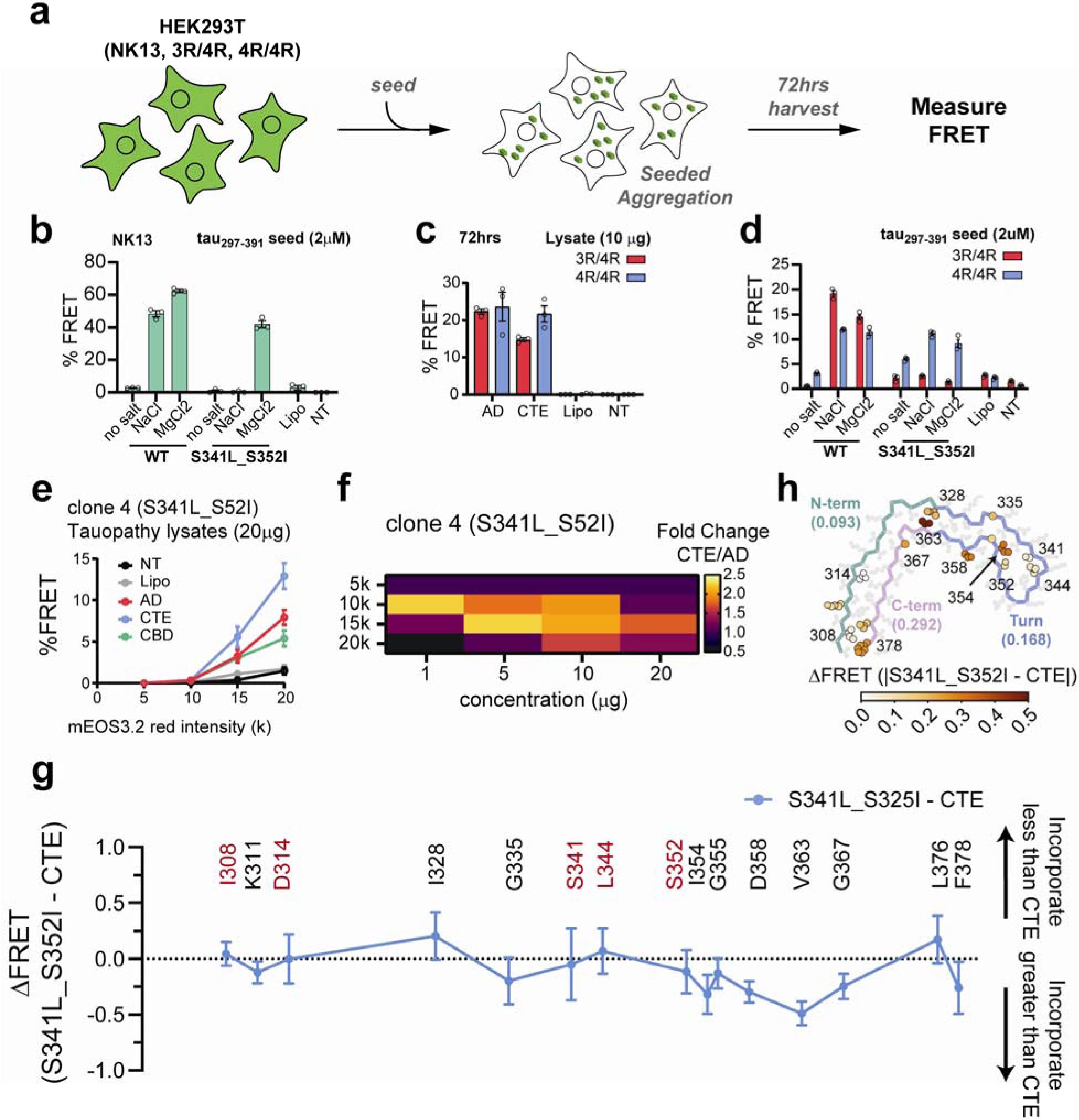
S341L/S352I confers CTE-like seeding and incorporation profiles in tau biosensors. a. Schematic of the cellular seeding assay. HEK293T biosensor cells were treated with recombinant tau aggregates or tauopathy-derived brain lysates and seeded intracellular aggregation was quantified by FRET. NK13 cells express P301S *tauRD*_246-378_-mCerulean3/mClover3 reporters, whereas the 3R/4R and 4R/4R biosensors express paired *tauRD*_246-408_-mCerulean3/mRuby3 reporters. b. Seeding of NK13 cells with 2 μM recombinant WT or S341L/S352I tau_297-391_ endpoint samples generated in the absence of salt or in the presence of NaCl or MgCl_2_. Cells were analyzed 96 h after treatment. Lipofectamine-only (Lipo) and no treatment (NT) cells served as controls. c. Seeding of 3R/4R and 4R/4R biosensors with 10 μg of AD- or CTE-derived brain lysate. FRET was measured 72 h after treatment. d. Seeding of 3R/4R and 4R/4R biosensors with 2 μM recombinant WT or S341L/S352I tau_297-391_ endpoint samples. Cells were analyzed 96 h after treatment. e. Expression-dependent seeding of the S341L/S352I tauRD_246-408_-mEOS3.2 clone 4 reporter by 20 μg of AD-, CTE-, or CBD-derived brain lysate. The percentage of FRET-positive cells was quantified within mEOS3.2 red-intensity gates centered at 5k, 10k, 15k and 20k arbitrary units. Lipofectamine-only and non-transduced cells served as controls. f. Heat map showing the fold difference between CTE- and AD-induced FRET in clone 4 as a function of brain-lysate concentration and mEOS3.2 red-intensity gate. Values greater than 1 indicate preferential seeding by CTE-derived lysate. g. Position-resolved comparison of alanine-mutant incorporation into S341L/S352I- and CTE-derived aggregates. Differences were calculated as normalized FRET(S341L/S352I-seeded) – normalized FRET(reference-seeded); values near zero indicate similar incorporation into the two aggregate types. Residues highlighted in red correspond to positions selected from the AD/CTE structural analysis. Across all 15 positions, the mean absolute ΔFRET was 0.181 for CTE. h. Absolute differences between the S341L/S352I and CTE incorporation profiles mapped onto the mature CTE fibril fold. Residues are colored according to mean absolute ΔFRET and regional labels (green, N-terminal; blue, turn; purple, C-terminal) indicate the mean absolute difference for the N-terminal segment, turn and C-terminal segment. Unless otherwise indicated, data represent three replicate wells and are shown as mean ± SD.

We next tested whether these recombinant assemblies exhibited differential activity in reporter cells that were previously developed to distinguish between AD- and CTE-derived seeds^15^. These biosensors express paired 3R/4R or 4R/4R tauRD_246-408_ constructs fused to mCerulean3 and mRuby3 (Fig. 5a). Consistent with previous findings^15^, AD-derived brain lysate seeded the 3R/4R and 4R/4R reporters with comparable efficiency, whereas CTE-derived lysate produced a larger FRET-positive population in the 4R/4R reporter than in the 3R/4R reporter (Fig. 5c). This differential response established the relative preference of the 4R/4R reporter for CTE-associated seeding activity under these conditions. Lipofectamine-only and no treatment controls produced negligible FRET. WT and S341L/S352I tau_297-391_ endpoint samples were then introduced into both reporter backgrounds (Fig. 5d). WT aggregates formed in NaCl or MgCl_2_ seeded both cell lines, with greater activity in the 3R/4R reporter. By contrast, S341L/S352I aggregates generated under either salt condition preferentially seeded the 4R/4R reporter. This reversal in reporter preference was not attributable to a general increase in seeding activity: the mutant seeds were less active than WT in the 3R/4R cells but retained substantial activity in the 4R/4R cells. The S341L/S352I substitutions therefore altered the conformational compatibility of the recombinant aggregates and shifted their cellular seeding profile toward that produced by CTE-derived material.

To test the compatibility of the S341L/S352I sequence directly with disease-derived seeds, we generated cells stably expressing WT or S341L/S352I tauRD_246-408_ C-terminally tagged with mEOS3.2. Partial photoconversion of mEOS3.2 using UV light produces green- and red-emitting species that form a FRET pair upon tau assembly. Transduced cells were sorted to generate polyclonal populations and an S341L/S352I single-cell clone with higher reporter expression, clone 4, was isolated for further analysis. Cells were seeded with recombinant or patient derived AD or CTE aggregates and the green-red FRET was quantified using flow cytometry (Supplementary Fig. 20). Because reporter abundance influenced the measured fraction of FRET-positive cells, seeding was evaluated across red-fluorescence gates centered at 5k, 10k, 15k and 20k arbitrary units.

At the 5k-intensity gate, both WT and S341L/S352I polyclonal reporter populations responded to disease-derived tau seeds in a concentration-dependent manner (Supplementary Fig. 19a). WT cells responded strongly to both AD and CTE lysates and showed substantial activity with CBD lysate at intermediate concentrations. The S341L/S352I polyclonal cells generally produced lower absolute FRET responses than WT but showed a greater relative response to CTE than to AD. Accordingly, the CTE/AD response ratio was higher in the S341L/S352I polyclonal population than in WT across multiple lysate concentrations (Supplementary Fig. 19b). Comparison with the original 3R/4R and 4R/4R reporters at the available concentration further showed that the S341L/S352I polyclonal population provided strong discrimination between CTE- and AD-derived seeds. Because clone 4 expressed the S341L/S352I reporter at a higher level, its responses were examined across the series of red mEOS3.2 intensity gates. At the lowest 5k intensity gate, clone 4 showed little response to any disease-derived lysate. Seeding increased markedly with reporter abundance, however, and CTE-derived lysate produced the strongest response at the higher intensity gates (Fig. 5e). The preference for CTE was also dependent on seed concentration: the CTE/AD response ratio was most pronounced at intermediate and high lysate concentrations within the 10-20k intensity gates (Fig. 5f). Complete dose-response measurements confirmed that AD-, CTE- and CBD-derived lysates produced progressively greater seeding as both lysate concentration and reporter intensity increased, with CTE yielding the largest response at 20 μg (Fig. 5f and Supplementary Fig. 19c). Recombinant S341L/S352I tau_297-391_ seeds also produced an expression-dependent response, confirming that the reporter could detect aggregates formed from the same sequence. These results demonstrate that reporter abundance affects assay sensitivity but does not account for the preferential compatibility of S341L/S352I tau with CTE-derived seeds.

### Cellular incorporation assays suggest that aggregates templated from recombinant S341L/S352I seeds resemble on-pathway CTE intermediates

We next asked whether the similarity between S341L/S352I and CTE extended beyond the double-mutant reporter to a broader residue-level incorporation signature. We compared normalized incorporation profiles for 15 alanine substitutions in cells containing AD-, CTE-, CBD-, or S341L/S352I-derived aggregates (Supplementary Fig. 21a). Although incorporation varied substantially by residue and aggregate type, the S341L/S352I profile was generally most similar to CTE. Its differences from AD were larger at selected positions, whereas its differences from CBD were more extensive across the ordered core (Supplementary Fig. 21b). The CTE-like behavior of S341L/S352I therefore reflects a distributed pattern of sequence compatibility rather than an effect confined to residues 341 and 352.

Direct comparison of the S341L/S352I and CTE incorporation profiles identified both closely matched positions and localized differences (Fig. 5g). I308, D314, S341, L344 and S352 showed relatively small differences between the two profiles, whereas larger deviations occurred towards the C-terminus around D358, V363 and G367. Mapping the absolute differences onto the CTE structure revealed a corresponding regional pattern (Fig. 5h). The average difference was lowest for the segment comprising residues 306-323 and 365-378, intermediate within the turn formed by residues 335-355 and greatest for the segment encompassing residues 323-334 and 356-364. Thus, the overall similarity between the S341L/S352I and CTE profiles was accompanied by localized divergence within a specific structural region.

Comparison with available CTE fibril structures further illustrated the conformational variability of this region. The previously determined CTE fold and the major- and minor-interface architectures observed in additional CTE structures shared a common overall topology but differed in the relative positions of individual structural segments (Supplementary Fig. 21c,d). The largest variation in the major-interface architecture occurred within the segment encompassing residues 323-334 and 356-364, which was also the region with the greatest average difference between the S341L/S352I and CTE incorporation profiles (Supplementary Fig. 21d, Segment 2). These observations suggest that the residual differences may reflect conformational heterogeneity within the CTE structural family rather than uniform incompatibility of S341L/S352I with CTE-like assemblies.

Together, the reciprocal seeding experiments support a model in which S341L/S352I biases tau toward a CTE-like propagation state. Recombinant S341L/S352I aggregates preferentially seeded the 4R/4R reporter favored by CTE-derived material, polyclonal and clonal S341L/S352I reporters showed enhanced selectivity for CTE over AD seeds and the residue-level incorporation profile of S341L/S352I aggregates more closely resembled CTE than AD or CBD. These cellular measurements do not establish that the recombinant aggregates reproduce the atomic structure of an ex vivo CTE fibril. Indeed, their predominantly straight morphology did not support a high-resolution cryo-EM reconstruction. Nevertheless, the convergence of reporter selectivity, disease-seed compatibility and mutational incorporation profiles indicates that S341L/S352I promotes assemblies with conformational features preferentially compatible with CTE-associated tau propagation.

## Discussion

This study establishes a structure-guided framework for identifying sequence determinants of tau amyloid compatibility. Saturation mutagenesis across six disease-associated tau fibril structures revealed both shared constraints and fold-selective interaction networks distributed throughout the ordered core. Experimental analysis of the AD-CTE comparison showed that combining substitutions within one such network alters aggregation kinetics, fibril packing and cellular propagation. In particular, S341L/S352I produced assemblies and reporters that were preferentially compatible with CTE-derived seeds. These findings demonstrate that local sequence changes can bias tau toward a CTE-like propagation state, while also showing that compatibility with a mature fibril template does not by itself ensure formation of a single defined amyloid structure. The computational screen additionally identified candidate fold-selective substitutions for AD, CBD, AGD, PSP and GGT (Supplementary Fig. 10), providing a resource for testing whether the same principles extend across the broader tau conformational landscape. These predictions remain to be experimentally evaluated. Nevertheless, the convergence of computational energetics, biochemical assembly, cryo-EM and cellular seeding supports a model in which tau polymorphism is governed by distributed but locally modular interaction networks. Individual substitutions reshape particular packing environments, whereas combinations of substitutions can shift the balance among competing assembly and propagation states.

### Saturation mutagenesis reveals fold-selective sequence constraints

The repetitive cross-β architecture and availability of atomic fibril models make amyloids particularly amenable to residue-level energetic analysis. Amyloid cores contain recurring interactions within and between β-strands, including steric zipper-like interfaces and repeated contacts between adjacent fibril layers.^37,40^ These features allow many mutational effects to be related to specific packing environments. Amyloids are not structurally simple, however, long-range contacts, conformational coupling, polymorphism and pathway-dependent assembly can all influence the consequences of a substitution. Saturation mutagenesis is therefore most informative when interpreted as a measure of compatibility with a defined structural template rather than as a direct prediction of aggregation or fold selection. Across approximately 12,000 sequence-structure mutants, the analysis revealed a rich and heterogeneous energetic landscape. Some positions were broadly sensitive to substitution across tau fibril templates, consistent with shared requirements for backbone geometry or core packing. Other positions tolerated chemically diverse substitutions or displayed distinct preferences among related folds. Proline, bulky aromatic residues and substitutions at tightly constrained glycine sites were often unfavorable, whereas many other changes were tolerated or predicted to improve compatibility. The native tau sequence therefore appears to occupy one region of a much larger sequence space capable of supporting cross-β structure. Because the same tau sequence can access multiple disease-associated folds, it may be compatible with several conformations without being optimized for any one of them. This residue-level view complements generative approaches such as ProteinMPNN, which can identify globally compatible amyloid sequences with the capacity to cross-template wild-type proteins.^19,26^ Generative designs frequently contain multiple simultaneous substitutions, making it difficult to attribute their behavior to particular residues or interactions. Saturation mutagenesis instead defines which side-chain chemistries are tolerated, disruptive or favorable at each position and allows their energetic contributions to be examined. The two strategies are therefore complementary: generative methods efficiently explore higher-order sequence solutions, whereas saturation mutagenesis provides an experimentally interpretable route for identifying and combining substitutions at structurally informative sites.

### Alanine scanning identifies hotspots but incompletely defines fold compatibility

The experimental alanine scan illustrates both the value and the limitations of single-chemistry mutagenesis. Removing side-chain bulk identified positions important for AD-compatible incorporation and highlighted regions in which amyloid-core energetics correspond to cellular propagation. Predicted alanine-substitution energies correlated with incorporation into AD-derived aggregates but not into CTE-derived aggregates and alanine mutants had limited ability to distinguish the two propagation states. Local differences in residue burial and packing were more informative than total alanine-substitution energies for explaining the divergence between AD and CTE. Alanine scanning primarily tests the contribution of the native side chain; it does not establish which alternative chemistry is optimal at that position. This distinction becomes important in relatively open or underpacked environments, where removing a side chain may have little effect even though introducing a larger hydrophobic or aromatic group could favor one conformation. Saturation mutagenesis extends hotspot identification by sampling the full range of amino-acid chemistries. In the AD-CTE comparison, this broader search nominated substitutions at S341, S352, S356 and L357 that would not have emerged from alanine scanning alone and provided a rational basis for testing combinatorial designs.

### Local sequence changes bias the tau assembly landscape

The S341L/S352I variant provides a proof of principle that substitutions selected from a fold-comparison landscape can produce a measurable conformational bias. The double mutant altered the kinetics and buffer dependence of tau assembly, changed fibril morphology and local packing within the tau_328-363_ fragment and enhanced compatibility with CTE-derived seeds in cellular assays. Recombinant S341L/S352I aggregates preferentially seeded the 4R/4R reporter favored by CTE material, while S341L/S352I reporter cells displayed increased discrimination between CTE- and AD-derived seeds. In addition, aggregates generated from the mutant produced a residue-level incorporation profile that more closely resembled CTE than AD or CBD. Together, these orthogonal measurements support the conclusion that S341L/S352I shifts tau toward a CTE-compatible propagation state. The data do not establish that S341L/S352I recreates the atomic structure of an ex vivo CTE fibril. The mutant fibrils were morphologically heterogeneous and their predominantly straight morphology did not support a high-resolution reconstruction. Moreover, the incorporation profile retained localized differences from CTE, particularly around residues 323-334 and 356-364. This region also varies among available CTE structures, raising the possibility that the mutant occupies an intermediate or an alternative member of a broader CTE-compatible conformational family. Defining that relationship will require higher-resolution structures of the recombinant and cell-propagated assemblies. This distinction is important for amyloid design. A sequence can be energetically compatible with a mature fold yet reach a different endpoint because nucleation, intermediates, cofactors and solution conditions reshape the accessible pathway. Structure-guided mutations should therefore be viewed as changing the probability of competing assembly states rather than uniquely specifying one structure. The ability of two substitutions to produce a reproducible bias is nevertheless encouraging: it indicates that the amyloid landscape can be shifted through targeted manipulation of local interaction networks, even when complete structural control has not yet been achieved.

### PAM4 defines an amyloid-competent module and suggests pathway-dependent motif engagement

A second important finding is that tau_328-363_ forms ordered amyloid in the absence of the canonical ^306^VQIVYK^311^ motif. Cryo-EM showed that PAM4 contributes the core of these assemblies and that S341L/S352I remodels packing within this region. PAM4 is therefore not merely a passive element of mature tau fibrils; within this fragment, it can support ordered assembly together with interactions involving ^337^VEVKSE^342^. These observations reinforce the view that tau contains multiple amyloid-competent modules with distinct structural preferences. The results also raise the possibility that the order in which these modules engage helps determine the resulting fibril. In FL tau, ^275^VQIINK^280,306^VQIVYK^311,337^VEVKSE^342^ and PAM4 can participate in different combinations of intra- and intermolecular contacts. Early engagement of VQIVYK could favor one set of downstream interactions, whereas PAM4-centered nucleation could bias the chain toward another. S341L/S352I may influence this competition by stabilizing PAM4-centered interactions or altering the accessibility of alternative pathways. Our experiments do not directly establish the temporal order of motif engagement and testing this model will require methods that resolve early assembly intermediates. It nevertheless provides a mechanistic framework for understanding how local substitutions can produce large changes in kinetics, morphology and cellular compatibility.

### Limitations and future directions

Several limitations define the current scope of the framework. The energy calculations evaluate compatibility with fixed backbone models of mature fibrils and do not directly capture nucleation barriers, alternative polymorphs, solvent effects, cofactors, post-translational modifications or the ensemble of intermediates that precedes fibril formation. The cellular FRET assays measure incorporation and propagation compatibility rather than atomic structural identity. The tau_328-363_ fragment isolates a mechanistically informative interaction network but may not reproduce the assembly landscape of FL tau. Finally, experimental validation focused on the AD-CTE comparison, the predicted determinants of CBD, AGD, PSP and GGT compatibility remain hypotheses. These limitations also identify productive next steps. Large, complete computational mutagenesis datasets paired with systematic experimental measurements should enable models tailored to specific outcomes, including mature-fold compatibility, aggregation kinetics and cellular propagation. Structural characterization of designed variants in longer tau constructs and in seeded cellular systems will be essential for determining when energetic compatibility produces the intended fold and when it instead stabilizes an intermediate or alternative polymorph. Extending combinatorial tests to the other tauopathy networks will reveal whether local sequence optimization is a general strategy for navigating disease-associated tau conformations.

### Implications for detection of species barriers and therapeutic targeting

The results support a sequence-based model of amyloid transmission analogous to a molecular species barrier. For a soluble tau molecule to bind to a growing fibril, its sequence must be able to adopt the conformation and satisfy the packing interactions presented by the seed.^11,15^ A substitution that disrupts this compatibility should reduce templating even if it remains compatible with another fold, conversely, a substitution that improves a particular packing environment may promote selective incorporation. The S341L/S352I reporter illustrates how this principle can be exploited experimentally. Although it does not identify an atomic structure by itself, its preferential response to CTE-derived material provides a route toward reporters that distinguish closely related propagation states using sequence rather than tau isoform alone. More broadly, the same framework could guide the design of conformation-biased amyloid models and ultimately, inhibitors of strain amplification. Mutations that enhance compatibility can help stabilize experimentally accessible states, whereas variants or peptides that engage a fibril edge without completing the required interaction network may act as dominant-negative inhibitors. These applications remain prospective but they follow directly from the sequence-structure relationships defined here. Tau amyloid formation emerges not as the consequence of a single dominant motif but as a modular folding problem in which multiple sequence elements cooperate and compete. By resolving both shared constraints and fold-selective determinants, saturation mutagenesis provides a tractable route for probing this landscape and for engineering tools that detect, model and potentially disrupt pathological tau propagation.

## Methods

All the research described in this work complies with ethical regulations. All human samples utilized in these experiments were derived from deceased subjects. All autopsies were performed only with the permission of the decedent’s next of kin or other person(s) legally authorized to provide such permission. All human materials utilized in these experiments were deidentified and no individually identifiable protected health information was available to the investigators using these materials. The UT Southwestern Institutional Review Board has determined that such studies are exempt from human subjects regulations as codified in federal law.

### In silico ΔΔG calculations

The in silico mutation scan^11^ relies on a Rosetta framework adapted from the Flex ddG protocol described by Barlow et al.^41^ developed to interpret protein interface stability by comparing bound and unbound states. Here, we focus on how the single residue substitutions change the energy of the whole fibril assembly. The protocol published previously by Mullapudi et al.^11^ can be found under total_mutation_ddg_simple at https://zenodo.org/records/7549915. In this protocol, the resfile-driven residue selection and the paired wild-type/mutant comparison is retained but the backrub ensemble sampling and the bound/unbound interface calculation were not included.

For AD (6HRE) and CTE (6NWP) polymorphs, the input structures were a 3-layer, 6-layer, or 9-layer protofilament generated in Pymol (version 2.6.0), as described in^11^. For CBD (6TJX), AGD (7P6D), PSP (7P65), GGT (7P66) and the structures of the peptides generated in this work 9-layer protofilaments were generated. A single protofilament was used to reduce the required computational resources. The mutation site was defined along with the number of layers for each input structure to ensure that all layers were mutated.

For energetic minimization of the input structures, ref2015 score function, lbfgs Armijo nonmonotone minimizer with all backbone and side-chain torsions free, tolerance 0.000001 and maximum of 5000 iterations and 1 REU convergence threshold were used. Within the WT structure, the residues were repacked and minimized while in the mutant structures, the mutation was introduced followed by repacking and minimization. The mean change in energy over 35 independent replicates, drawn from random seed, for each mutation was reported as a ΔΔG (i.e., ΔG_mut_ - ΔG_wt_).^11^ Rosetta calculations were performed using version 2020.08 (Git commit cb1caba). To identify shared mutational-response patterns across tau fibril structures, saturation-mutagenesis data from six structures (PDB IDs: 6HRE, 6NWP, 6TJX, 7P6D, 7P65, and 7P66) were combined into a matrix in which each row represented a structure-amino acid substitution pair and each column represented a residue position. Rows were clustered according to their mutational profiles across positions using agglomerative hierarchical clustering with Ward’s linkage and a Euclidean distance metric, implemented with the linkage function in SciPy.^42^ The resulting hierarchy was partitioned into six clusters using the fcluster function. For each structure, the set of occupied clusters was determined and cluster overlap between 1) AD and CTE, 2) CBD and AGD and 3) PSP and GGT and for all six fibril structures was quantified using the Jaccard index. Structure-level centroids were calculated by averaging the mutational profiles across amino acid substitutions, and pairwise Euclidean distances between centroids were used to compare within- and between-disease-group similarity.

### Generation of tauopathy fold contact maps

Each distance measured and plotted for the contact maps was generated using an in-house python script. In short, the pdb coordinates for the protofilament was loaded into PyMol. Using the Cβ position, the script calculates the Euclidean 3D distance between every residue. For glycine, the Cα position was substituted. Next, paired, split maps were generated to illustrate the structural similarity between specific tauopathy folds (e.g., 6HRE vs. 6NWP, 6TJX vs. 7P6D and 7P65 vs. 7P66). Contacts ≤ 8 Å were plotted with longer distances masked white. The script then identifies residue contacts unique to each structure-pair and boxes them in orange.

### Generation of contact-informed tSNE of distinguishing residue positions

For the contact-informed tSNE analysis, the data set was first restricted to the finalized distinguishing residue positions identified from the paired tauopathy fold comparisons (L282, L284, N286, Q288, C291, S293, I308, Y310, S316, C322, K331, Q336, V337, E338, S341, L344, D348, S352, S356, L357, D358, N359, V363, T373, E380). For each mutation at these positions, a six-dimensional saturation mutagenesis profile was generated from the ΔΔG values across the six structures: 6HRE, 6NWP, 6TJX, 7P6D, 7P65 and 7P66. Next, to incorporate structural context, residue-level contact features were added using the paired Cβ-Cβ contact maps. For each residue, contact-derived features included structure-specific contact counts, unique-contact counts and nearest-contact distance summaries. The saturation mutagenesis and contact-derived features were independently standardized. Contact features were then weighted relative to the saturation mutagenesis features and concatenated with the six ΔΔG values to create a combined feature vector for each mutation. tSNE was performed on this combined feature matrix using Euclidean distance, PCA initialization, automatic learning rate and a fixed random seed for reproducibility.

### Structural comparison and RMSD calculations

Structural overlays were generated with the align command in PyMOL version 2.6.0 using Cα atoms from the shared ordered residue range. Reported RMSD values are Cα RMSDs after PyMOL’s default iterative outlier rejection, residues outside the shared ordered segment were excluded.

### Peptide synthesis

WT tau_328-363_ and mutant peptide constructs were synthesized with N-terminal acetylation and C-terminal amidation purified to > 95% by Genscript. Peptides were disaggregated using trifluoroacetic acid (TFA, Sigma-Aldrich) at a ratio of 2mg of peptide/200 mL of TFA and kept overnight in a fume hood to remove organic solvent, followed by brief resuspension in 200 mL MQ water prior to being flash frozen in liquid nitrogen and lyophilization. Disaggregated peptide powder was stored in -20°C until needed.

### Tau aggregation assays

Each assay was conducted over 6 days under the exact same conditions, 4 mg/ml protein, 10 mM sodium phosphate (NaPO_4_, Sigma-Aldrich), 10 mM dithiothreitol (DTT, Sigma-Aldrich), 3 mM Thioflavin-T (ThT, Sigma-Aldrich) at 37 °C (no salt, 200 mM NaCl, or 200 mM MgCl_2_).^14^ These aggregation assays were carried out in triplicate using 96-well clear-bottom plates (Corning) in a Tecan Spark plate reader. Measurements were recorded at an excitation of 446 nm and emission of 486 nm, with 10 sec of shaking at 216 rpm prior to each read, every 15 min. Data points were plotted as averages with standard deviation and t_1/2_ for each experiment was determined using a non-linear fit in AmyloFit^43^ (Table 1 and 4 in supplementary information).

### Transmission electron microscopy

Each aggregate sample was washed to remove excess ThT prior to imaging. A 50 mL aliquot of each aggregate sample was centrifuged at 50,000 x g for 1 hr, supernatant removed and pellet resuspended in 50 mL of milli-Q water, followed by a second round of centrifugation at 50,000 x g for 30 min, supernatant removed once more and pellet resuspended in 50 mL of milli-Q water. A 5 mL aliquot of each aggregate sample was then placed on a glow-discharged, 300 mesh formvar/carbon supported copper grid for 45 sec before blotting with filter paper, followed by a 5 mL aliquot of a 2% uranyl acetate solution for 45 sec before blotting with filter paper, followed by a 1 min swirl within a 50 mL droplet of fresh milli-Q water on parafilm. All TEM grids were then blotted with filter paper before air drying at RT for ≥ 10 min.

### Cryo-EM grid preparation and data collection/processing

Aggregate samples of interest were diluted with MQ water based on preliminary TEM screening (e.g., 1/4^th^ dilution). A 4 mL aliquot from each dilution was placed on a 300 mesh Quantifoil R 1.2/1.3 copper grids prior to plunge freezing utilizing a Leica GP2 (95% humidity, 278 K, 3 sec wait time, 5.0 front blot force). Grids were mounted onto an autogrid ring and secured with a C-clip prior to screening using a FEI Talos Glacios transmission electron microscope. Cryo-EM datasets were then collected utilizing a FEI Titan Krios transmission electron microscope. See Table 2 and 3 in the supplementary information for further details.

### Recombinant tau purification

Tau_297-391_ constructs were purified using similar protocols. WT and S341L/S352I tau_297-391_ were cloned into a pET28a vector using gene blocks cloned from IDT. Plasmids were transformed into BL21(DE3) competent cells and plated on Luria Broth (LB) agar (Sigma-Aldrich) plates supplemented with 50 mg/ml kanamycin (Sigma-Aldrich). Colonies were grown in 5 mL aliquots of LB (Research Products International, RPI) overnight in the presence of 50 mg/ml kanamycin. The tau_297-391_ overnight cultures were used to induce 2 L of LB auto-induction media and grown for 8 hr at 37 °C, 250 rpm followed by 16 hr at 24 °C, 220 rpm. The large LB auto-induction cultures were pelleted at 5,000 x g for 20 min. Cell pellets were resuspended in 40 mL lysis buffer per 1 L of culture (10 mM sodium phosphate, 10 mM dithiothreitol (DTT), 1 mM phenylmethylsulfonyl fluoride (PMSF, ThermoFisher Scientific), 10 mM ethylenediaminetetraacetic acid (EDTA, Sigma-Aldrich)). Resuspended cells were either lysed utilizing a homogenizer PandaPLUS 2000 at ∼15,000 psi for eight cycles or using a sonication probe at 50% amplitude. Lysate was kept on ice during homogenization. Following homogenization, the cell lysate was spun at 14,000 x g for 20 min.

The supernatant was collected and filtered using a 0.22 mm filter prior to loading the sample onto a 150 mL super loop attached to an AKTA pure chromatography system. Each tau construct was then loaded onto a 5 mL HiTrap SP column, washed with 5 column volumes of 10 mM sodium phosphate, 10 mM DTT buffer and eluted with 10 mM sodium phosphate, 10 mM DTT, 1 M sodium chloride buffer. Fractions containing Tau protein were collected, concentrated to 5 mL using appropriate MWCO MilliporeSigma AMicon Ultra-15 Centrifugal Filters (3k MWCO filters) and then loaded onto a chromatography column (Cytiva, Superdex 75 10/300). Tau protein was eluted with 10 mM sodium phosphate buffer. Purity was assessed using sodium dodecyl-sulfate polyacrylamide gel (Invitrogen) electrophoresis following each chromatography step. All tau protein was filtered with 0.22 mm filters prior to measuring protein concentration using a Pierce 660nm Protein Assay Kit. Tau protein was aliquoted, flash frozen and stored at -80 °C.

### Establishment of cellular S341L/S352I biosensor cell-lines

Lentivirus containing S341L/S352I tauRD (246-408), C-terminally fused to a photoconvertible clover/cerulean fluorescent protein (mEOS3.2), was generated as follows: 7.5 mL of TransIT (Mirus) was mixed with 142.5 mL of Opti-MEM (GIBCO) and incubated at room temperature for 5 min. This mixture was combined with psPAX2 (1.2 mg), VSV-G (0.4 mg) and FM5-CMV-S341L/S352I_246-408_ tau-mEOS (1 mg) plasmids and incubated at room temperature for 30 min before aliquoting 110 mL of the mixture drop-wise to a six-well dish containing HEK293T cell-line (ATCCCRL-3216) plated with 2 mL containing 300,000 cells, 24 hrs prior. Cell media (10% FBS, 1% Pen/Strep, 1% GlutaMax in Dulbecco’s modified Eagle’s medium) containing virus was collected after 72 hrs by centrifugation at 1000xg for 5 min. Supernatant was collected and aliquoted before freezing at -80 °C. HEK 293-T cells were transduced with the lentivirus and harvested 48 hrs later. Single cells were sorted into 96-well plates after the expression of the protein was confirmed by fluorescence. Individual cell lines were chosen based on expression, expanded and used for downstream experiments.

### Preparation of lysates from tauopathy patient material

Frontal cortex from tauopathy cases was added to 1× TBS supplemented with cOmplete ULTRA EDTA-free protease inhibitor cocktail (Roche) and homogenized with a PowerGen 125 tissue homogenizer (Fisher Scientific) to generate 10% w/v brain homogenates. Lysates were sonicated in a water bath (QSonica) for 10 min using 1-min on and 30-s off cycles at 65% amplitude, then centrifuged at 21,000 × g for 15 min at 4°C to remove debris. The supernatant was collected and total protein concentration was quantified by BCA assay (Thermo Scientific). Supernatants were aliquoted and stored at −80 °C until use in cell-based seeding or incorporation assays.

### Cell-based seeding assays

Three stable overexpression cell lines, NK13, 3R/4R and 4R/4R, previously developed^15,39^ were treated with several peptide aggregate samples. The NK13 cell line expresses a P301S_246-378_ tau-mCerulean3/mClover3 pairs whereas the 3R/4R and 4R/4R cell lines express ΔR2WT_246-408_ tau and WT_246-408_ tau or solely WT_246-408_ tau-mCerulean3/mRuby3 pairs, respectively. Specifically, NK13, 3R/4R, or 4R/4R cells were plated into 96-well cell-culture treated plates at 10,000 cells per well in 130 µL of medium (complete DMEM media with 10% FBS) 18-24 hrs before treatment with peptide or lysate aggregate samples. Fibrils were sonicated at an amplitude of 65 for 30 sec on and 30 sec off with a total on time of 10 min using a water bath sonicator (QSonica) at 4°C, prior to seeding. For seeding, 0.7 µL of Lipofectamine 2000(Invitrogen) and 9.3 µL Opti-MEM(Gibco) per well of a 96-well plate was incubated as a master-mix for 5 min at room temperature prior to being added to sonicated seeds. Opti-MEM was added to reach a desired concentration of peptide/tauopathy lysate (1 µM or 1-20 µg of total protein for patient-derived lysates, respectively). This mixture was incubated for 30 min, RT prior to being added dropwise to cells. Each experiment was performed as a biological triplicate.

### S341L/S352I seeding assays

For peptide seeding assays, cells were harvested 96 h after treatment, other cell-based seeding assays were harvested 48-72 h after treatment. Cells were incubated with 0.25% trypsin for 5 min at 37 °C, quenched with complete cell medium, transferred to 96-well round-bottom plates and centrifuged at 1,000 × g for 5 min. Cells were fixed in PBS containing 2% paraformaldehyde for 10 min and centrifuged again at 1,000 × g for 5 min. Finally, cells were resuspended in 200 µL DPBS and stored at 4 °C until analysis by flow cytometry. Before flow-cytometry analysis, S341L/S352I biosensor cells were partially photoconverted for 30 min to generate a green-red mEOS3.2 FRET pair.

### Cell-based incorporation assays

Cells expressing WT 3R/4R or 4R/4R tau (246–408) biosensors tagged with cyan or ruby fluorescent proteins were treated with recombinant fibrils or AD-, CTE-, or CBD-derived brain lysate. Cells were plated at 20,000 cells per well 24 h before seeding. For seeding, Lipofectamine 2000 (0.75 µL per well) and Opti-MEM (19.25 µL per well) were incubated for 5 min, followed by addition of 1 µM recombinant fibrils or 10 µg tauopathy lysate diluted to the desired concentration in Opti-MEM. The mixture was incubated for 30 min at room temperature and added dropwise to cells. After 72 h, treated cells were replated into a new 96-well plate. Medium was removed, 50 µL of 0.5% trypsin was added and cells were incubated for 5 min at 37 °C before quenching with 150 µL complete medium. After resuspension, 60 µL of cells was aliquoted into each of three wells containing 70 µL fresh media. Alanine-mutant tau246–408-mEOS3.2 lentivirus (20 µL) was then added to 130 µL of freshly plated, aggregate-harboring cells. Cells were harvested 72 h after reverse transduction and stored at 4 °C until analysis by flow cytometry.

### Lentivirus production for alanine mutants of tauRD-mEOS3.2

Alanine-mutant tau246–408-mEOS3.2 plasmids encoding fifteen alanine substitutions (residues 308, 311, 314, 328, 335, 341, 344, 352, 354, 355, 358, 363, 367, 376 and 378) were generated by Gibson assembly. HEK293T cells were plated at 300,000 cells per well in 6-well tissue-culture plates containing 2 mL complete medium. After 24 h, the plasmid of interest (1,000 ng) was incubated with psPAX2 (1,200 ng) and VSV-G (400 ng) for 5 min at room temperature. TransIT-293 (7.5 µL per well) and Opti-MEM (100 µL per well) were incubated separately for 5 min at room temperature and 107.5 µL of this mixture was added to the plasmid/helper-plasmid mixture. After 30 min at room temperature, the transfection mixture was added dropwise to cells. Virus-containing medium was collected 72 h after transfection, mixed with Lenti-X Concentrator at a 1:3 ratio and incubated overnight at 4 °C. The mixture was centrifuged at 1,500 × g for 1 h at 4 °C, the supernatant was aspirated and the pellet was resuspended in 200 µL PBS containing 25 mM HEPES. Concentrated virus was stored at −80 °C until use.

### Flow cytometry analysis

An Attune CytPix/CytKick Flow cytometer (ThermoFisher Scientific) was used to measure FRET within the fixed cells for cell-based seeding assays.

#### NK13 and S341L/S352I biosensor seeding assays

Using the NK 13 biosensors, mClover3, mCerulean3 and FRET were measured with the AmCyan channel, mCerulean3. To measure mCerulean3 and FRET signal, cells were excited with the 405 nm laser and fluorescence was captured with a 450/50m nm filter. To measure mClover3, cells were excited with a 488 laser and fluorescence was captured with a 525/50 nm filter. To quantify FRET, mCerulean3 bleed-through into the mClover3 and FRET channels was compensated for during data analysis using FlowJo v10 software (Treestar). The FRET gate was set according to biosensor cells that received lipofectamine alone and are thus FRET-negative. This allows for visualization of acceptor emission arising from the excitation of the donor at 405 nm. For each experiment, 10,000 double-positive events per replicate were analyzed and each condition was analyzed in biological triplicate.

Similarly, for S341L/S352I tauRD biosensors, mClover3, mRuby3 and PerCP (FRET between Clover and Ruby) were measured. mEOS3.2 is a photoconvertible fluorescent protein which on partial photoconversion forms a green-red FRET pair. Samples were gated on the acceptor intensity (mEOS3.2 red intensity: 5k, 10k, 15k, 20k arbitrary units) such that cells with similar intensities were analyzed to mitigate the contribution of differences in concentration leading to apparent changes in the fraction of FRET-positive cells for each condition.

3R/4R and 4R/4R tauRD incorporation assays. For 3R/4R and 4R/4R tau biosensors, mCerulean3, mRuby3, Qdot 650-A (mCerulean3/mRuby3 FRET), mClover3 and AmCyan-A (mCerulean3/mClover3 FRET) channels were measured. Cells were gated for double-positive expression of the mCerulean3 donor and mRuby3 acceptor, followed by quantification of FRET between this pair. Cells were then gated for high mClover3 intensity to select cells that received alanine-mutant virus, followed by quantification of FRET between mCerulean3 and mClover3 in the AmCyan-A channel. Full gating strategies are shown in Supplementary Figs. 12 and 20.

### Data normalization and statistical analysis

For incorporation assays, percent FRET was calculated as 100 × (FRET-positive events/high-mClover3 events). Within each seed dataset, replicate-level percentages were min-max normalized as (x - xmin)/(xmax - xmin) across the alanine-mutant series, WT and DP assay controls. Each mutant and seed condition were measured in three replicate wells, individual values are shown and summary values are reported as mean ± SD. For Figs. 2e and 5g, ΔFRET was calculated at each position as normalized FRET(CTE-seeded) - normalized FRET(reference-seeded) or FRET(S341L_S352I-seeded) - normalized FRET(reference-seeded) and profile similarity was summarized as the mean absolute ΔFRET across the fifteen positions. Correlation panel reports Pearson r with a two-sided p value and Spearman ρ. ThT aggregation curves and half-times were obtained from three replicate wells and are shown as mean ± SD.

## Data availability

The structural datasets generated during the current study are available in the Electron Microscopy Data Bank repository (https://www.ebi.ac.uk/emdb/) under accession numbers, EMD-70445 [https://www.ebi.ac.uk/emdb/EMD-70445] and EMD-77369 [https://www.ebi.ac.uk/emdb/EMD-77369]. The structural model that was fit into the density is available in the Protein Data Bank under PDB id 9OFW (https://www.rcsb.org/structure/unreleased/9OFW) and 36BU (https://www.rcsb.org/structure/unreleased/36BU). ThT aggregation data, ΔΔG calculations and cell-based aggregation data are available as Source Data. The raw saturation mutagenesis data is available on Zenodo under accession number 22131503 (<u>10.5281/zenodo.22131503</u>). Any other data sets generated during and/or analyzed during the current study are available from the corresponding author on reasonable request. PBD ID’s used in this study are: 6hre [http://doi.org/10.2210/pdb6HRE/pdb], 6nwp [http://doi.org/10.2210/pdb6NWP/pdb], 6tjx [https://www.ebi.ac.uk/pdbe/entry/pdb/6TJX], 7P6D [https://www.ebi.ac.uk/pdbe/entry/pdb/7p6d], 7P65 [https://www.ebi.ac.uk/pdbe/entry/pdb/7p65], 7P66 [https://www.ebi.ac.uk/pdbe/entry/pdb/7p66], 8Q8U [https://www.ebi.ac.uk/pdbe/entry/pdb/8q8u], 8Q8V [https://www.ebi.ac.uk/pdbe/entry/pdb/8q8v], 8Q8W [https://www.ebi.ac.uk/pdbe/entry/pdb/8q8w], 8Q8X [https://www.ebi.ac.uk/pdbe/entry/pdb/8q8x], 8Q8Y [https://www.ebi.ac.uk/pdbe/entry/pdb/8q8y], 8Q8Z [https://www.ebi.ac.uk/pdbe/entry/pdb/8q8z], 8Q97 [https://www.ebi.ac.uk/pdbe/entry/pdb/8q97], 8Q98 [https://www.ebi.ac.uk/pdbe/entry/pdb/8q98], 8Q99 [https://www.ebi.ac.uk/pdbe/entry/pdb/8q99], 8Q9A [https://www.ebi.ac.uk/pdbe/entry/pdb/8q9a], 8Q9B [https://www.ebi.ac.uk/pdbe/entry/pdb/8q9b], 8Q9C [https://www.ebi.ac.uk/pdbe/entry/pdb/8q9c], 8Q9D [https://www.ebi.ac.uk/pdbe/entry/pdb/8q9d], 8Q9E [https://www.ebi.ac.uk/pdbe/entry/pdb/8q9e], 8QCR [https://www.ebi.ac.uk/pdbe/entry/pdb/8qcr], 8Q9F [https://www.ebi.ac.uk/pdbe/entry/pdb/8q9f], 8Q9G [https://www.ebi.ac.uk/pdbe/entry/pdb/8q9g], 8Q9H [https://www.ebi.ac.uk/pdbe/entry/pdb/8q9h], 8Q9I [https://www.ebi.ac.uk/pdbe/entry/pdb/8q9i], 8Q9J [https://www.ebi.ac.uk/pdbe/entry/pdb/8q9j], 8Q9K [https://www.ebi.ac.uk/pdbe/entry/pdb/8q9k], 8Q9L [https://www.ebi.ac.uk/pdbe/entry/pdb/8q9l], and 8Q90 [https://www.ebi.ac.uk/pdbe/entry/pdb/8q90].

## Code Availability

The software used to analyze and determine a cryo-EM structure of the WT and S341L_S352I tau_328-363_ fibril included CTFFIND v4.1^44^, RELION v4.0^34^ [https://www3.mrc-lmb.cam.ac.uk/relion/index.php/Main_Page], COOT v9.4^45^ [https://www2.mrc-lmb.cam.ac.uk/personal/pemsley/coot/]. All ΔΔG calculations on the cryo-EM fibril structures were performed using an adapted flex_ddg protocol^41^ available on [https://git.biohpc.swmed.edu/s184069/flex_ddg_ala_scn_runner] using Rosetta v3.12 [https://www.rosettacommons.org/].^11^ Flow cytometry data was analyzed using FlowJo v10 [https://www.flowjo.com/solutions/flowjo].

## Acknowledgements

The U.S. Army Medical Research Acquisition Activity, 808 Schreider Street, Fort Detrick MD 21702-5014 is the awarding and administering acquisition office and this work was supported by The Assistant Secretary of Defense for Health Affairs endorsed by the Department of Defense, in the amount of $1,639,837 through the Peer Reviewed Medical Research Program under Award Number HT9425-24-1-0641. Opinions, interpretations, conclusions and recommendations are those of the author(s) and are not necessarily endorsed by The Assistant Secretary of Defense for Health Affairs endorsed by the Department of Defense. In the conduct of research utilizing recombinant DNA, the investigator(s) adhered to NIH Guidelines for research involving recombinant DNA molecules. L.A.J. was additionally supported by an Effie Marie Cain Scholarship in Medical Research and a grant from the NIH-NIA (RF1AG076459). Transmission electron microscopy was performed at the UT Southwestern Electron Microscopy Core Facility, supported by the National Institutes of Health (1S10OD021685-01A1 and 1S10OD020103-01). Cryo-EM data were acquired at the Cryo-Electron Microscopy Facility (CEMF) and the Structural Biology Laboratory (SBL) at UT Southwestern, which are supported by a grant from the Cancer Prevention & Research Institute of Texas (RP220582). Computational resources were provided by the BioHPC cluster supported by the Lyda Hill Department of Bioinformatics at UT Southwestern. We thank members of the Joachimiak laboratory for helpful discussions and feedback on the manuscript.

## Author Contributions

C.M.D. and L.A.J. conceived of and directed the research as well as wrote the manuscript. In silico mutagenesis and data analysis of datasets was performed by I.M., K.J. and C.M.D. Cellular alanine incorporation experiments were performed by N.M. and C.M.D. In vitro aggregation and TEM experiments were performed by C.M.D. Cryo-EM analysis was performed by B.A.N. and C.M.D. L.S. and S.S. provided guidance on cryo-EM analysis. Seeding with designed tau cell lines was performed by S.R. and C.M.D., using tauopathy patient samples collected and characterized by C.L.W. All authors contributed to the revisions of the manuscript.

## References

1. Greenberg, S. G. & Davies, P. A preparation of Alzheimer paired helical filaments that displays distinct tau proteins by polyacrylamide gel electrophoresis. Proc. Natl. Acad. Sci. U. S. A. 87, 5827–5831 (1990).

2. Arriagada, P. V., Growdon, J. H., Hedley-Whyte, E. T. & Hyman, B. T. Neurofibrillary tangles but not senile plaques parallel duration and severity of Alzheimer’s disease. Neurology 42, 631–639 (1992).

3. Braak, H. & Braak, E. Staging of Alzheimer’s disease-related neurofibrillary changes. Neurobiol. Aging 16, 271–278; discussion 278-284 (1995).

4. Holmes, B. B., et al. Proteopathic tau seeding predicts tauopathy in vivo. Proc. Natl. Acad. Sci. 111, E4376–E4385 (2014).

5. Fitzpatrick, A. W. P. et al. Cryo-EM structures of tau filaments from Alzheimer’s disease. Nature 547, 185–190 (2017).

6. Falcon, B. et al. Structures of filaments from Pick’s disease reveal a novel tau protein fold. Nature 561, 137–140 (2018).

7. Shi, Y. et al. Structure-based classification of tauopathies. Nature 598, 359–363 (2021).

8. Falcon, B. et al. Novel tau filament fold in chronic traumatic encephalopathy encloses hydrophobic molecules. Nature 568, 420–423 (2019).

9. Zhang, W. et al. Novel tau filament fold in corticobasal degeneration. Nature 1–5 (2020).

10. van der Kant, R., Louros, N., Schymkowitz, J. & Rousseau, F. Thermodynamic analysis of amyloid fibril structures reveals a common framework for stability in amyloid polymorphs. Struct. Lond. Engl. 1993 30, 1178–1189.e3 (2022).

11. Mullapudi, V. et al. Network of hotspot interactions cluster tau amyloid folds. Nat. Commun. 14, 895–19 (2023).

12. Louros, N. et al. Local structural preferences in shaping tau amyloid polymorphism. Nat. Commun. 15, 1028 (2024).

13. Lövestam, S. et al. Twelve phosphomimetic mutations induce the assembly of recombinant full-length human tau into paired helical filaments. eLife 14, RP104778 (2026).

14. Lövestam, S. et al. Assembly of recombinant tau into filaments identical to those of Alzheimer’s disease and chronic traumatic encephalopathy. bioRxiv 2021.12.16.472950 (2021).

15. Vaquer-Alicea, J. et al. Functional classification of tauopathy strains reveals the role of protofilament core residues. Sci. Adv. 11, eadp5978 (2025).

16. Santambrogio, A. et al. Serial amplification of tau filaments using Alzheimer’s brain homogenates and C322A or C322S recombinant tau. FEBS Lett. 599, 2768–2778 (2025).

17. Tarutani, A. et al. Cryo-EM structures of tau filaments from SH-SY5Y cells seeded with brain extracts from cases of Alzheimer’s disease and corticobasal degeneration. FEBS Open Bio 13, 1394–1404 (2023).

18. Jumper, J. et al. Highly accurate protein structure prediction with AlphaFold. Nature 596, 583–589 (2021).

19. Dauparas, J. et al. Robust deep learning-based protein sequence design using ProteinMPNN. Science 378, 49–56 (2022).

20. Pinheiro, F., Santos, J. & Ventura, S. AlphaFold and the amyloid landscape. J. Mol. Biol. 433, 167059 (2021).

21. Wilkinson, M. et al. Structural evolution of fibril polymorphs during amyloid assembly. Cell 186, 5798–5811.e26 (2023).

22. Lövestam, S. et al. Disease-specific tau filaments assemble via polymorphic intermediates. Nature 625, 119–125 (2024).

23. Chen, D. et al. Tau local structure shields an amyloid-forming motif and controls aggregation propensity. Nat. Commun. 10, 2493 (2019).

24. Karamanos, T. K., Kalverda, A. P. & Radford, S. E. Generating Ensembles of Dynamic Misfolding Proteins. Front. Neurosci. 16, 881534 (2022).

25. Raskatov, J. A. & Teplow, D. B. Using chirality to probe the conformational dynamics and assembly of intrinsically disordered amyloid proteins. Sci. Rep. 7, 12433 (2017).

26. Gadhe, L. et al. Reverse-engineering amyloid strains with generative protein design. Preprint at 10.64898/2026.05.08.723915 (2026).

27. Schweers, O., Mandelkow, E. M., Biernat, J. & Mandelkow, E. Oxidation of cysteine-322 in the repeat domain of microtubule-associated protein tau controls the in vitro assembly of paired helical filaments. Proc. Natl. Acad. Sci. U. S. A. 92, 8463–8467 (1995).

28. Prifti, E. et al. The Two Cysteines of Tau Protein Are Functionally Distinct and Contribute Differentially to Its Pathogenicity in Vivo. J. Neurosci. Off. J. Soc. Neurosci. 41, 797–810 (2021).

29. Jayan, P. et al. Cysteines are critical determinants of spontaneous and seeded tau aggregation in cells. Preprint at 10.64898/2026.03.12.711133 (2026).

30. Weismiller, H. A., Holub, T. J., Krzesinski, B. J. & Margittai, M. A thiol-based intramolecular redox switch in four-repeat tau controls fibril assembly and disassembly. J. Biol. Chem. 297, 101021 (2021).

31. Rocklin, G. J. et al. Global analysis of protein folding using massively parallel design, synthesis, and testing. Science 357, 168–175 (2017).

32. Tsuboyama, K. et al. Mega-scale experimental analysis of protein folding stability in biology and design. Nature 620, 434–444 (2023).

33. Ferrari, Á. J. R. et al. Large-scale discovery, analysis and design of protein energy landscapes. Nature 654, 1108–1118 (2026).

34. Scheres, S. H. W. Amyloid structure determination in RELION-3.1. Acta Crystallogr. Sect. Struct. Biol. 76, 94–101 (2020).

35. Shekhar, M. et al. CryoFold: determining protein structures and data-guided ensembles from cryo-EM density maps. Matter 4, 3195–3216 (2021).

36. Adams, P. D. et al. PHENIX: a comprehensive Python-based system for macromolecular structure solution. Acta Crystallogr. D Biol. Crystallogr. 66, 213–221 (2010).

37. Thompson, M. J. et al. The 3D profile method for identifying fibril-forming segments of proteins. Proc. Natl. Acad. Sci. 103, 4074–4078 (2006).

38. Sawaya, M. R. et al. Atomic structures of amyloid cross-beta spines reveal varied steric zippers. Nature 447, 453–457 (2007).

39. Hitt, B. D. et al. Ultrasensitive tau biosensor cells detect no seeding in Alzheimer’s disease CSF. Acta Neuropathol. Commun. 9, 99 (2021).

40. Sawaya, M. R. et al. Atomic structures of amyloid cross-β spines reveal varied steric zippers. Nature 447, 453–457 (2007).

41. Barlow, K. A. et al. Flex ddG: Rosetta Ensemble-Based Estimation of Changes in Protein-Protein Binding Affinity upon Mutation. J. Phys. Chem. B 122, 5389–5399 (2018).

42. Virtanen, P. et al. SciPy 1.0: fundamental algorithms for scientific computing in Python. Nat. Methods 17, 261–272 (2020).

43. Meisl, G. et al. Molecular mechanisms of protein aggregation from global fitting of kinetic models. Nat. Protoc. 11, 252–272 (2016).

44. Rohou, A. & Grigorieff, N. CTFFIND4: Fast and accurate defocus estimation from electron micrographs. J. Struct. Biol. 192, 216–221 (2015).

45. Emsley, P. & Cowtan, K. Coot: model-building tools for molecular graphics. Acta Crystallogr. D Biol. Crystallogr. 60, 2126–2132 (2004).

