## Supplementary Information for "Large-Scale Mutagenesis Identifies Sequence Changes That Bias Tau Toward CTE-like Aggregation"

**Supplementary Tables and Figures**

| **Construct** | **Buffer** | **t_1/2_ ± Std (p-value)** |
| --- | --- | --- |
| WT tau_328-363_ | No salt | 1.464 ± 0.135 |
|  | NaCl | 2.177 ± 0.250 |
|  | MgCl_2_ | 4.014 ± 1.505 |
| S341A tau_328-363_ | No salt | 2.294 ± 0.287 |
|  | NaCl | 1.094 ± 0.046 |
|  | MgCl_2_ | 1.527 ± 0.583 |
| S352A tau_328-363_ | No salt | 2.082 ± 0.507 |
|  | NaCl | 1.669 ± 0.411 |
|  | MgCl_2_ | 0.922 ± 0.090 |
| S356A tau_328-363_ | No salt | -- |
|  | NaCl | 0.658 ± 0.075 |
|  | MgCl_2_ | 0.507 ± 0.075 |
| L357A tau_328-363_ | No salt | -- |
|  | NaCl | -- |
|  | MgCl_2_ | -- |
| S341A_S352A tau_328-363_ | No salt | 1.840 ± 0.097 |
|  | NaCl | 1.134 ± 0.268 |
|  | MgCl_2_ | 0.327 ± 0.034 |
| S357F tau_328-363_ | No salt | 4.280 ± 0.198 |
|  | NaCl | 1.245 ± 0.114 |
|  | MgCl_2_ | 0.156 ± 0.017 |
| S341A_S357F tau_328-363_ | No salt | 0.307 ± 0.017 |
|  | NaCl | 0.176 ± 0.017 |
|  | MgCl_2_ | 0.146 ± 0.017 |
| S352A_S357F tau_328-363_ | No salt | 0.584 ± 0.061 |
|  | NaCl | 1.366 ± 0.087 |
|  | MgCl_2_ | 0.408 ± 0.188 |
| S341A_S352A_L357F tau_328-363_ | No salt | 0.166 ± 0 (0.0001) |
|  | NaCl | 0.105 ± 0 (0.0001) |
|  | MgCl_2_ | 0.136 ± 0.030 (0.011) |
| S341L tau_328-363_ | No salt | 0.869 ± 0.017 |
|  | NaCl | 0.839 ± 0.034 |
|  | MgCl_2_ | 0.658 ± 0.092 |
| S352I tau_328-363_ | No salt | 4.793 ± 0.060 |
|  | NaCl | 0.288 ± 0.033 |
|  | MgCl_2_ | 0.031 ± 0.010 |
| S341L_S352I tau_328-363_ | No salt | 0.048 ± 0.006 (0.003) |
|  | NaCl | 0.045 ± 0.024 (0.004) |
|  | MgCl_2_ | 0.024 ± 0.006 (0.044) |
| S356L_L357S tau_328-363_ | No salt | 3.467 ± 0.188 |
|  | NaCl | 3.869 ± 0.448 |
|  | MgCl_2_ | 2.502 ± 0.707 |
| **Construct** | **Buffer** | **t_1/2_ ± Std (p-value)** |
| WT_328-363_ | No salt | 1.46 ± 0.136 |
|  | NaCl | 2.17 ± 0.250 |
|  | MgCl_2_ | 4.01 ± 1.500 |
| S341A_328-363_ | No salt | 2.29 ± 0.287 |
|  | NaCl | 1.09 ± 0.046 |
|  | MgCl_2_ | 1.52 ± 0.58 |
| S352A_328-363_ | No salt | 2.08 ± 0.50 |
|  | NaCl | 1.67 ± 0.41 |
|  | MgCl_2_ | 0.92 ± 0.09 |
| S356A_328-363_ | No salt | -- |
|  | NaCl | 0.66 ± 0.076 |
|  | MgCl_2_ | 0.51 ± 0.076 |
| L357A_328-363_ | No salt | -- |
|  | NaCl | -- |
|  | MgCl_2_ | -- |
| S341A_S352A_328-363_ | No salt | 1.84 ± 0.097 |
|  | NaCl | 1.13 ± 0.268906 |
|  | MgCl_2_ | 0.32 ± 0.034935 |
| S357F_328-363_ | No salt | 4.280 ± 0.198 |
|  | NaCl | 1.245 ± 0.114 |
|  | MgCl_2_ | 0.156 ± 0.017 |
| S341A_S357F _328-363_ | No salt | 0.307 ± 0.017 |
|  | NaCl | 0.176 ± 0.017 |
|  | MgCl_2_ | 0.146 ± 0.017 |
| S352A_S357F_328-363_ | No salt | 0.584 ± 0.061 |
|  | NaCl | 1.366 ± 0.087 |
|  | MgCl_2_ | 0.408 ± 0.188 |
| S341A_S352A_L357F_328-363_ | No salt | 0.166 ± 0 (0.0001) |
|  | NaCl | 0.105 ± 0 (0.0001) |
|  | MgCl_2_ | 0.136 ± 0.030 (0.011) |
| S341L_328-363_ | No salt | 0.869 ± 0.017 |
|  | NaCl | 0.839 ± 0.034 |
|  | MgCl_2_ | 0.658 ± 0.092 |
| S352I_328-363_ | No salt | 4.793 ± 0.060 |
|  | NaCl | 0.288 ± 0.033 |
|  | MgCl_2_ | 0.031 ± 0.010 |
| S341L_S352I_328-363_ | No salt | 0.048 ± 0.006 (0.003) |
|  | NaCl | 0.045 ± 0.024 (0.004) |
|  | MgCl_2_ | 0.024 ± 0.006 (0.044) |
| S356L_L357S_328-363_ | No salt | 3.467 ± 0.188 |
|  | NaCl | 3.869 ± 0.448 |
|  | MgCl_2_ | 2.502 ± 0.707 |

**Supplementary Table 1.** Thioflavin T aggregation kinetics for tau constructs.

| Construct | 328-363 wt |
| --- | --- |
|  | No salt |
| **Data collection and processing** |  |
| Magnification | 130,000 |
| Voltage (kV) | 300 |
| Detector | Falcon4 |
| Energy filter | Selectris |
| Electron exposure (e^-^/Å^2^) | 50 |
| Exposure rate (e^-^/pixel/s) | 8.604 |
| Nominal defocus range (mm) | -2.2 to -0.8 (15 steps) |
| Pixel size (Å) | 0.936 |
| Movies collected | 4,296 |
| Initial particle images (no.) | 1,127,246 |
| Final particle images (no.) | 140,439 |
| Map resolution (Å) | 3.6 |
| FSC threshold | 0.143 |
| Map resolution range (Å) | 3.4 – 6.6 |
| Symmetry imposed | hss |
| Helical parameters |  |
| Helical twist (°) | -1.84 (178.16) |
| Helical rise (Å) | 4.73 |
| Crossover distance (nm) | 463 |
| **Refinement** |  |
| Map sharpening *B* factor (Å^2^) | -30.5497 |
| Cross correlation (masked) | 0.82 |
| Model resolution | 3.456 |
| FSC threshold | 0.143 |
| Model composition |  |
| Non-hydrogen atoms | 3320 |
| Protein residues total | 420 |
| Ligand/solvent molecules | 0 |
| Chains per helical layer | 4 |
| Helical layers modelled | 5 |
| *B* factors (Å^2^) |  |
| Protein | 76.18 |
| Ligand/solvent | -- |
| R.m.s deviations |  |
| Bond lengths (Å) | 0.004 |
| Bond angles (°) | 1.131 |
| Validation |  |
| MolProbity score | 1.12 |
| Clashscore | 3.25 |
| Poor rotamers (%) | 0.00 |
| Ramachandran plot |  |
| Favored (%) | 100.00 |
| Allowed (%) | 0.00 |
| Disallowed (%) | 0.00 |

**Supplementary Table 2.** Data collection and analysis parameters for WT tau_328-363._

| Construct | 328-363 S341L S352I | | |
| --- | --- | --- | --- |
| Buffer conditions | No salt | 200 mM NaCl | 200 mM MgCl_2_ |
| **Data collection and processing** |  |  |  |
| Magnification | 105,000 | 130,000 | 130,000 |
| Voltage (kV) | 300 | 300 | 300 |
| Detector | Falcon4 | Falcon4 | Falcon4 |
| Energy filter | Selectris | Selectris | Selectris |
| Electron exposure (e^-^/Å^2^) | 50 | 50 | 50 |
| Exposure rate (e^-^/pixel/s) | 8.697 | 9.033 | 7.916 |
| Nominal defocus range (mm) | -2.2 to -0.8 (14 steps) | -2.2 to -0.8 (14 steps) | -2.2 to -0.8 (14 steps) |
| Pixel size (Å) | 0.834 | 0.936 | 0.936 |
| Movies collected | 6,294 | 7,019 | 6,768 |
| Initial particle images (no.) | 5,304,411 | 1,139,747 | 2,925,771 |
| Final particle images (no.) |  |  | 152,763 |
| Map resolution (Å) |  |  | 2.74 |
| FSC threshold |  |  | 0.143 |
| Map resolution range (Å) |  |  | 2.5 – 5.0 |
| Symmetry imposed | hss | hss | hss |
| Helical parameters |  |  |  |
| Helical twist (°) |  |  | -1.285 (178.715) |
| Helical rise (Å) |  |  | 2.37 |
| Crossover distance (nm) |  |  | 332.7 |
| **Refinement** |  |  |  |
| Map sharpening *B* factor (Å^2^) |  |  | -64.0196 |
| Cross correlation (masked) |  |  | 0.74 |
| Model resolution |  |  | 2.2 |
| FSC threshold |  |  | 0.143 |
| Model composition |  |  |  |
| Non-hydrogen atoms |  |  | 472 |
| Protein residues total |  |  | 60 |
| Ligand/solvent molecules |  |  | 0 |
| Chains per helical layer |  |  | 4 |
| Helical layers modelled |  |  | 1 |
| *B* factors (Å^2^) |  |  |  |
| Protein |  |  | 32.57 |
| Ligand/solvent |  |  | -- |
| R.m.s deviations |  |  |  |
| Bond lengths (Å) |  |  | 0.008 |
| Bond angles (°) |  |  | 0.729 |
| Validation |  |  |  |
| MolProbity score |  |  | 2.02 |
| Clashscore |  |  | 6.12 |
| Poor rotamers (%) |  |  | 3.57 |
| Ramachandran plot |  |  |  |
| Favored (%) |  |  | 96.15 |
| Allowed (%) |  |  | 3.85 |
| Disallowed (%) |  |  | 0 |

**Supplementary Table 3.** Data collection and analysis parameters for S341L/S352I tau_328-363_ in different buffer conditions.

| **Construct** | **Buffer** | **t_1/2_ ± Std (p-value)** |
| --- | --- | --- |
| 297-391 wt | No salt | **--** |
|  | NaCl | 3.09 ± 0.63 |
|  | MgCl_2_ | 4.10 ± 0.43 |
| 297-391 S341L_L352I | No salt | 0.97 ± 0.066 |
|  | NaCl | 1.67 ± 0.017 |
|  | MgCl_2_ | 4.53 ± 0.918 |

**Supplementary Table 4.** Thioflavin T aggregation kinetics for tau_297-391_ constructs.

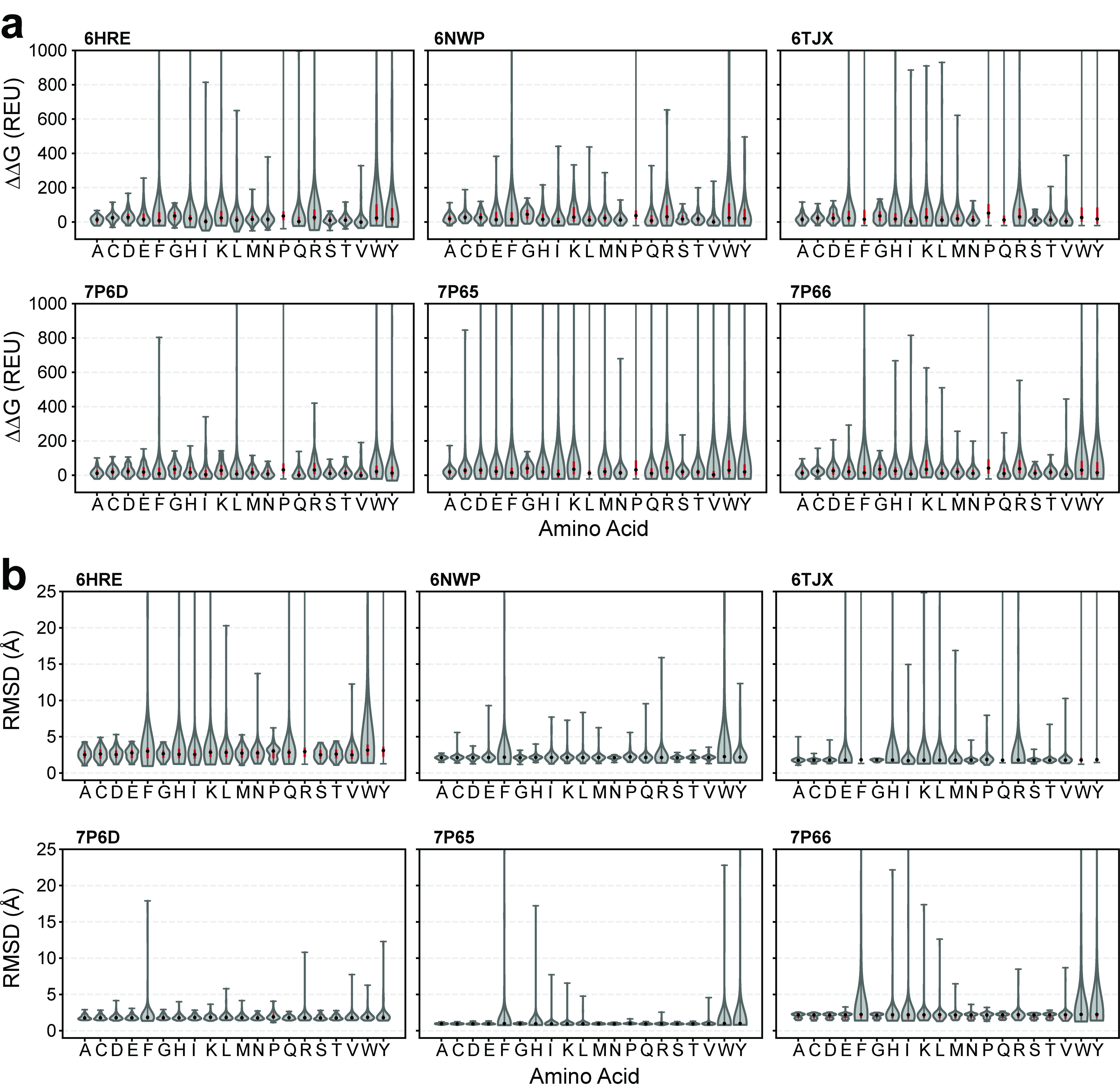

**Supplementary Figure 1. Distribution of in silico saturation mutagenesis energetics for tauopathy protofilament folds.** a. Violin plots illustrating the energetic distribution (DDG) across mutation types for each of the tauopathy folds. Bulky or highly restrained residue types are observed to have higher, more unfavorable, energetic distributions. b. Violin plots showing the RMSD (Å) distributions across mutation types for each of the tauopathy folds. The data supports that larger residues tend to have higher distributions. This data suggests that bulkier residues are most likely to cause unfavorable interactions within the tauopathy folds. Red bars indicate the interquartile range; medians are indicated by the central mark.

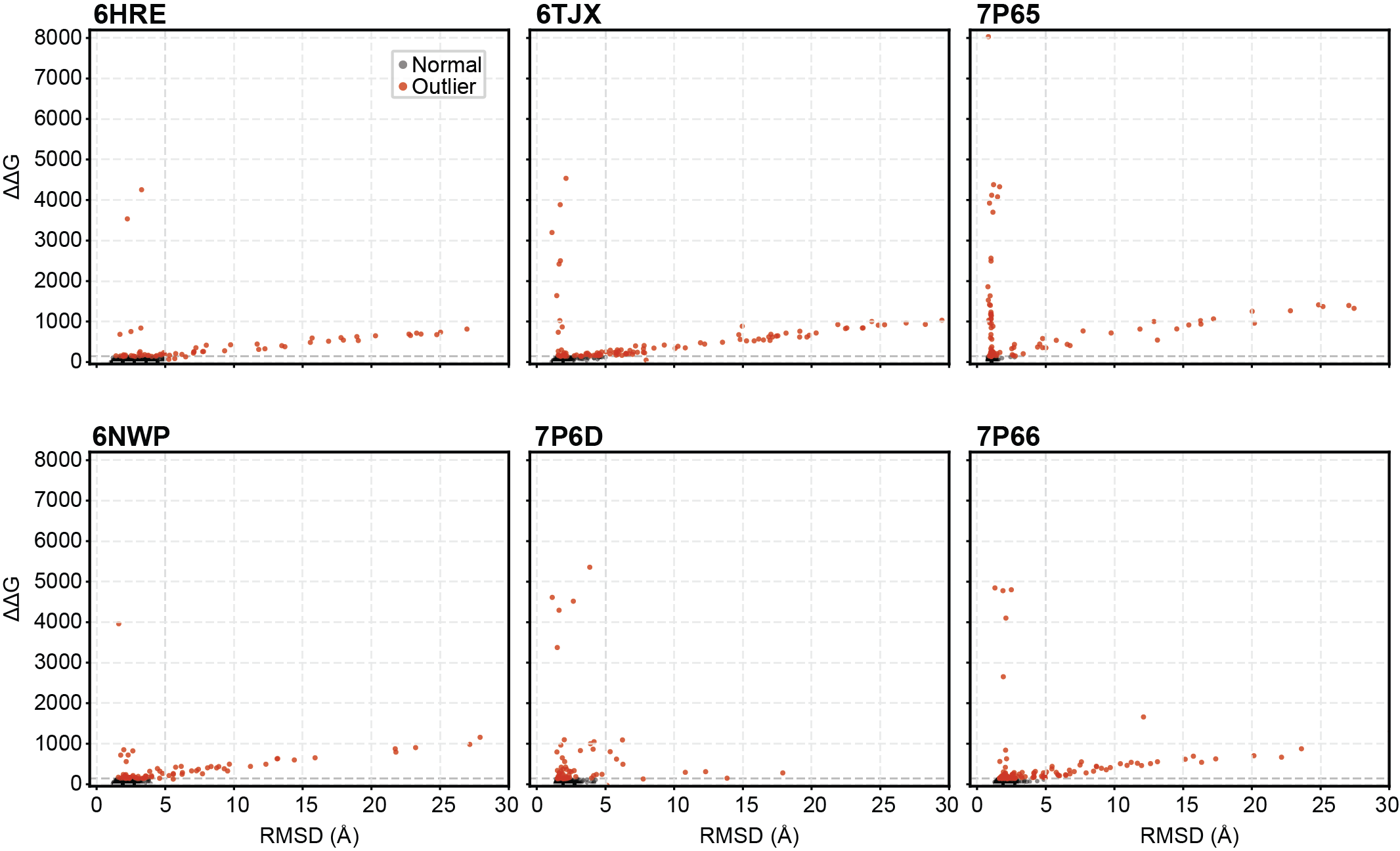

**Supplementary Figure 2. Identifying outliers in the in silico saturation mutagenesis datasets.** Scatter plots comparing the energetic and structural influence of each mutation for each taupathy fold. Outliers were determined based on energetic and RMSD cutoffs of >138 units and 5Å respectively. These cutoff values retained > 93% of each dataset.

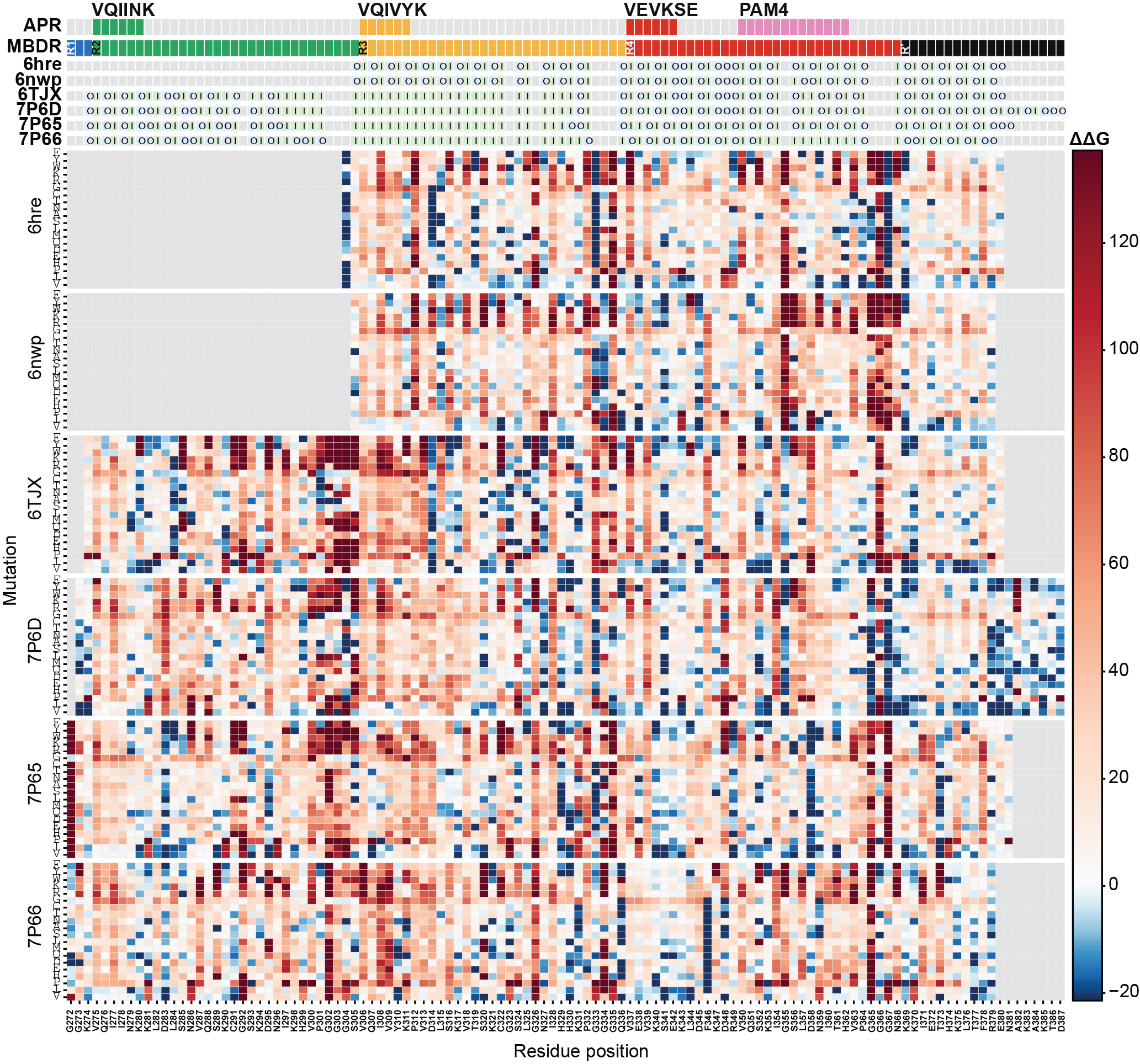

**Supplementary Figure 3. In silico saturation mutagenesis for tauopathy protofilament folds.** Heat map showing predicted energetic effects of all amino acid substitutions across the ordered tau amyloid core 272-387 for AD (PDB ID 6HRE), CTE (PDB ID 6NWP), CBD (PDB ID 6TJX), AGD (PDB ID 7P6D), PSP (PDB ID 7P65) and GGT (PDB ID 7P66) tau fibril templates. Columns correspond to residue positions in tau and rows correspond to amino acid substitutions grouped by fibril structure. The upper annotations indicate aggregation-prone motifs VQIINK (green), VQIVYK (orange), VEVKSE (red) and PAM4 (pink), microtubule-binding domain repeat (MBDR) organization (R1 in blue, R2 in green, R3 (orange), R4 (red) and R4’(black)) and whether each residue is buried inside the fibril (I) or outside and solvent accessible (O) across the analyzed fibril templates. The color scale reports predicted ΔΔG values, with red indicating substitutions predicted to be destabilizing and blue indicating substitutions predicted to be stabilizing or more tolerated. Gray regions indicate residues not resolved or not evaluated in the corresponding fibril template.

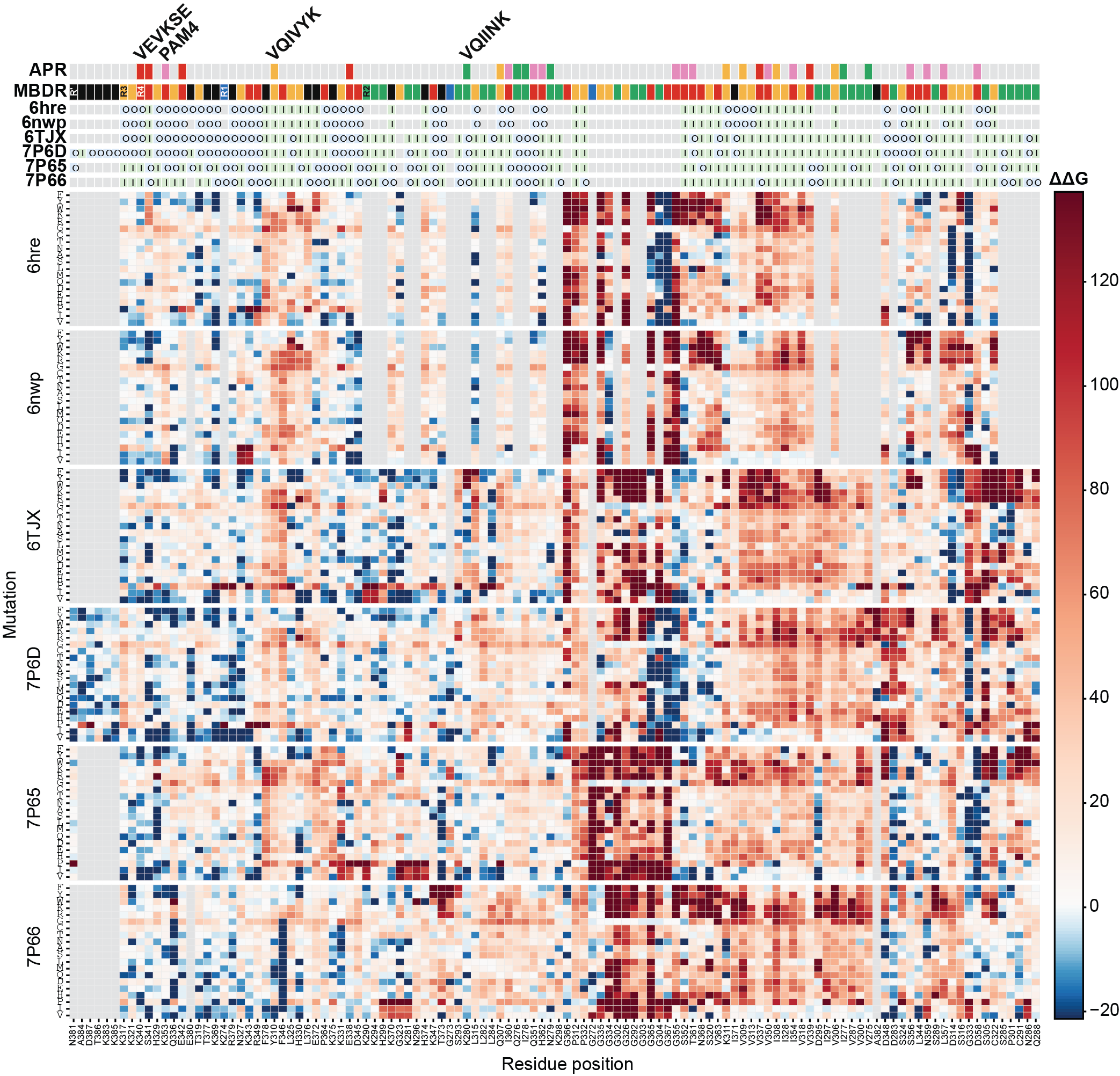

**Supplementary Figure 4. Hierarchical clustering of in silico saturation mutagenesis ΔΔG for tauopathy protofilament folds.** Heat map showing predicted energetic effects of all amino acid substitutions across the ordered tau amyloid core for six disease-associated tau fibril templates: AD (PDB ID 6HRE), CTE (PDB ID 6NWP), CBD (PDB ID 6TJX), AGD (PDB ID 7P6D), PSP (PDB ID 7P65) and GGT (PDB ID 7P66). Columns correspond to tau residue positions clustered across all six structures based on similarity in their mutational energy profiles and rows correspond to amino acid substitutions within each fibril template. This position-based clustering groups residues with shared energetic behavior across folds and highlights positions with fold-selective mutational sensitivity. Upper annotations for amyloid motifs, repeat and burial of amino acid are as described in Supplementary Figure 3. The color scale reports predicted ΔΔG values, with red indicating substitutions predicted to be destabilizing and blue indicating substitutions predicted to be stabilizing or more tolerated. Gray regions indicate residues that were not resolved or not evaluated in the corresponding fibril template.

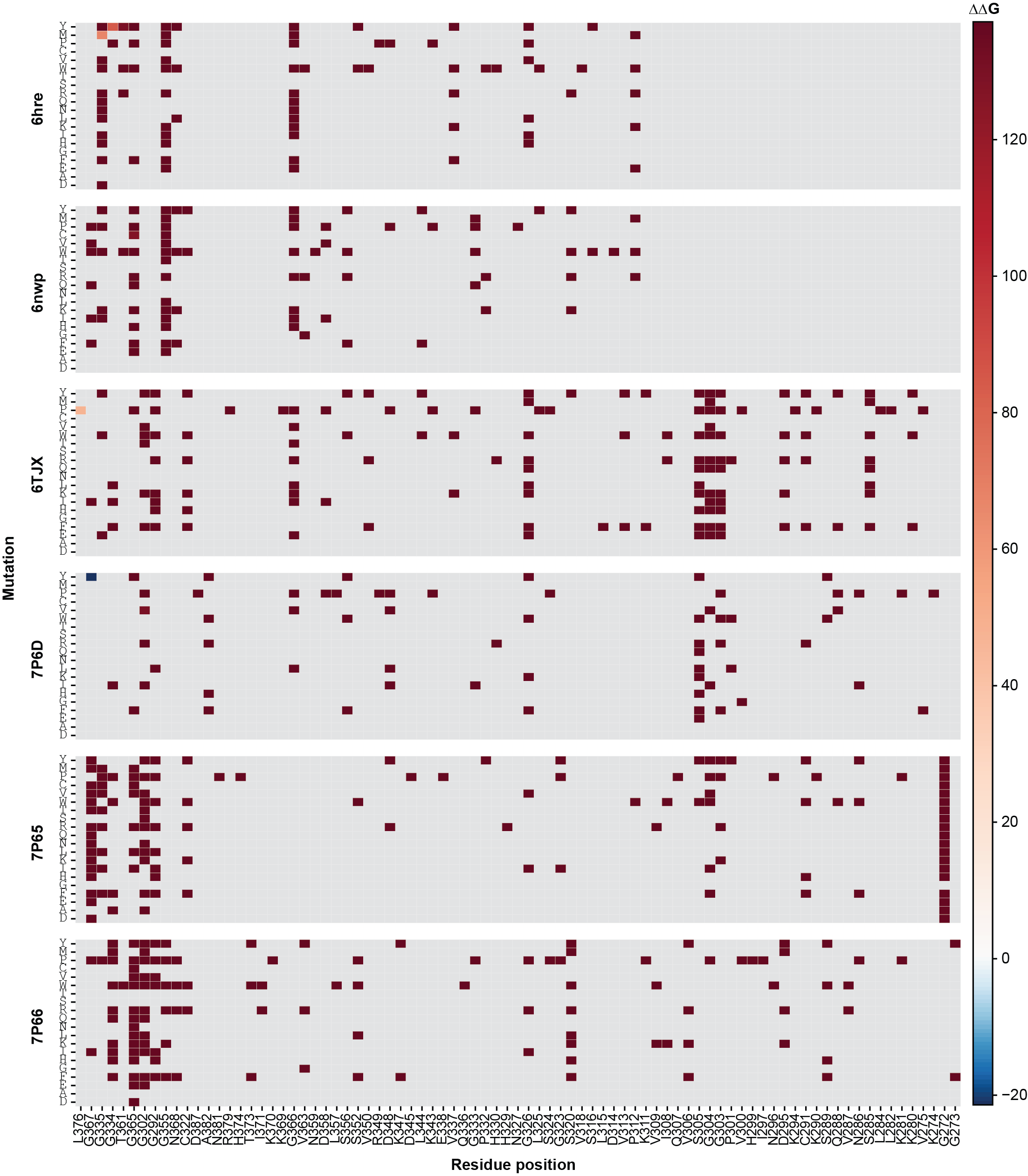
**Supplementary Figure 5. Outliers within tauopathy saturation mutagenesis datasets.** Heat map showing predicted energetic effects of all amino acid substitutions across the ordered tau amyloid core for AD (PDB ID 6HRE), CTE (PDB ID 6NWP), CBD (PDB ID 6TJX), AGD (PDB ID 7P6D), PSP (PDB ID 7P65) and GGT (PDB ID 7P66) tau fibril templates. Columns correspond to residue positions in tau and rows correspond to amino acid substitutions grouped by fibril structure. The color scale reports predicted ΔΔG values, with red indicating substitutions predicted to be destabilizing and blue indicating substitutions predicted to be stabilizing or more tolerated. Gray regions indicate residues that were not resolved or not evaluated in the corresponding fibril template. Outliers were determined based on energetic and RMSD cutoffs of >138 units and 5Å respectively. These outliers largely map to broadly constrained regions within the tauopathy folds – the native residues most often affected are small, flexible positions (i.e., G). Mutation outliers are mostly bulky, aromatic, proline, or charged substitutions introduced at constrained small residues. The list of outliers is not composed of structurally selective mutations but rather mutations that break packing, backbone geometry, or conserved structural constraints.

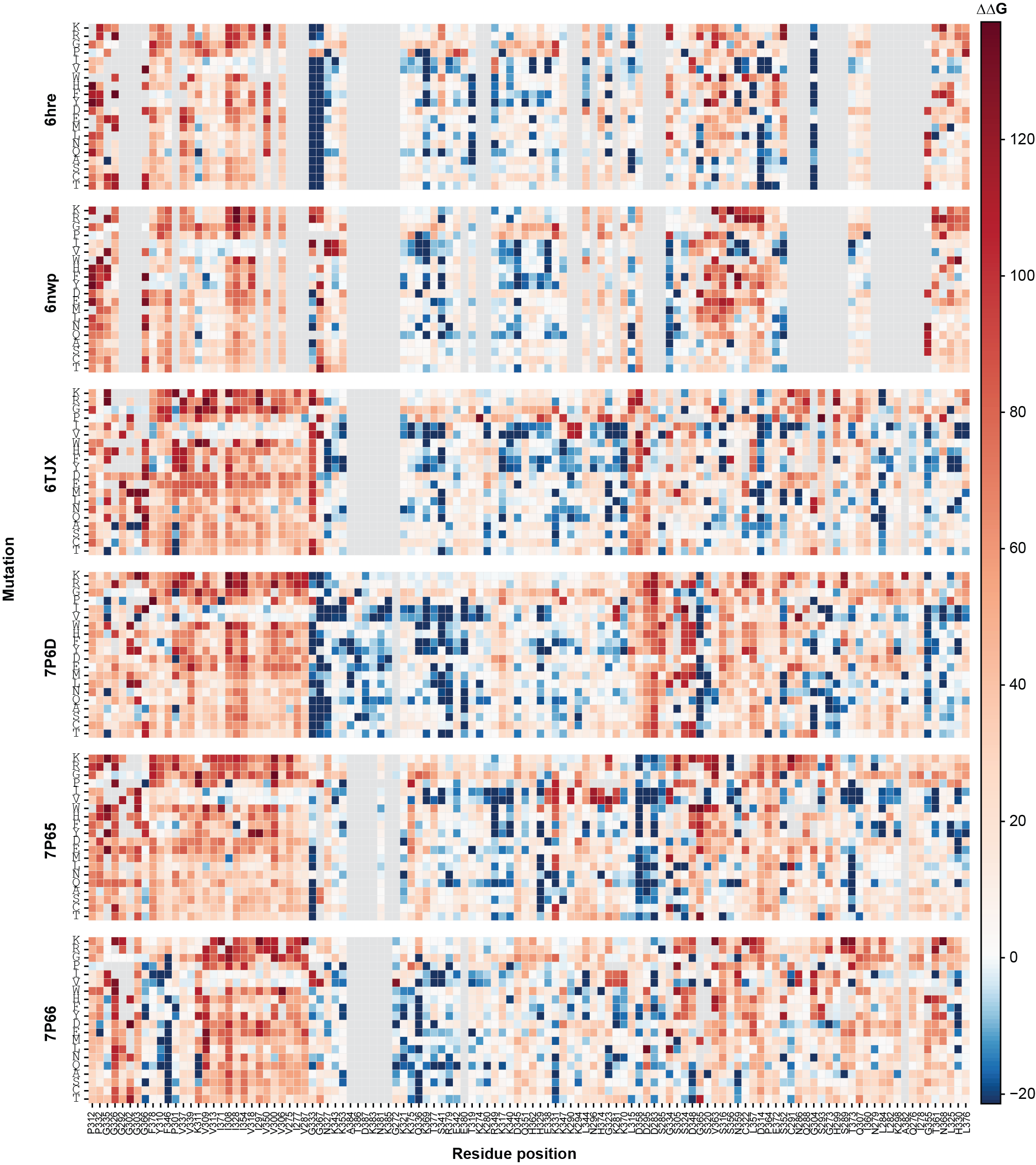

**Supplementary Figure 6. Removing outliers visually alters clustering of in silico saturation mutagenesis but does not alter selective mutations for tauopathy protofilament folds.** Heat map showing predicted energetic effects of all amino acid substitutions across the ordered tau amyloid core for AD (PDB ID 6HRE), CTE (PDB ID 6NWP), CBD (PDB ID 6TJX), AGD (PDB ID 7P6D), PSP (PDB ID 7P65) and GGT (PDB ID 7P66) tau fibril templates. Columns correspond to residue positions in tau and rows correspond to amino acid substitutions grouped by fibril structure. The color scale reports predicted ΔΔG values, with red indicating substitutions predicted to be destabilizing and blue indicating substitutions predicted to be stabilizing or more tolerated. Gray regions indicate residues that were not resolved or not evaluated in the corresponding fibril template. Outliers were determined based on energetic and RMSD cutoffs of >138 units and 5 Å respectively. Removing outliers alters clustering topology, indicating that extreme destabilizing/disruptive mutations influence the visual ordering. However, removal of these outliers does not create new selective patterns and the main structure-selective conclusions remain.

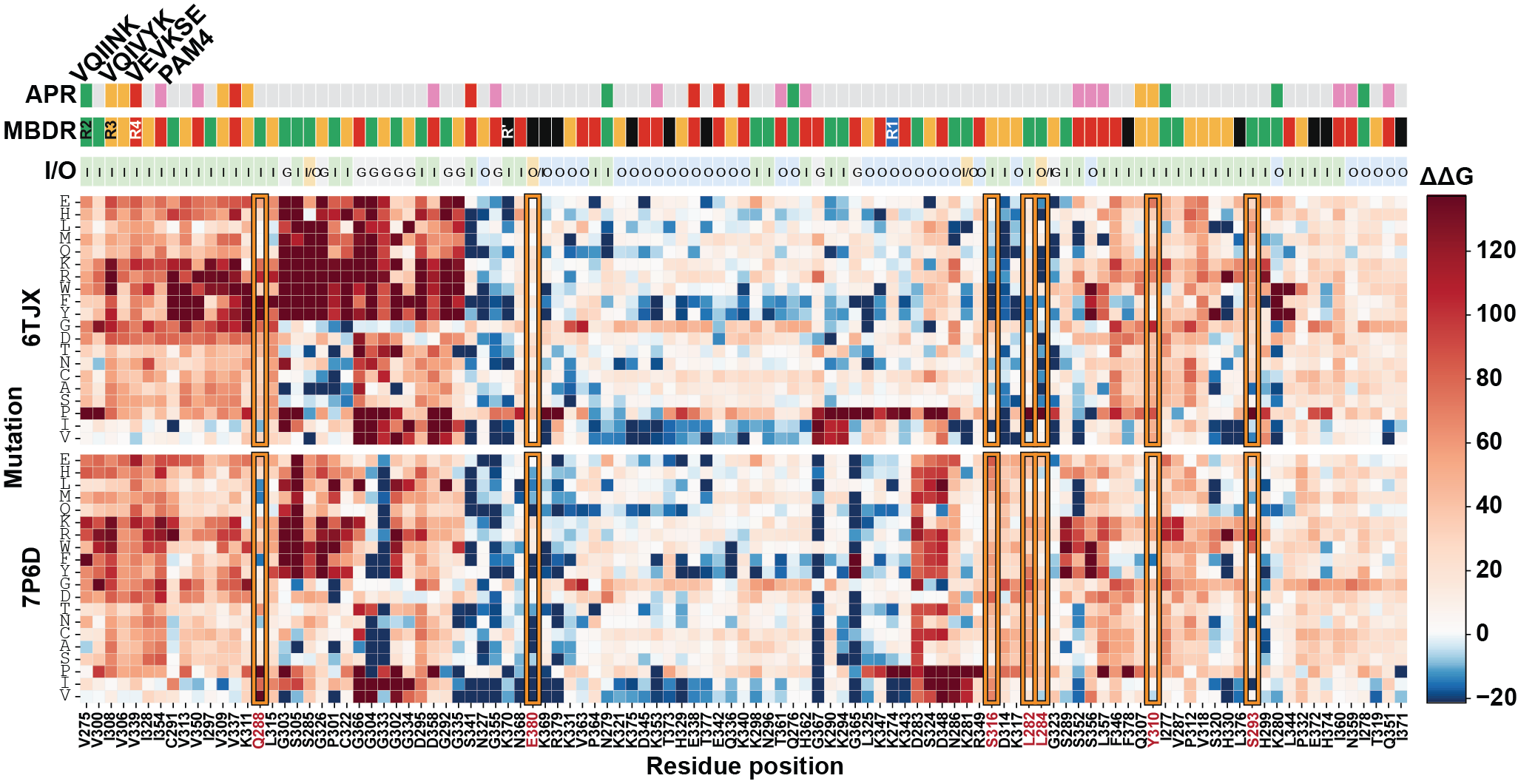

**Supplementary Figure 7. Clustered saturation mutagenesis ΔΔG heatmap for CBD and AGD tauopathy fibril structures.** Heat map showing predicted energetic effects of all amino acid substitutions across the ordered tau amyloid core for CBD (PDB ID 6TJX) and AGD (PDB ID 7P6D) tau fibril templates. Columns correspond to tau residue positions clustered across the two structures based on similarity in their mutational energy profiles and rows correspond to amino acid substitutions within each fibril template. This position-based hierarchical clustering highlights residues with shared energetic behavior between CBD and AGD, as well as positions with fold-selective mutational sensitivity. Upper annotations for amyloid motifs, repeat and burial of amino acid are as described in Supplementary Figure 3. The color scale reports predicted ΔΔG values, with red indicating substitutions predicted to be destabilizing and blue indicating substitutions predicted to be stabilizing or more tolerated. Gray regions indicate residues that were not resolved or not evaluated in the corresponding fibril template. Boxed regions highlight positions with strong or differential mutational sensitivity between CBD and AGD.

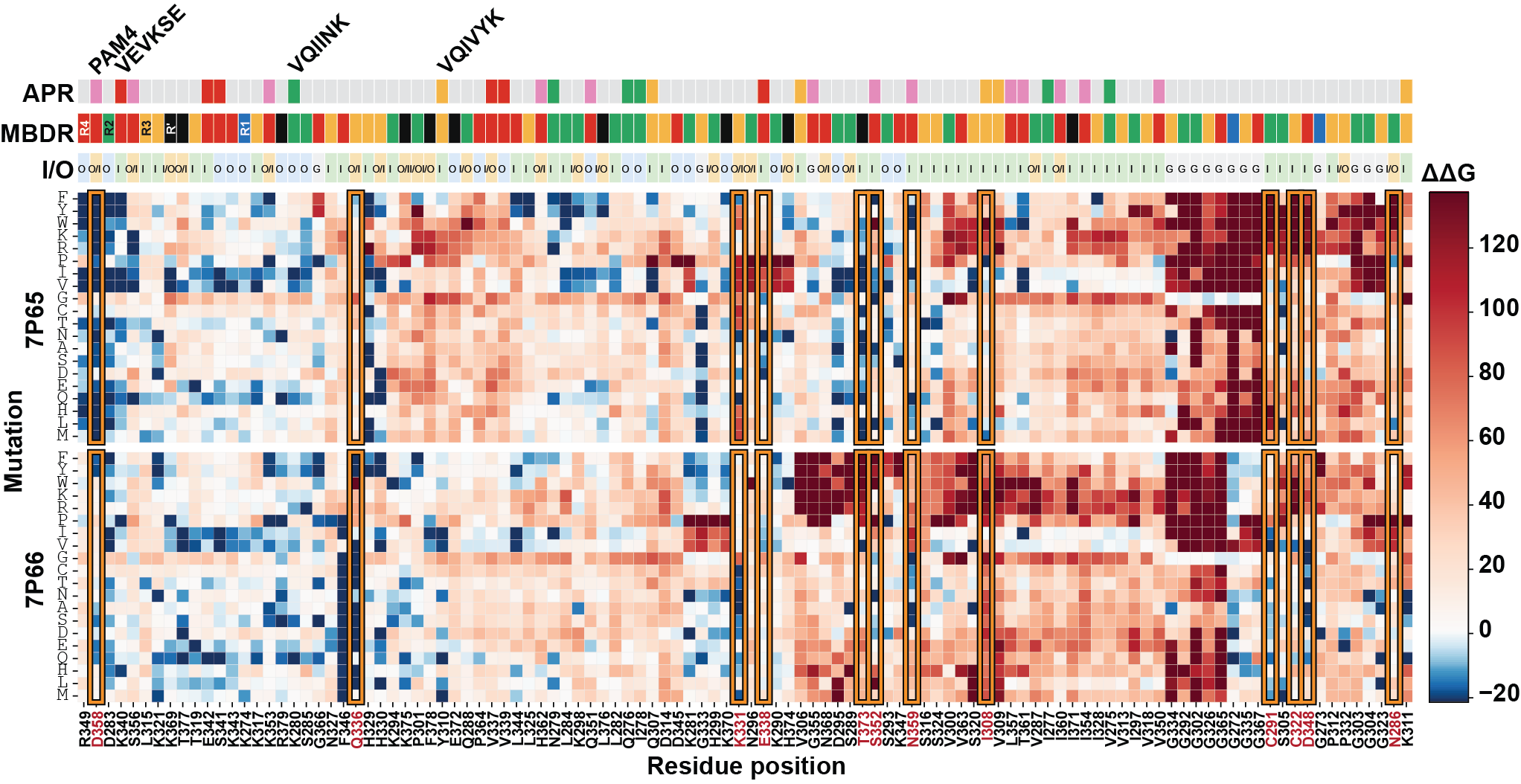

**Supplementary Figure 8. Clustered saturation mutagenesis ΔΔG heatmap for PSP and GGT tauopathy fibril structures.** Heat map showing predicted energetic effects of all amino acid substitutions across the ordered tau amyloid core for PSP (PDB ID 7P65) and GGT (PDB ID 7P66) tau fibril templates. Columns correspond to tau residue positions clustered across the two structures based on similarity in their mutational energy profiles and rows correspond to amino acid substitutions within each fibril template. This position-based hierarchical clustering highlights residues with shared energetic behavior between PSP and GGT, as well as positions with fold-selective mutational sensitivity. Upper annotations for amyloid motifs, repeat and burial of amino acid are as described in Supplementary Figure 3. The color scale reports predicted ΔΔG values, with red indicating substitutions predicted to be destabilizing and blue indicating substitutions predicted to be stabilizing or more tolerated. Gray regions indicate residues that were not resolved or not evaluated in the corresponding fibril template. Boxed regions highlight positions with strong or differential mutational sensitivity between PSP and GGT.

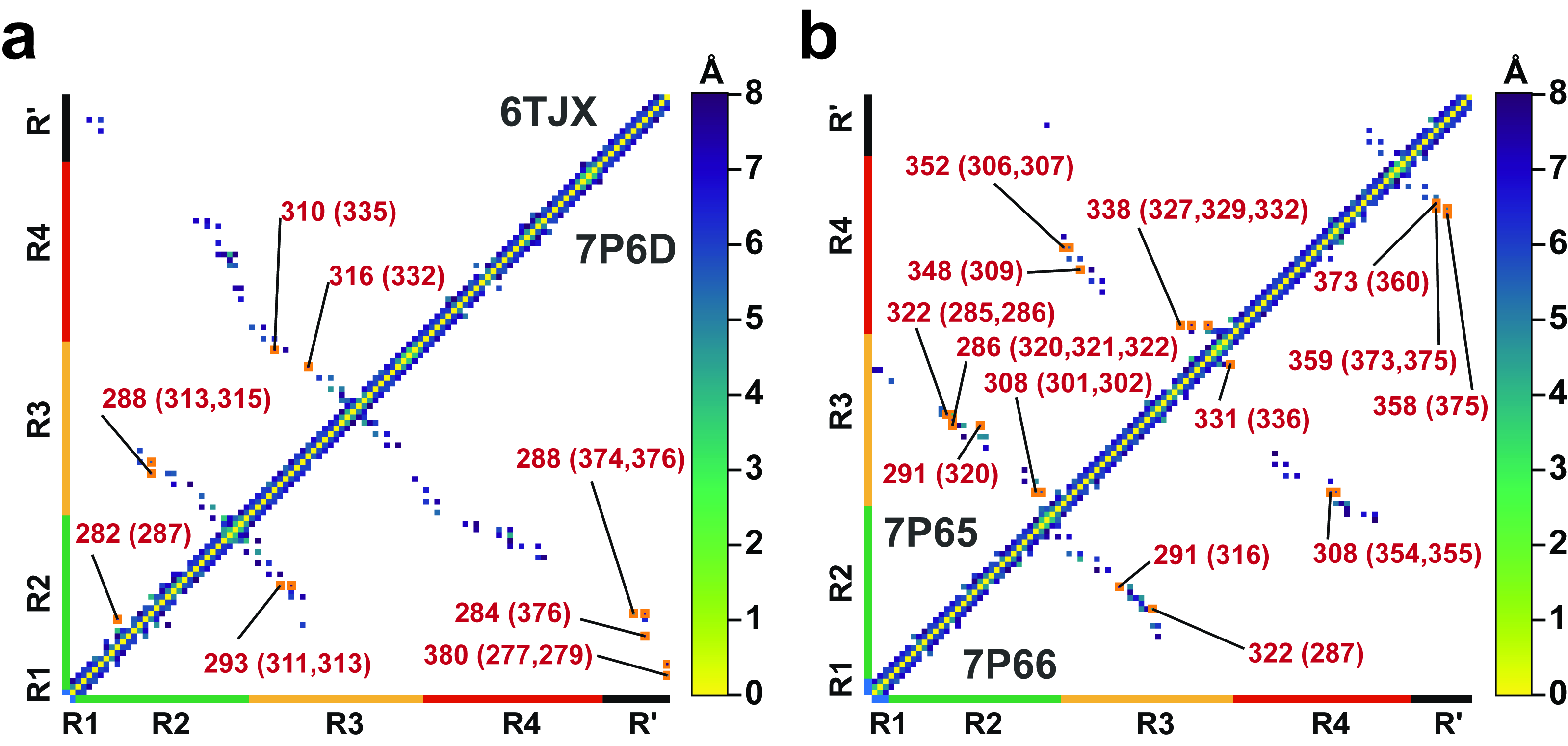

**Supplementary Figure 9. Contact maps comparing tauopathy fibril structures to illustrate differences in conformations and their contribution to stability.** a. contact-map comparison between CBD (PDB ID 6TJX) and AGD (PDB ID 7P6D) tau fibril templates. b. contact-map comparison between PSP (PDB ID 7P65) and GGT (PDB ID 7P66) tau fibril templates. Axes correspond to tau repeat-domain residue positions, with repeat regions indicated along the margins. The diagonal represents local sequence-neighbor contacts, whereas off-diagonal signals identify long-range residue-residue interactions within the fibril core. Colored pixels indicate contact strength or contact-rank score, with brighter colors marking higher-priority differential contacts. Labeled residues indicate positions where predicted contacts differ between the compared structures; residues outside parentheses correspond to one template and residues in parentheses correspond to the paired template. These analyses identify fold-specific interaction networks across R2, R3, R4 and R′ that may contribute to selective stabilization of CBD-, AGD-, PSP- and GGT-associated tau conformations.

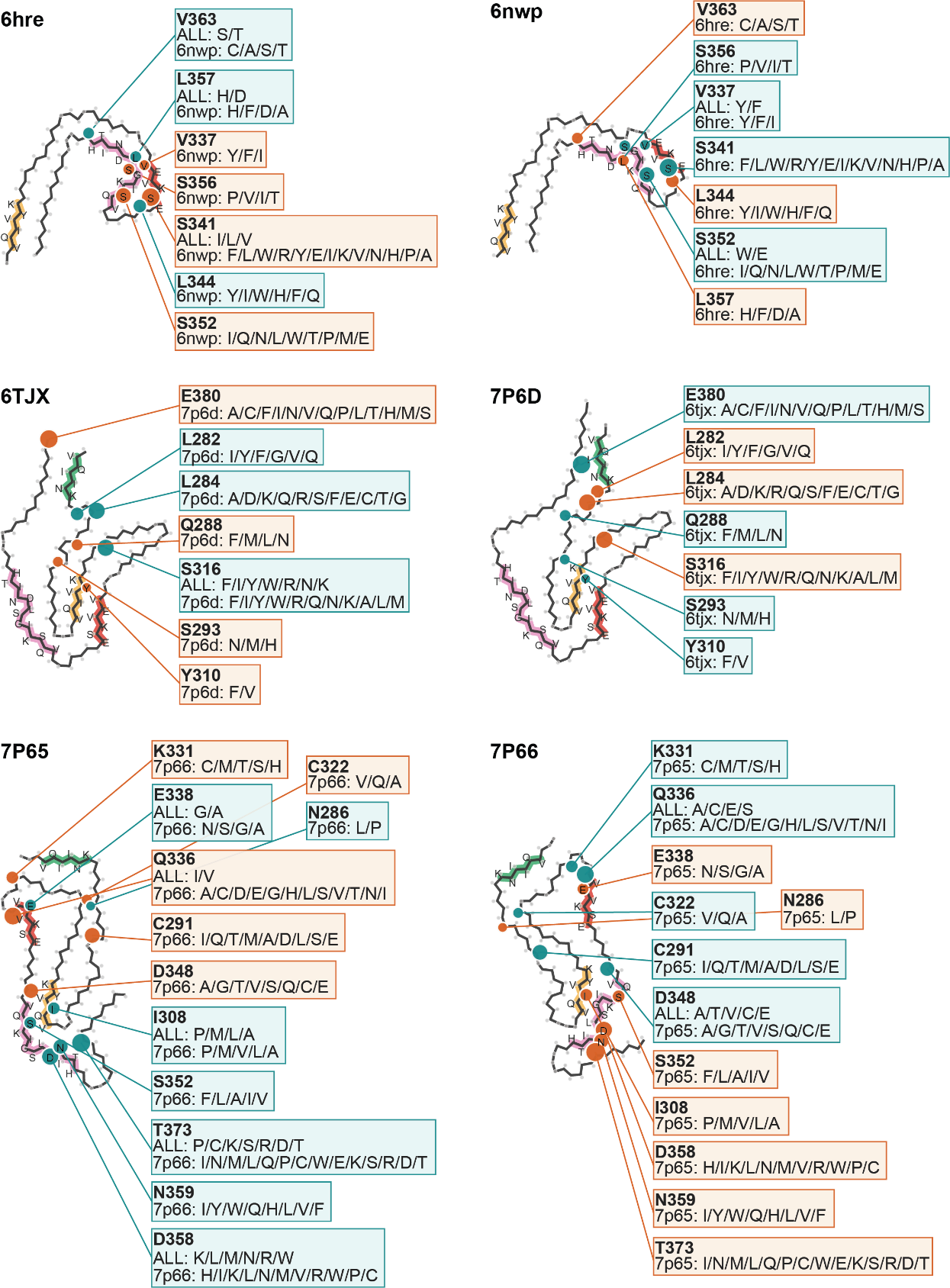

**Supplementary Figure 10. Mapping fold-discriminating residues and tolerated substitutions onto disease-associated tau fibril structures.** Representative tau fibril cores are shown for AD (PDB ID 6HRE), CTE (PDB ID 6NWP), CBD (PDB ID 6TJX), AGD (PDB ID 7P6D), PSP (PDB ID 7P65) and GGT (PDB ID 7P66). Highlighted residues indicate positions identified by saturation mutagenesis and contact-map analyses as candidate determinants of fold compatibility. Callout boxes list amino acid substitutions predicted to be tolerated or favored at each position. For each residue, substitutions labeled “ALL” indicate amino acids mutations that yield selectivity of this structure related to all five other structures, whereas a structure-specific labels indicate substitutions predicted to be selective relative to the indicated. Cyan and orange highlights distinguish substitutions associated with the indicated fold that stabilize or destabilize that conformation, respectively. For AD and CTE, residues including V337, S341, L344, S352, S356, L357 and V363 cluster near the R3/R4 interface and define local packing differences between the two folds. For CBD and AGD, residues including L282, L284, Q288, S293, Y310, S316 and E380 highlight distinct interaction networks spanning R2, R3 and R4. For PSP and GGT, residues including N286, C291, I308, C322, K331, Q336, E338, D348, S352, D358, N359 and T373 define broader fold-specific compatibility networks across the ordered tau core. Together, these maps show that tau fold compatibility is encoded by distributed residue networks rather than by a single amyloid motif and they identify candidate substitutions for testing strain-selective stabilization or destabilization.

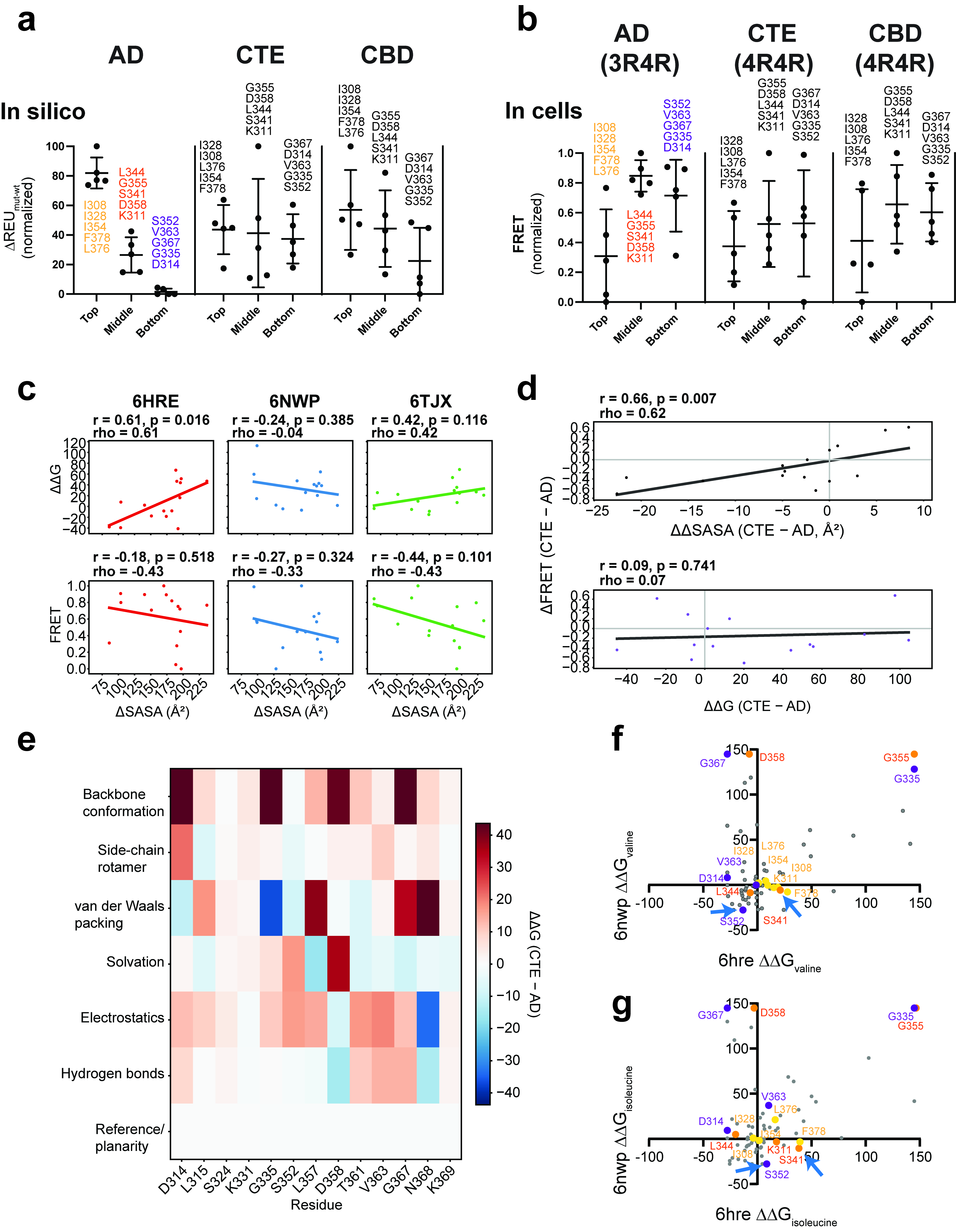

**Supplementary Figure 11. Energetic and cellular profiling identifies fold-dependent packing constraints in tau fibrils. a.** Predicted energetic effects of alanine substitutions in the AD (PDB ID 6HRE), CTE (PDB ID 6NWP) and CBD (PDB ID 6TJX) fibril folds. Rosetta energy differences between each alanine mutant and the corresponding WT sequence were min-max normalized within each fold. Residues were grouped according to their predicted effects in the AD structure: high destabilization (“top,” gold), intermediate or mixed effects (“middle,” orange) and low destabilization or greater tolerance (“bottom,” purple). The same AD-defined residue groups are shown for CTE and CBD. Points represent individual positions; horizontal bars and error bars indicate the group mean ± SD. **b.** Normalized FRET incorporation of the same alanine substitutions into AD-derived inclusions in 3R/4R cells, CTE-derived inclusions in 4R/4R cells and CBD-derived inclusions in 4R/4R cells. Mutants are arranged according to the AD-defined groups in panel a. Points represent individual positions; horizontal bars and error bars indicate the group mean ± SD. **c.** Relationships between residue burial and predicted or experimentally measured substitution effects. Top, correlations between ΔSASA and predicted alanine-substitution ΔΔG for AD, CTE and CBD fibril structures. Bottom, correlations between ΔSASA and normalized FRET incorporation into the corresponding disease-derived inclusions. ΔSASA was calculated as the solvent-accessible surface area of the residue in the isolated monomer minus that in the modeled fibril; positive values therefore indicate burial upon fibril assembly. Lines show linear regressions. Pearson r with a two-sided p value and Spearman ρ are shown. **d.** Relationships between position-resolved differences in cellular incorporation and structural or energetic differences between CTE and AD. Top, Position-wise CTE - AD differences in FRET compared with CTE - AD differences in ΔSASA (top) or ΔΔG (bottom). Lines show linear regressions; Lines show linear regressions. Pearson r with a two-sided p value and Spearman ρ are shown. **e.** Rosetta score-term decomposition of the CTE–AD difference in alanine-substitution ΔΔG at selected positions. Contributions from backbone conformation, side-chain rotamers, van der Waals packing, solvation, electrostatics, hydrogen bonding and amino-acid reference or polarity terms are shown. Positive values indicate a larger energetic penalty in CTE than in AD, whereas negative values indicate a larger penalty in AD. **f,g.** Comparison of predicted substitution effects in the AD and CTE folds for valine (**f**) and isoleucine (**g**). Colored residues correspond to the AD-defined high-, intermediate- and low-destabilization groups used in panels a and b; gray points represent the remaining positions. Blue arrows indicate the S341 and S352 region selected for experimental design. Positions displaced from the diagonal relationship exhibit fold-biased responses to increased side-chain size and hydrophobicity.

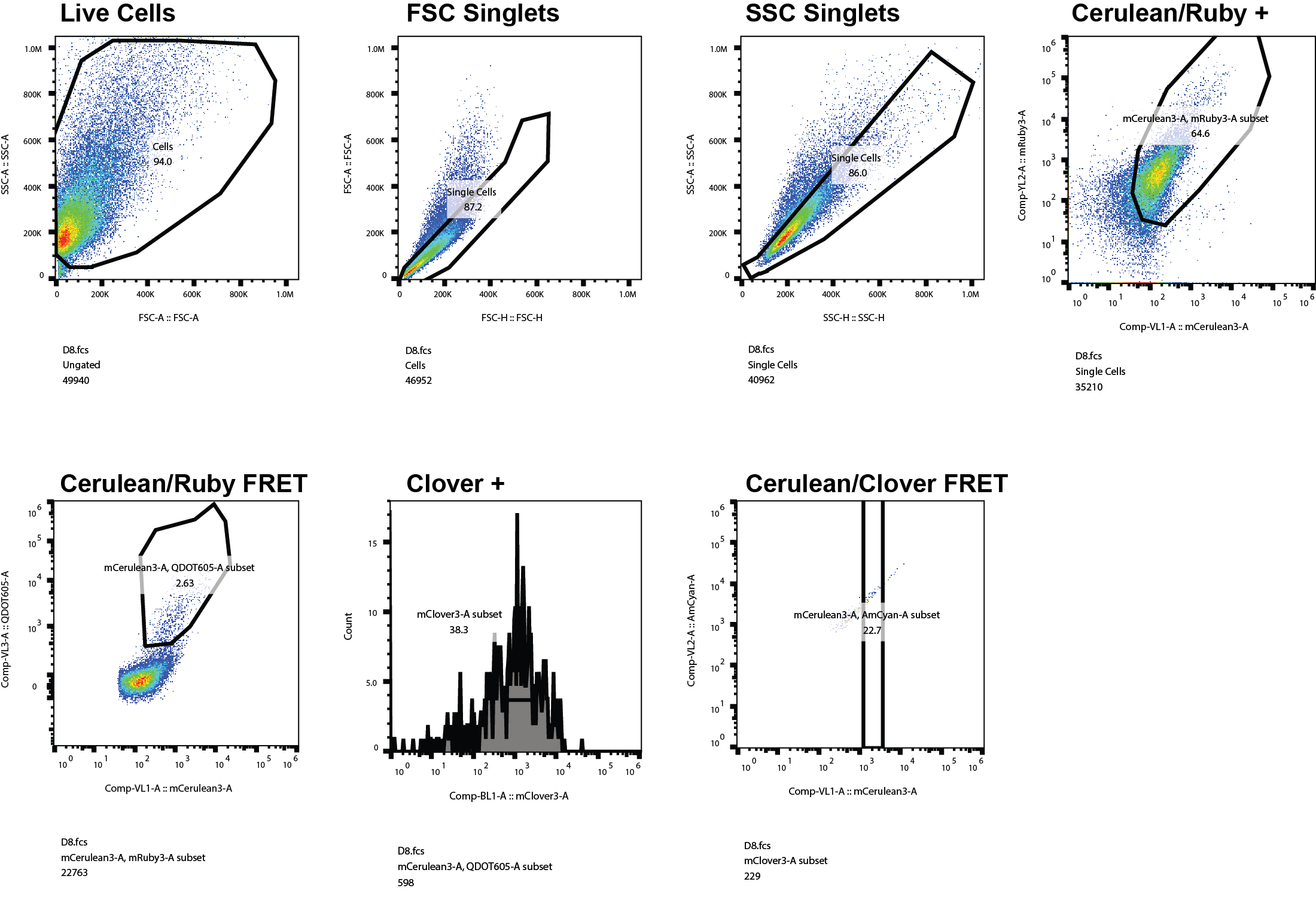

**Supplementary Figure 12. Gating strategy for alanine mutation incorporation assay in 4R/4R tauRD Cerulean/Ruby biosensors seeded with various tauopathy seeds.** Gating is sequentially done on live, single, mCerulean3 and mRuby channel double-positive cells, FRET quantification between mCerulean3/QDOT605-A (Cerulean/Ruby), expression level of mClover3-A positive cells and finally FRET between mCerulean3-A/AmCyan-A (Cerulean/Clover).

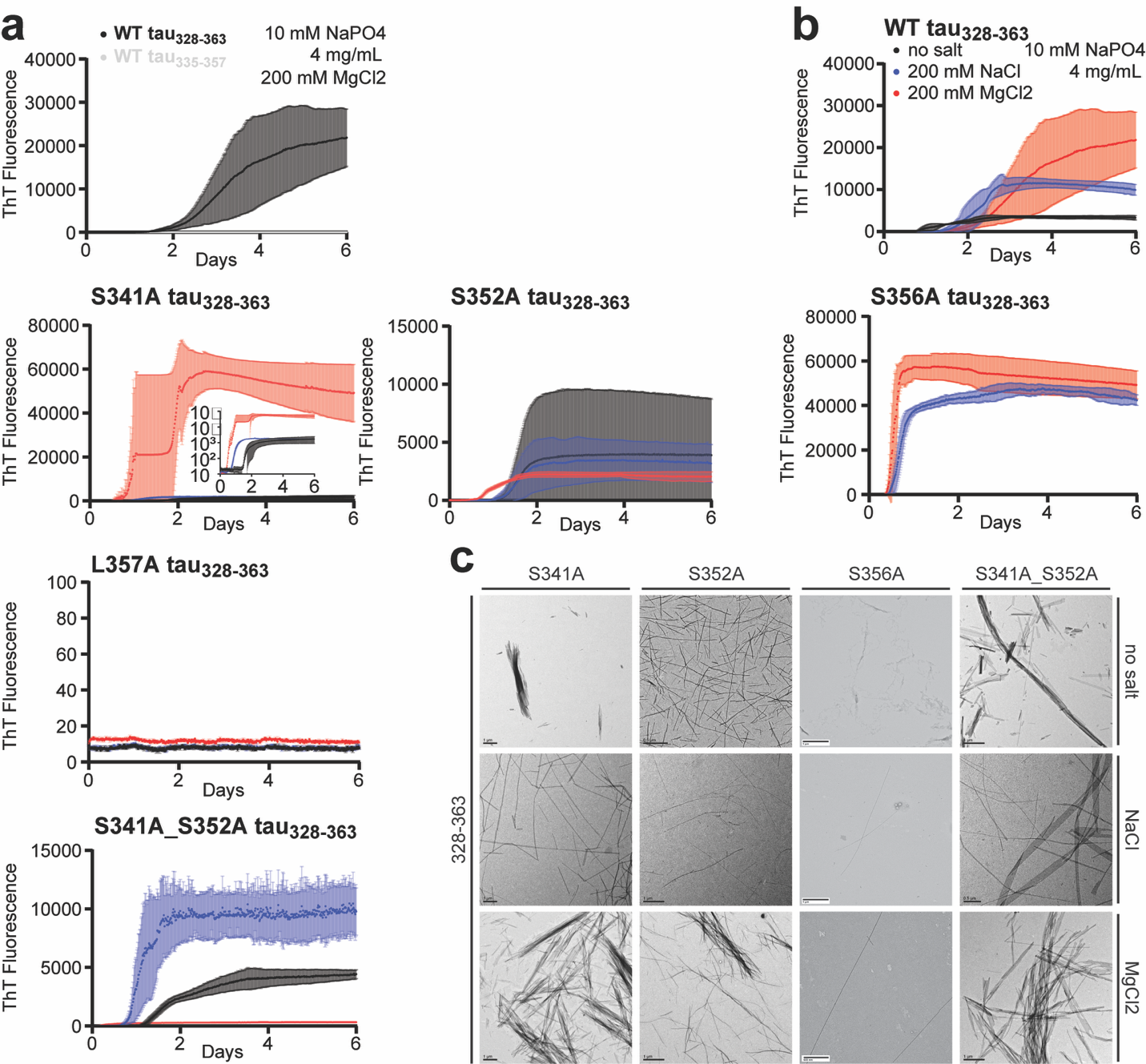

**Supplementary Figure 13. Alanine substitutions in tau328–363 differentially alter aggregation kinetics and fibril formation.** a. ThT aggregation kinetics comparing WT tau_328-363_ and WT tau_335-357_ in 10 mM sodium phosphate **with** 200 mM MgCl_2_. While the 328-363 construct showed an increase in ThT fluorescence over time, the 335-357 construct had no increase in ThT fluorescence. **b.** ThT aggregation kinetics for WT tau_328-363_ and alanine-substituted variants. Peptides were incubated in 10 mM sodium phosphate without added salt, with 200 mM NaCl, or with 200 mM MgCl_2_ and ThT fluorescence was monitored over time. WT tau_328-363_ aggregated under all conditions, with the strongest signal in MgCl_2_. S341A, S352A and S356A altered the rate and amplitude of aggregation in a buffer-dependent manner, whereas L357A showed little to no detectable ThT fluorescence under any condition. The S341A/S352A double mutant retained aggregation and showed enhanced aggregation in NaCl relative to the no-salt and MgCl_2_ conditions. Shaded regions indicate variation among replicate measurements. c. Representative negative-stain electron microscopy images of tau_328-363_ alanine variants after aggregation reactions. ThT-positive samples generally contained fibrillar assemblies, whereas conditions with low or absent ThT signal showed reduced fibril formation or sparse aggregates. Together, these data indicate that residues within the 328-363 region make distinct contributions to amyloid formation, with L357 emerging as a critical determinant of aggregation and S341, S352 and S356 modulating aggregation in a buffer-dependent manner.

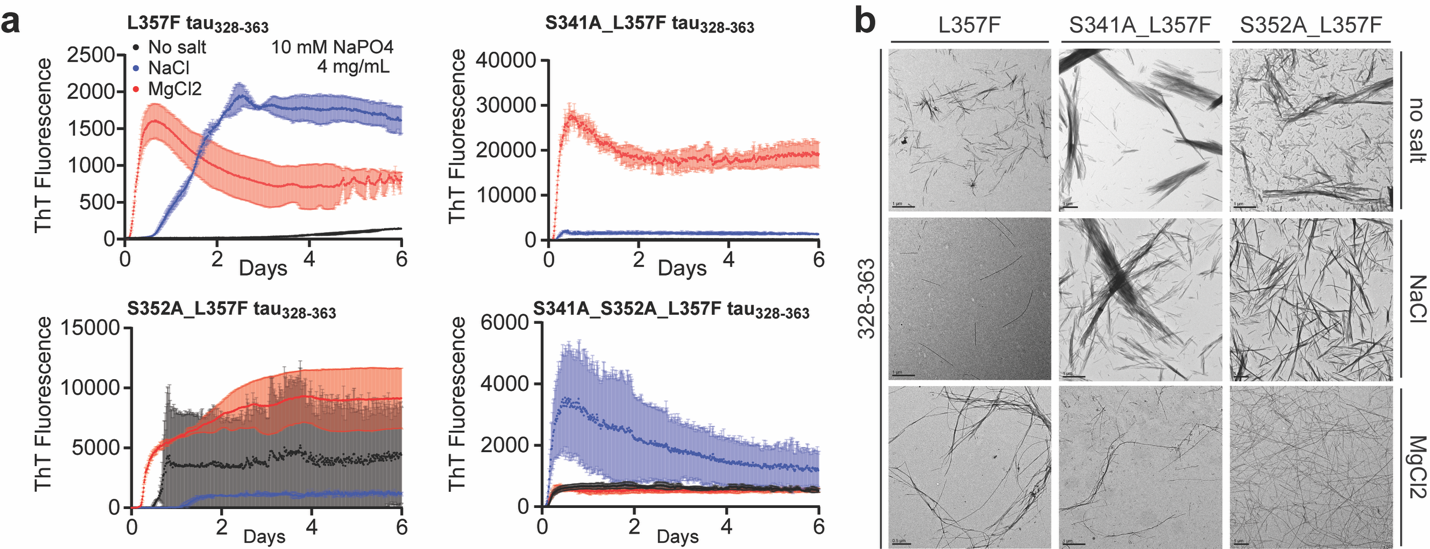

**Supplementary Figure 14. AD-stabilizing tau_328-363_ substitutions alter aggregation kinetics and fibril formation in a buffer-dependent manner.** **a.** Thioflavin T (ThT) aggregation kinetics for tau_328-363_ variants containing L357F alone or in combination with S341A and/or S352A. Peptides were incubated in 10 mM sodium phosphate without added salt, with 200 mM NaCl, or with 200 mM MgCl₂ and ThT fluorescence was monitored over time. L357F restored aggregation compared with the aggregation-deficient L357A variant, but showed strong buffer dependence, with more rapid aggregation in salt-containing conditions. Combining L357F with S341A or S352A further altered aggregation kinetics and fluorescence amplitude. The S341A/S357F and S352A/L357F variants showed rapid MgCl₂-dependent aggregation, whereas the S341A/S352A/L357F triple mutant showed strongest aggregation in NaCl and reduced signal in MgCl₂. Shaded regions indicate variation among replicate measurements. b. Representative negative-stain electron microscopy images of L357F-containing tau_328-363_ variants after aggregation reactions. ThT-positive reactions contained fibrillar assemblies with variable abundance and morphology across variants and buffer conditions. Together, these data indicate that L357 is a key determinant of tau_328-363_ assembly and that an aromatic residue at this position can support fibril formation, with aggregation outcome further tuned by substitutions at S341 and S352.

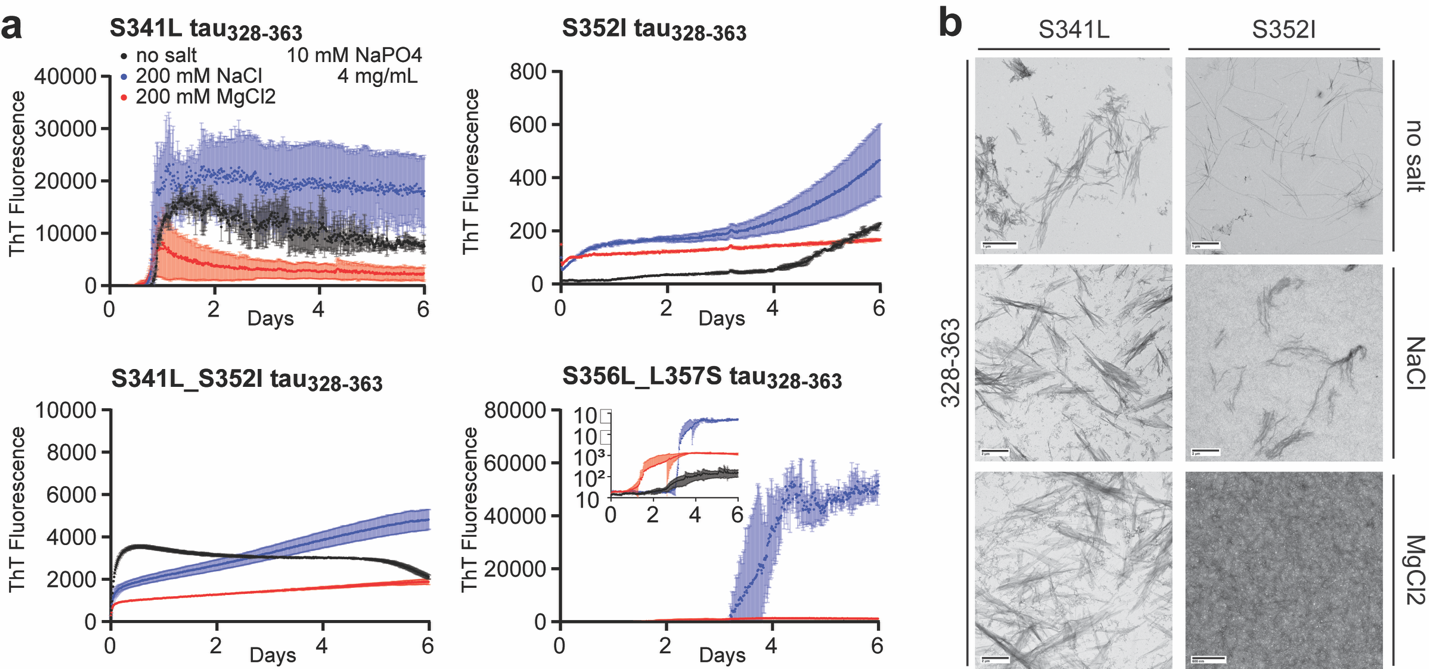

**Supplementary Figure 15. CTE-biased tau328–363 substitutions alter aggregation kinetics and endpoint fibril formation.** a. ThT aggregation kinetics for S341L, S352I, S341L/S352I and S356L/L357S in 10 mM sodium phosphate without added salt, with 200 mM NaCl, or with 200 mM MgCl₂. Curves show mean ± SD for three replicate wells. b. Endpoint negative-stain TEM images collected after the six-day aggregation reactions. Scale bars are shown in each micrograph. The data show that fold-biased substitutions alter packing compatibility and aggregation in a sequence- and buffer-dependent manner.

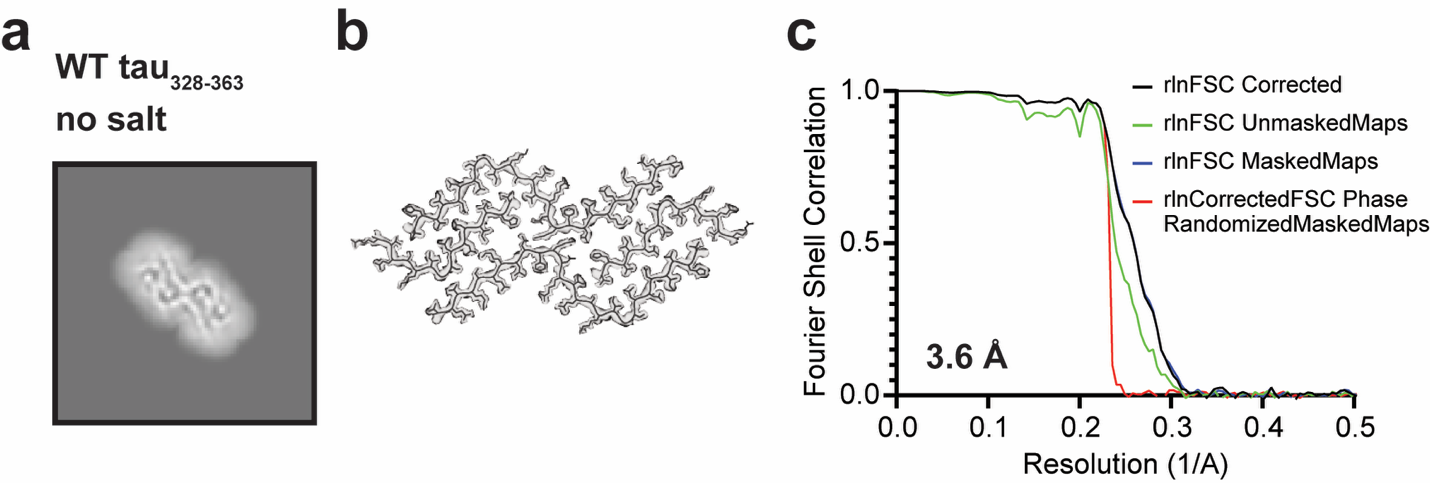

**Supplementary Figure 16. Cryo-EM data processing for WT tau_328-363_ aggregates.** a. Representative cryo-EM analysis of tau_328-363_ fibrils. Left, two-dimensional class average showing the fibril cross-section. b. reconstructed cryo-EM density map with fitted atomic model of the tau_328-363_ amyloid core. c. Fourier shell correlation curves for the unmasked map, masked map, phase-randomized masked map and corrected FSC analysis. The final reconstruction reached an estimated resolution of approximately 3.46 Å, post refinement.

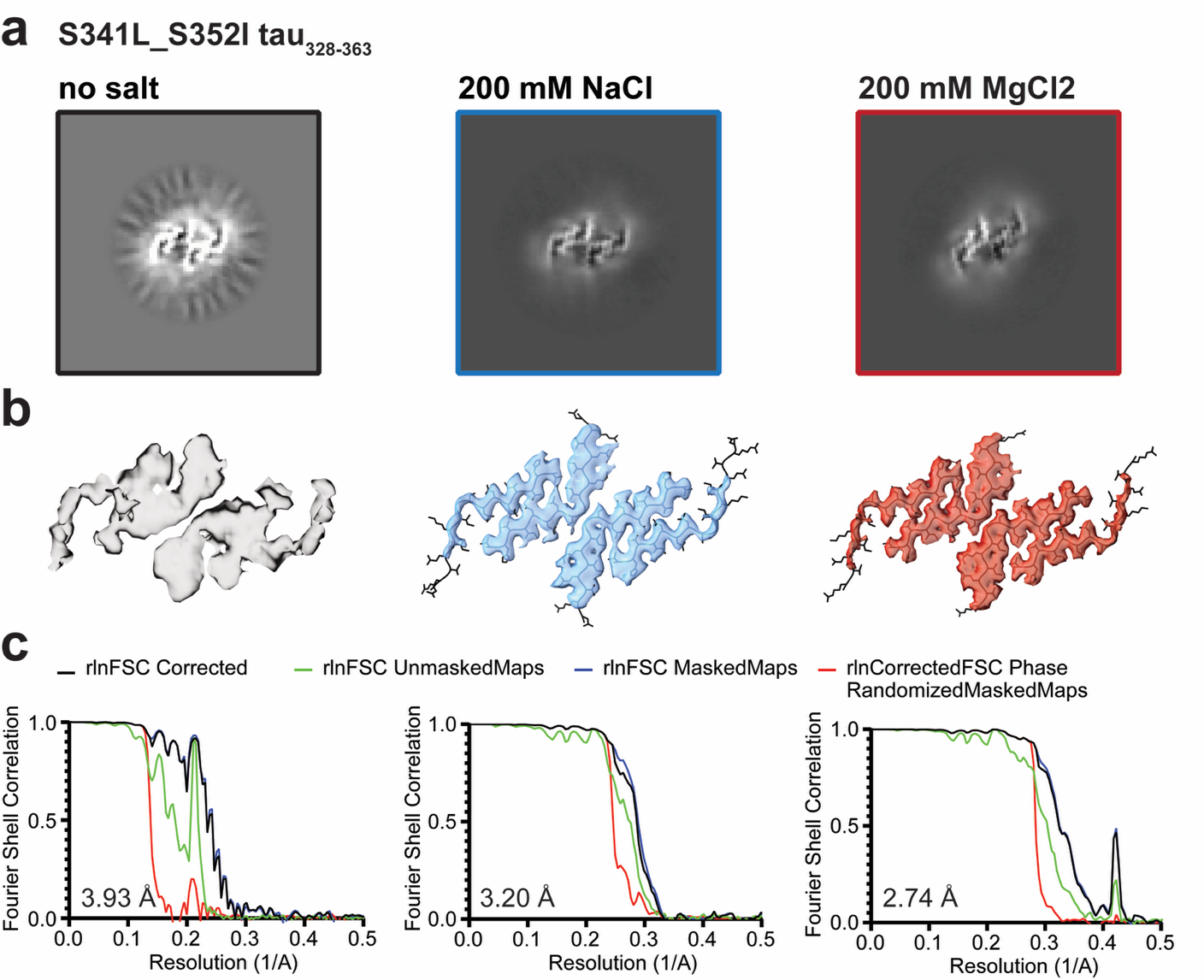

**Supplementary Figure 17. Cryo-EM reconstructions of S341L/S352I tau328–363 fibrils formed under different ionic conditions.** **a.** Representative two-dimensional class averages of tau328–363 fibrils formed in 10 mM sodium phosphate without added salt **(left)**, with 200 mM NaCl **(middle)**, or with 200 mM MgCl_2_ **(right)**. The class averages show ordered fibril cross-sections under each condition. **b.** Corresponding cryo-EM density maps and fitted atomic models for fibrils formed without added salt **(left)**, with 200 mM NaCl **(middle)**, or with 200 mM MgCl_2_ **(right)**. The reconstructions indicate that tau_328-363_ forms ordered amyloid assemblies under all three buffer conditions, with condition-dependent differences in fibril packing and map quality. **c.** Fourier shell correlation curves for the no-salt **(left)**, NaCl **(middle)** and MgCl_2_ **(right)** reconstructions. Curves are shown for the unmasked map, masked map, phase-randomized masked map and corrected FSC analysis. The final reconstruction for the MgCl_2_ conditions reached an estimated resolution of approximately 2.2 Å, post refinement.

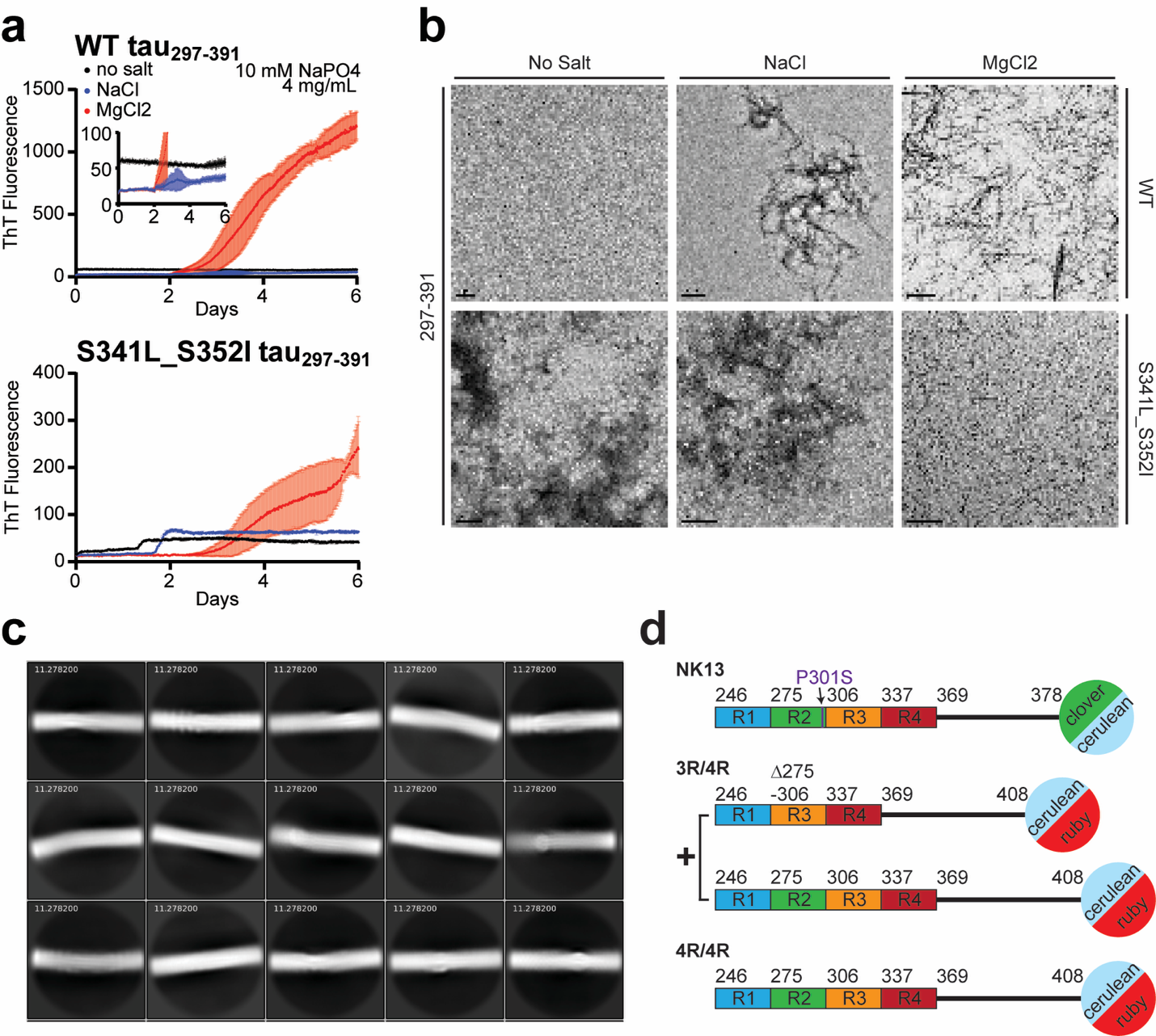

**Supplementary Figure 18. S341L/S352I alters tau297–391 aggregation and is compatible with cellular reporter expression.** a. ThT aggregation kinetics for WT and S341L/S352I tau297–391 in no-salt, 200 mM NaCl, or 200 mM MgCl₂ conditions. Curves show mean ± SD for three replicate wells. b. Endpoint negative-stain TEM images collected after the six-day aggregation assay; scale bars are shown in each image. c. Representative cryo-EM 2D class averages of S341L/S352I tau_297-391_ fibrils. d. Reporter constructs: NK13 contains P301S tauRD_246-378_ fused separately to mClover3 or mCerulean3, whereas 3R/4R and 4R/4R reporters contain tauRD_246-408_ fusions to mCerulean3 or mRuby3.

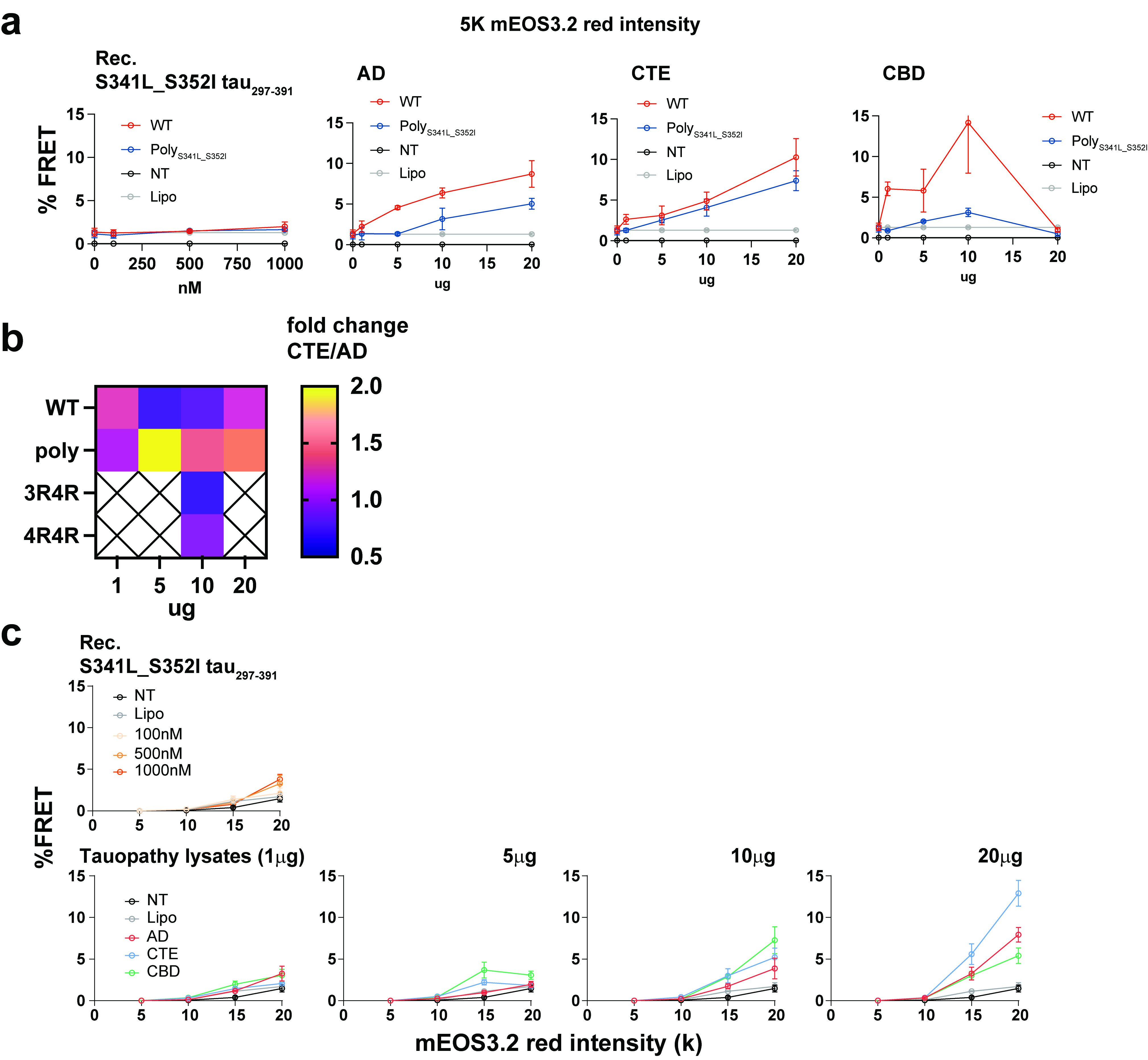

**Supplementary Figure 19. S341L/S352I tauRD-mEOS3.2 biosensors show reduced sensitivity but enhanced selectivity for CTE-derived seeds.** **a.** Seeding responses of WT and S341L/S352I tauR*D*_246-408_-mEOS3.2 polyclonal biosensor populations analyzed within the 5,000-arbitrary-unit mEOS3.2 red-intensity gate. Left, cells treated with increasing concentrations of recombinant S341L/S352I tau_297-391_ aggregates*.* Right, cells treated with increasing amounts of AD-, CTE-, or CBD-derived brain lysate. No treatment (NT) and Lipofectamine-only (Lipo) cells served as controls. Compared with WT, the S341L/S352I polyclonal reporter generally produced lower absolute FRET responses but showed a stronger relative preference for CTE- over AD-derived lysate. **b.** Heat map showing the fold difference between CTE- and AD-induced FRET in WT and S341L/S352I polyclonal mEOS3.2 reporters across the indicated lysate amounts. Values for the 3R/4R and 4R/4R mCerulean3/mRuby3 biosensors are included at 10 μg for comparison; crossed cells indicate conditions that were not measured. Values greater than 1 indicate preferential seeding by CTE-derived lysate. **c.** Expression-dependent seeding of the high-expressing S341L/S352I single-cell clone 4. Top, responses to 100, 500, or 1,000 nM recombinant S341L/S352I tau_297-391_ aggregates. Bottom, responses to 1, 5, 10, or 20 μg of AD-, CTE-, or CBD-derived brain lysate. FRET-positive cells were quantified within mEOS3.2 red-intensity gates centered at 5k, 10k, 15k and 20k arbitrary units. NT and Lipo cells served as controls. Data represent three technical replicates and are shown as mean ± SEM.

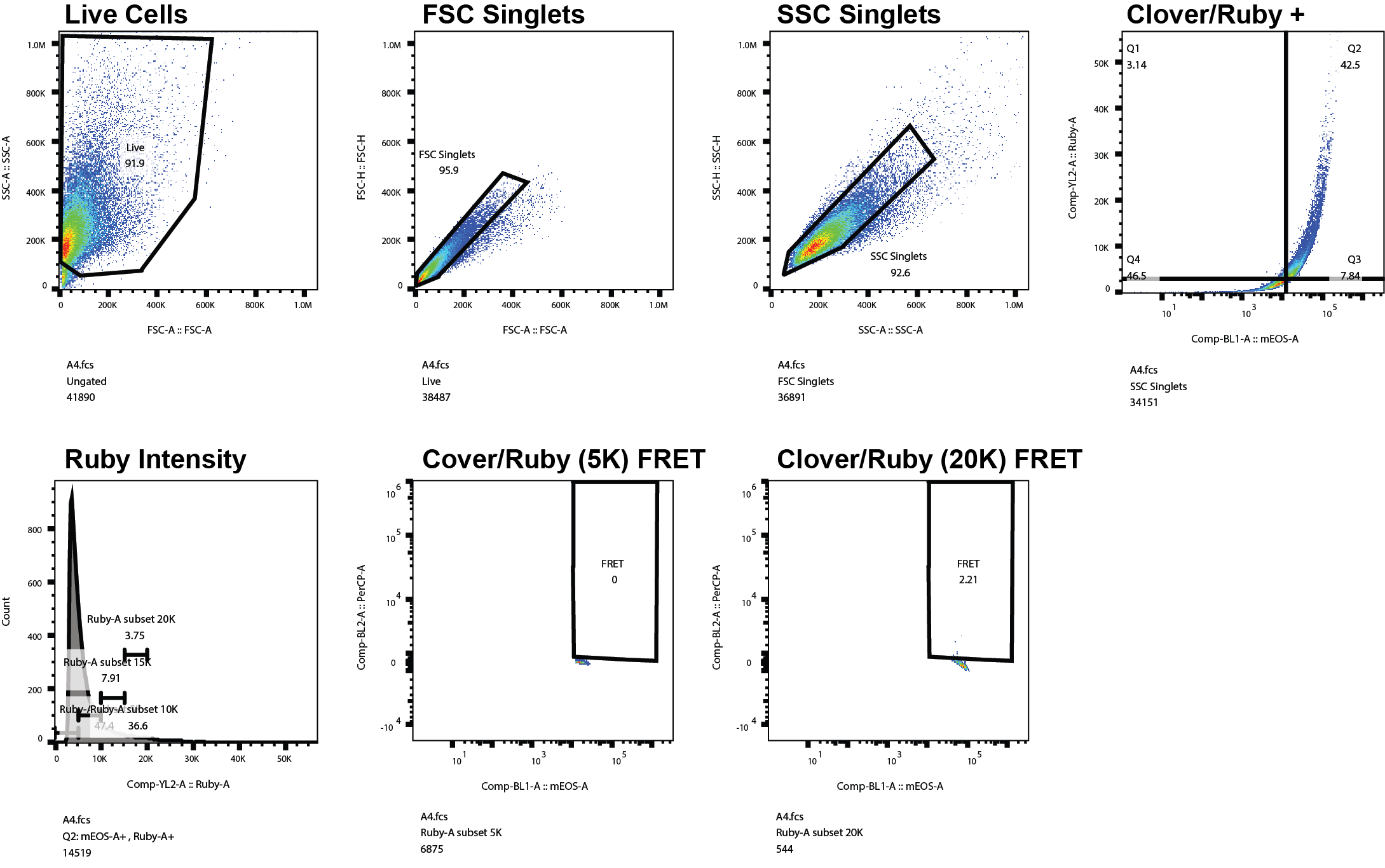

**Supplementary Figure 20. Gating strategy for seeding assays in clone4 S341L/S352I tauRD mEOS3.2 biosensors seeded with various tauopathy seeds.** Gating is sequentially done on live, single, mEOS-A and Ruby-A channel double-positive cells, expression level of Ruby-A positive cells and finally FRET between mEOS-A/PerCP-A (Clover/Ruby).

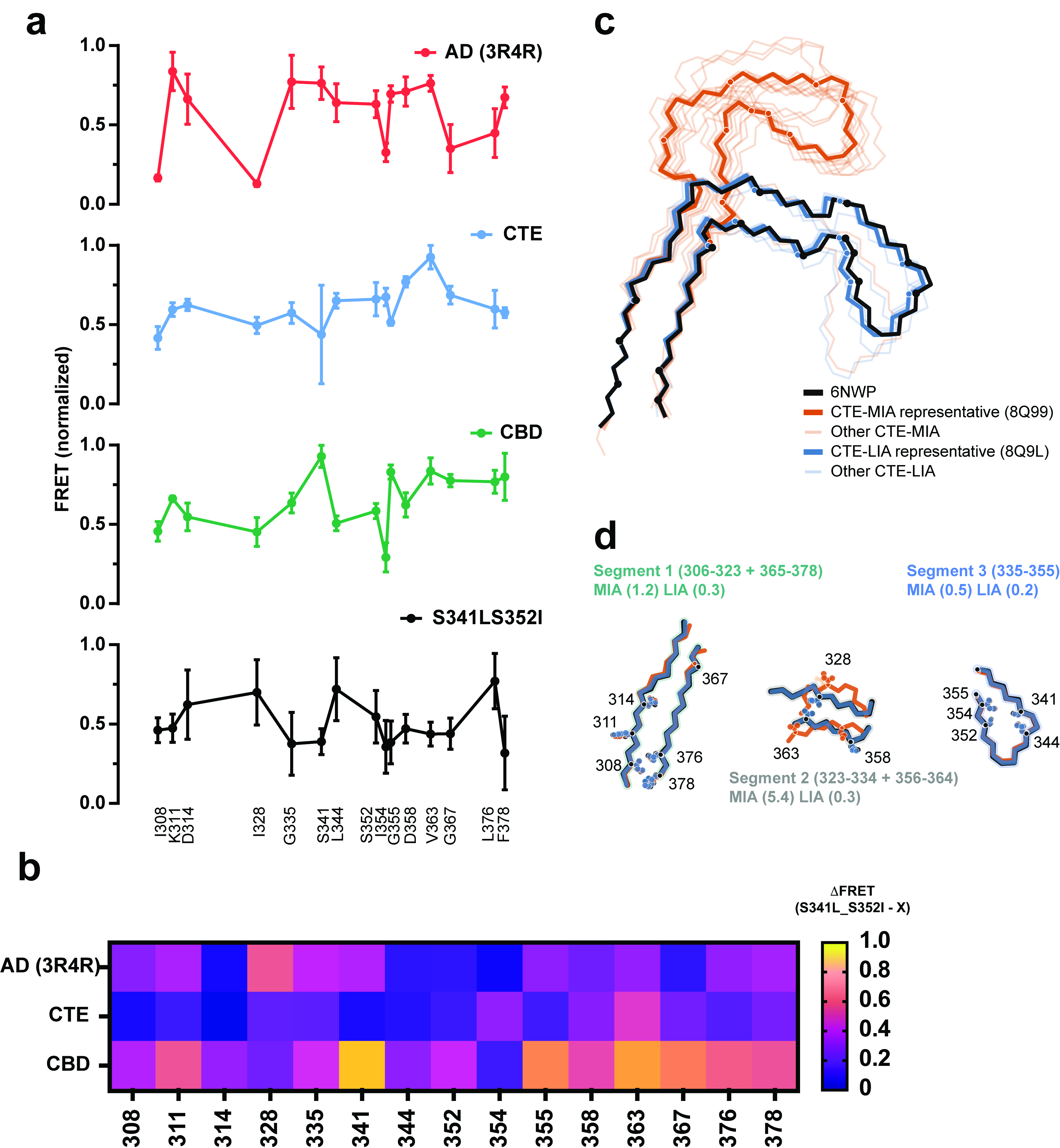

**Supplementary Figure 21. S341L/S352I-seeded inclusions exhibit a CTE-like alanine-incorporation profile.** **a.** Normalized FRET incorporation profiles for 15 tau alanine mutants in cells containing AD-, CTE-, CBD-, or recombinant S341L/S352I *tau*_297-391_-seeded inclusions. FRET values were min-max normalized within each seed dataset. Points and error bars represent mean ± SD from three replicate wells. b. Heat map showing the absolute position-wise differences between the S341L/S352I incorporation profile and the AD, CTE, or CBD reference profile. Position-wise ΔFRET was calculated as normalized FRET(S341L/S352I-seeded) – normalized FRET (reference-seeded) and their absolute magnitudes are displayed. Lower values indicate greater similarity to the S341L/S352I profile. Across all 15 positions, the mean absolute ΔFRET was 0.181 for CTE, compared with 0.258 for AD and 0.499 for CBD. **c.** Structural comparison of the mature CTE protofilament (PDB 6NWP, black) with representative middle intermediate amyloid (MIA; PDB ID 8Q99, orange) and late intermediate amyloid (LIA; PDB ID 8Q9L, blue) conformations. Additional MIA and LIA structures are shown in lighter shades. **d.** Segment-resolved alignments of the mature CTE, MIA and LIA structures. Segment 1 comprises residues 306-323 and 365-378; segment 2 comprises residues 323-334 and 356-364; and segment 3 comprises residues 335-355. Values in parentheses report the RMSD, in Å, of each representative intermediate relative to the corresponding segment of the mature CTE structure. Segment 2 exhibits substantially greater structural divergence in the MIA state than the other segments, identifying the middle region of the fold as the principal site of intermediate-state variability
